# A multiple-trait Bayesian Lasso for genome-enabled analysis and prediction of complex traits

**DOI:** 10.1101/852749

**Authors:** Daniel Gianola, Rohan L. Fernando

## Abstract

A multiple-trait Bayesian LASSO (MBL) for genome-based analysis and prediction of quantitative traits is presented and applied to two real data sets. The data-generating model is a multivariate linear Bayesian regression on possibly a huge number of molecular markers, and with a Gaussian residual distribution posed. Each (one per marker) of the *T* × 1 vectors of regression coefficients (*T* : number of traits) is assigned the same *T* –variate Laplace prior distribution, with a null mean vector and unknown scale matrix **Σ**. The multivariate prior reduces to that of the standard univariate Bayesian LASSO when *T* = 1. The covariance matrix of the residual distribution is assigned a multivariate Jeffreys prior and **Σ** is given an inverse-Wishart prior. The unknown quantities in the model are learned using a Markov chain Monte Carlo sampling scheme constructed using a scale-mixture of normal distributions representation. MBL is demonstrated in a bivariate context employing two publicly available data sets using a bivariate genomic best linear unbiased prediction model (GBLUP) for benchmarking results. The first data set is one where wheat grain yields in two different environments are treated as distinct traits. The second data set comes from genotyped *Pinus* trees with each individual was measured for two traits, rust bin and gall volume. In MBL, the bivariate marker effects are shrunk differentially, i.e., “short” vectors are more strongly shrunk towards the origin than in GBLUP; conversely, “long” vectors are shrunk less. A predictive comparison was carried out as well where, in wheat, the comparators of MBL where bivariate GBLUP and bivariate Bayes C*π*, a variable selection procedure. A training-testing layout was used, with 100 random reconstructions of training and testing sets. For the wheat data, all methods produced similar predictions. In *Pinus*, MBL gave better predictions that either a Bayesian bivariate GBLUP or the single trait Bayesian LASSO. MBL has been implemented in the Julia language package JWAS and is now available for the scientific community to explore with different traits, species and environments. It is well known that there is no universally best prediction machine and MBL represents a new piece in the armamentarium for genome-enabled analysis and prediction of complex traits.

## 2 Introduction

Two main paradigms have been employed for investigating statistical associations between molecular markers and complex traits: marker-by-marker genome-wide association studies (GWAS) and whole-genome regression approaches (WGR). GWAS is dominant in human genetics; Visscher et al. (2017) present a perspective and Gianola et al. (2016) formulate a statistically orientated critique. WGR was developed mostly in animal and plant breeding (e.g., Lande and Thompson 1990; Meuwissen et al. 2001; Gianola et al. 2003) primarily for predicting future performance, but it has received some attention in human genetics as well (e.g., Lee et al. 2011; Yang et al. 2010; de los Campos et al. 2011; Makowsky et al. 2011; López de Maturana et al. 2014). de los Campos et al. (2013), Gianola (2013) and Isik et al. (2017) reviewed an extensive collection of WGR approaches. Other studies noted that WGR can be used both for “discovery” of associations and for prediction (Moser et al. 2015; Goddard et al. 2016; Fernando et al. 2017). Hence, WGR methodology is an active area of research.

Multiple-trait analysis has been of great interest in plant and animal breeding for a long-time, mainly from the point of view of joint selection for many traits (Smith 1936; Hazel 1943; Walsh and Lynch 2018). Henderson and Quaas (1976) developed multi-trait best linear unbiased prediction of breeding values for all individuals and traits measured in a population of animals, a methodology that gradually became routine in the field. For example, Gao et al. (2018), described an application of a 9-variate model to data representing close to seven million and four million Holstein and Nordic Red cattle, respectively; the nine traits were milk, fat and protein yields in each of the first three lactations of the cows.

A multiple-trait analysis is also a natural choice in quests for understanding and dissecting genetic correlations between traits using molecular markers, e.g., evaluating whether pleiotropy or linkage disequilibrium are at the roots of between-trait associations (Gianola et al. 2015; Cheng et al. 2018a). For instance, Galesloot et al. (2014) compared six methods of multivariate GWAS via simulation and found that all delivered a higher power than single-trait GWAS, even when genetic correlations were weak. Many single-trait WGR methods extend directly to the multiple-trait domain, e.g., genomic best linear unbiased prediction (GBLUP; Van Raden 2007). Other procedures such as Bayesian mixture models are more involved, but extensions are available (Calus and Veerkamp 2011; Jia and Jannink 2012; Cheng et al. 2018a). The mixture model of Cheng et al. (2018a) is particularly interesting because it provides insight into whether markers affect all, some or none of the traits addressed. For example, the proportion of markers in each of the (0, 0), (0, 1), (1, 0) and (1, 1) categories, where (0, 0) means “no effect” and (1, 1) denotes “effect” on each of two disease traits in *Pinus taeda* was estimated by Cheng et al. (2018a) using SNPs (single nucleotide polymorphisms). The proportion of MCMC samples falling into the (1, 1) class was less than 3%, with about 140 markers appearing as candidates for further scrutiny of pleiotropy; 97% of the SNP were in the (0, 0) class and 0.5% were in the (0, 1) and (1, 0) classes. It must be noted that Cheng et al. (2018a) used Bayesian model averaging, so posterior estimates of effects and of their uncertainties constitute averages over all possible configurations. The resulting “average model” is not truly sparse as Bayesian mixture models always assign some posterior probability to each of the possible configurations. An alternative to a mixture is to use a prior distribution that produces strong shrinkage towards the origin of “weak” vectors of marker effects; here, each marker has a vector with dimension equal to the number of traits.

The LASSO (least absolute shrinkage and selection operator) presented by Tibshirani (1996) is a single-response method based on minimizing a linear regression residual sum of squares subject to a constraint based on an *L*_1_ norm. It can produce a sparse model, i.e., if the linear regression model has *p* regression coefficients, the LASSO yields a smaller model (i.e., model selection) but with a complexity that cannot exceed *N*, the number of observations. Tibshirani (1996) noted that the LASSO solutions can also be obtained by calculating the mode of the conditional posterior distribution of the regression coefficients in a Bayesian model in which each coefficient is assigned the same conditional double exponential or Laplace prior. Using a ridge regression reformulation of LASSO, it can be seen (Tibshirani 1996; Gianola 2013) that its Bayesian version shrinks small-value regression coefficients very strongly towards zero, whereas large-effect variants are regularized to a much lesser extent than in ridge regression. Yuan and Lin (2006) and Yuan et al. (2007) considered the problem of clustering regression coefficients into groups (factors), with the focus becoming factor selection, as opposed to the predictor variable selection that takes place in LASSO. For instance, a cluster could consist of a group of markers in tight physical linkage. These authors noted that, in some instances, grouping enhances prediction performance over ridge regression, while in others, it does not. Such finding is consisting with knowledge accumulated in close to two decades of experience with genome-enabled prediction in animal breeding: there is no universally best prediction machine. A multiple-trait application of a LASSO penalty on regression coefficients was presented by Li et al. (2015). These authors assigned a multivariate Laplace distribution to the model residuals and a group-LASSO penalty (Yuan and Lin 2006) to the regression coefficients. The procedure differs from Tibshirani’s LASSO in that the model selects vectors of regressors (corresponding to regressions of a given marker over traits) as opposed to single-trait predictor variables.

Park and Casella (2008) introduced a fully Bayesian LASSO (BL). Contrary to LASSO, BL produces a model where all regression coefficients are non-null (even if *p* > *N*); most regressions are often tiny in value, except those associated with covariates (markers) with strong effects. In short, LASSO produces a sparse model whereas BL yields an effectively sparse specification, similar to Bayesian mixture models such as Bayes B (Meuwissen et al. 2001). The fist application of the BL in quantitative genetics was made by Yi and Xu (2008) in the context of quantitative trait locus (QTL) mapping, with subsequent applications in de los Campos et al. (2009), Legarra et al. (2011), Li et al. (2011) and Lehermeier et al. (2013).

It appears that a multiple-trait generalization of the BL has not been reported hereto. The present paper describes a multi-trait Bayesian LASSO (MBL) model based on adopting a multivariate Laplace distribution with unknown scale matrix as prior distribution for the markers or variants under scrutiny. The MBL is introduced and compared with a multiple-trait GBLUP (MTGBLUP) using wheat and pine tree data sets. Section “The multi-trait regression model” describes MBL, including a Markov chain Monte Carlo sampling algorithm. Subsequently, MBL is compared with MTGBLUP using a wheat data set. Finally, bivariate MBL and bivariate MTGBLUP are contrasted from a predictive perspective, showing a better performance of MBL over BLUP and over a single-trait Bayesian LASSO specification, corroborating the usefulness of multiple-trait analyses. The paper concludes with a general discussion and technical details are presented in Appendices.

### 3 The multi-trait regression model

Assume there are *T* traits observed in each of *N* individuals and let ***β***_*j*_ = {*β*_*jt*_} be a *T* × 1 vector of allelic substitution effects at marker *j* = 1, 2, …, *p*, with *β*_*jt*_ representing the effect of marker *j* on trait *t* (*t* = 1, 2, …,*T*). The multi-trait regression model (assuming no nuisance location effects other than a mean) for the *T* responses is

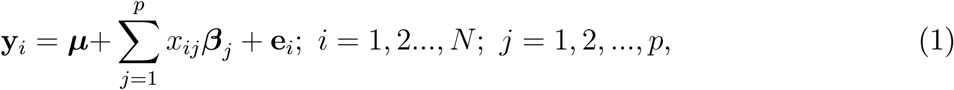

where y_*i*_ is a *T* × 1 vector of responses observed in individual *i*; ***µ*** = {*µ*_*t*_} is the vector of trait means and *x*_*ij*_ is the genotype individual *i* possesses at marker locus *j*. The residual vector **e**_***i***_ (*T* × 1) is assumed to follow the Gaussian distribution **e**_***i***_|**R**_0_ ∼ *N* (0, **R**_0_), where **R**_0_ is a *T* × *T* covariance matrix. All **e**_***i***_ vectors are assumed to be mutually independent and identically distributed.

If traits are sorted within individuals, the probability model associated with (1) can be represented as

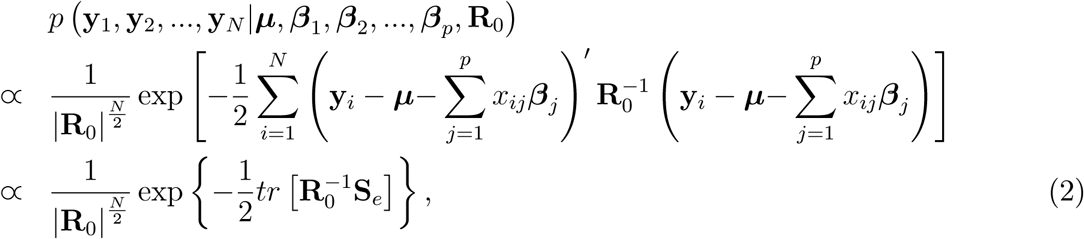

where

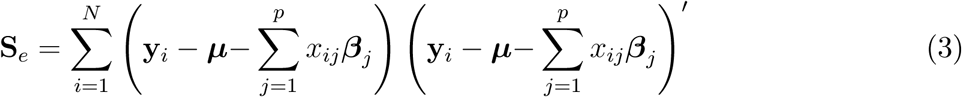

is a *T* × *T* matrix of sums of squares and products of the unobserved regression residuals.

The regression model can be formulated in an equivalent manner by sorting individuals within traits; we will use *T* = 3 hereinafter. Let 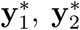 and 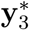 be response vectors of order *N* each observed for traits 1, 2, and 3, respectively. The representation of the model is

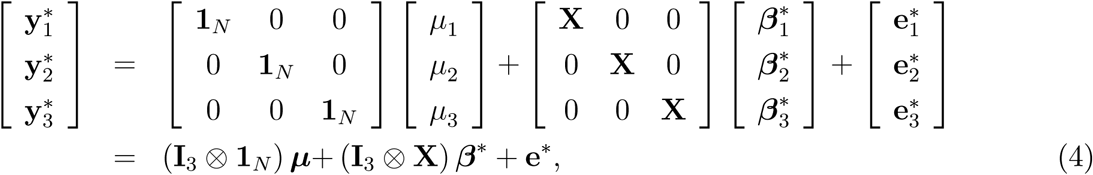

where **1**_*N*_ is an *N* × 1 vector of 1′s, **X** = {*x*_*ij*_} is an *N × p* matrix of marker genotypes, and 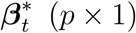 and 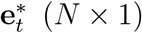 are vectors of regression coefficients and of residuals for trait *t*, respectively. Above, 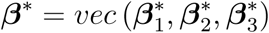 is a 3*p*×1 vector and 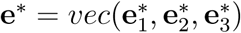 has dimension 3*N* × 1. Note that *Var* (**e**\*) = **R**_0_ ⊗ **I** = **R**. Putting 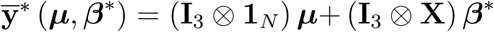, the probability model is

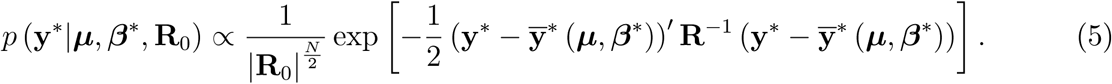

We will work with either (1) or (4), depending on the context.

#### 3.1 Bayesian prior assumptions

##### 3.1.1 Parameters *µ* and R_0_

The vector ***µ*** will be assigned a “flat” improper prior and Jeffreys non-informative prior (e.g., Sorensen and Gianola, 2002) will be adopted for **R**_0_ so that their joint prior density is

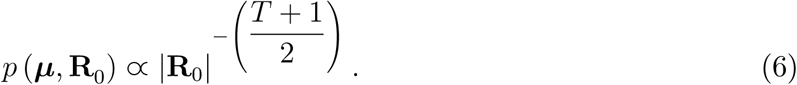

##### 3.1.2 Multivariate Laplace prior distribution (MLAP) for marker effects

The same *T*-variate Laplace prior distribution with a null mean vector will be assigned to each of the *T* × 1 vectors ***β***_*j*_ (*j* = 1, 2, …, *p*), assumed mutually independent, *a priori*. Gómez et al. (2007) presented a multi-dimensional version of the power exponential family of distributions; one special case is the multivariate Laplace distribution (MLAP). The density of the MLAP with a zero-mean vector used here is

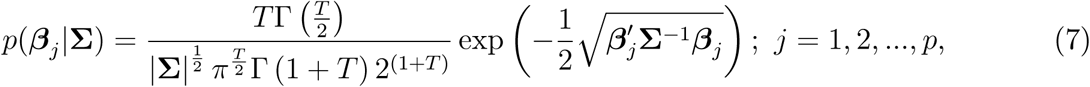

where **Σ** = {Σ_*tt*′_} is a *T* × *T* positive-definite scale matrix. The variance-covariance matrix of MLAP is

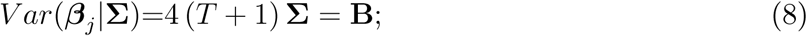

note that the absolute values of the elements of **B**, the inter-trait variance-covariance of marker effects, are larger than those of **Σ**. Hence, 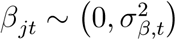 for ∀*j*, where 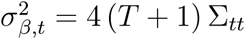 is the appropriate diagonal element of **B**; likewise, 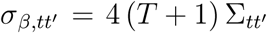 is the covariance of marker effects between traits *t* and *t*′, for all *j*. Putting *T* = 1 in (7) yields

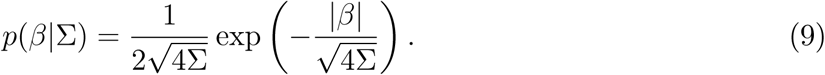

The preceding is the density of a double exponential (DE) distribution with null mean, parameter 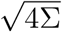 and variance *Var*(*β*) = 8Σ. As mentioned earlier, Tibshirani (1996) and Park and Casella (2008) used the DE distribution as conditional (given Σ) prior for regression coefficients in the BL, a member of the “Bayesian Alphabet” (Gianola et al. 2009). Gianola et al. (2018) assigned the DE distribution to residuals of a linear model for the purpose of attenuating outliers and Li et al. (2015) used the MLAP distribution for the residuals in a “robust” linear regression model for QTL mapping.

MLAP is therefore an interesting candidate prior for multi-trait marker effects in a multiple trait generalization of the Bayesian LASSO (MBL). A zero-mean MLAP distribution has a sharp peak at the 0 coordinates. Although when *T* = 1 MLAP reduces to a DE distribution, the marginal and conditional densities of MLAP are not DE. Gómez et al. (2007) showed that such densities are elliptically contoured, and thus not DE. Appendix A and Figures S1-S3 in the Supplemental material give background on MLAP.

Gómez-Sánchez-Manzano et al. (2008) showed that MLAP can be represented as a scaled mixture of normal distributions under the hierarchy: 1) 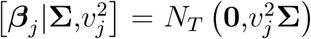, and 2) 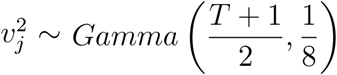 for *j* = 1, 2, …, *p*; 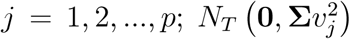 denotes a *T*–variate normal distribution with null mean and covariance matrix 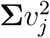. The density of 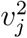 is

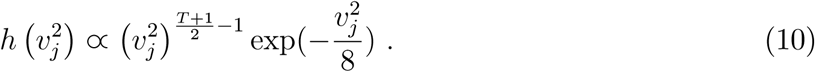

Let the collection of all marker effects over traits be represented by the *Tp* × 1 vector

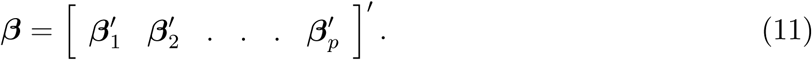

If independent and identical MLAP prior distributions are assigned to each of the sub-vectors, the joint prior density of all marker effects, given **Σ**, can be represented as

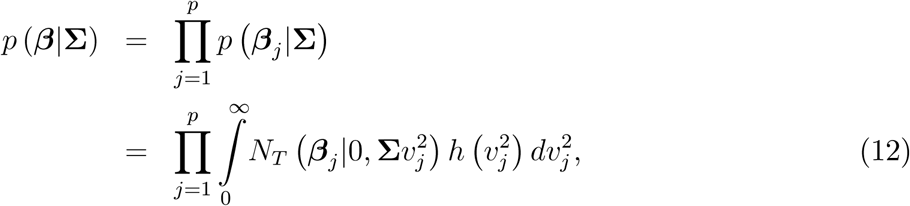

and the joint density of ***β*** and 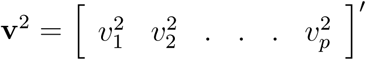 is

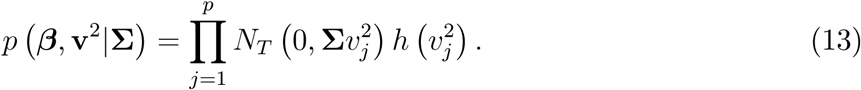

When individuals are sorted within traits (e.g., *T* = 3), note that [***β***\*|**Σ, v**^2^] is a *Tp*–dimensional normal distribution with null mean vector and covariance matrix

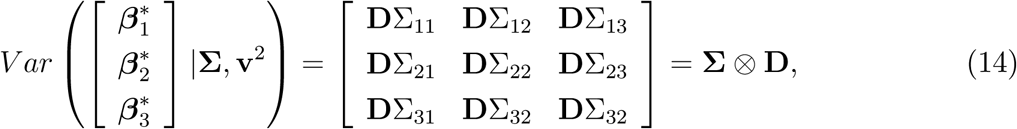

where 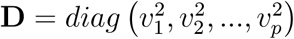 is a diagonal matri. Hence,

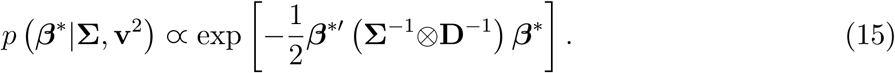

##### 3.1.3 Scale matrix Σ

The scale matrix **Σ** of MLAP can be given a fixed value (becoming a hyper-parameter) or inferred, in which case a prior distribution is needed. Here, an inverse-Wishart (*IW*) distribution with scale matrix Ω_*β*_ and *ν*_*β*_ degrees of freedom will be assigned as prior. The density is

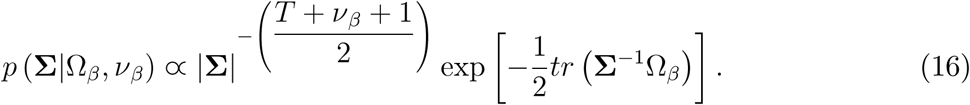

#### 3.2 Joint posterior and fully-conditional distributions

The joint posterior distribution, including 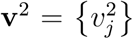 from the scale-mixture of normals representation of the prior distribution of ***β***, was assumed to take the form

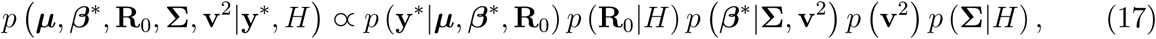

where *H* denotes the hyper-parameters; recall that **y**\* is the data vector sorted by individuals within trait The fully conditional distributions are presented below, with *ELSE* used to denote all parameters that are kept fixed, together with *H*, in a specific conditional distribution.

##### 3.2.1 Parameters *µ* and *β*\* given *ELSE*

From (17) and using representations (4) and (15), the fully conditional posterior distribution of ***µ*** and ***β***\* has density

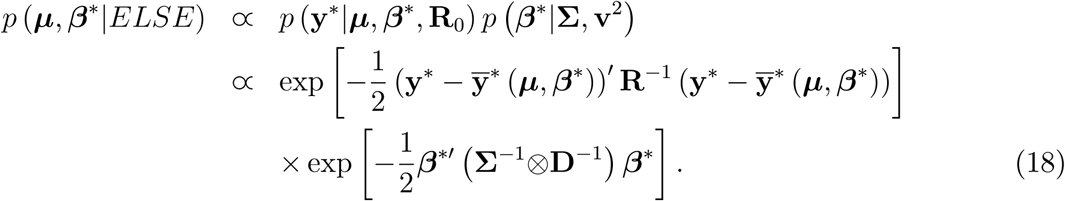

The preceding is a multivariate normal density (e.g., Sorensen and Gianola 2002). The mean vector of the distribution is

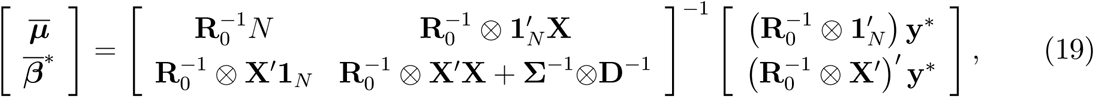

and the variance-covariance matrix is

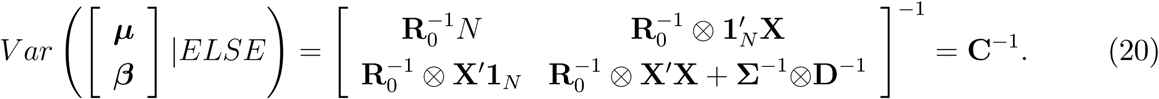

A more explicit representation is presented in Appendix B for the case *T* = 3.

##### 3.2.2 Fully conditional distributions of partitions of the location vector

For details, see Van Tassell and Van Vleck (1996) and Sorensen and Gianola (2002). Since the joint posterior of the location parameters, given **Σ, v**^2^ and **R**_0_, is multivariate normal, all conditionals and linear combinations thereof are normal as well. In particular (*T* = 3),

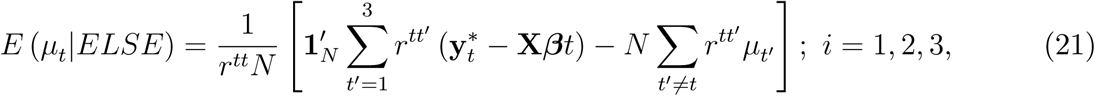

and

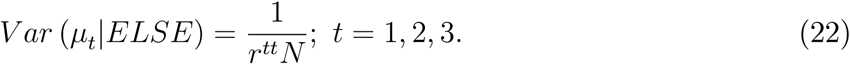

Likewise

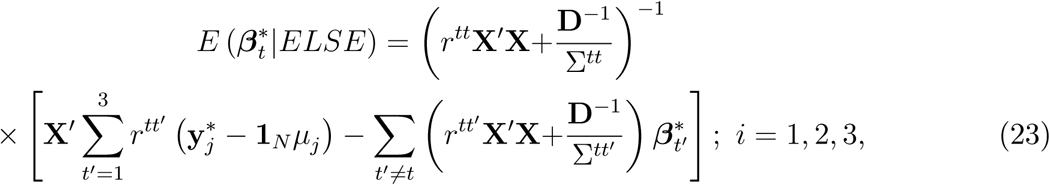

and

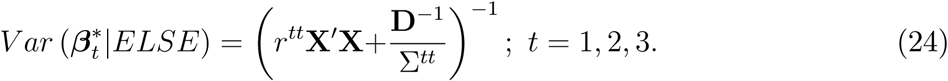

##### 3.2.3 Fully conditional distributions of R_0_ and Σ

From (17) using (2) and (6)

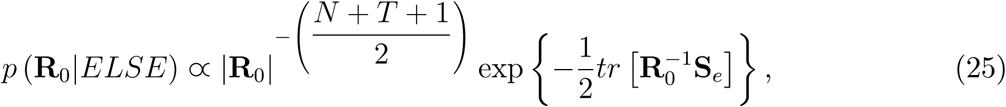

so [**R**_0_|*ELSE*] is an IW distribution with *N* + *T* degrees of freedom and scale matrix **S**_e_. In *IW*, the kernel of the density is often written as 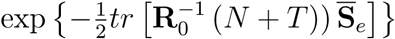, where 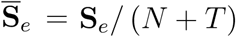.

Recall that

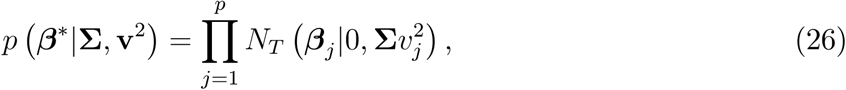

so from (17)

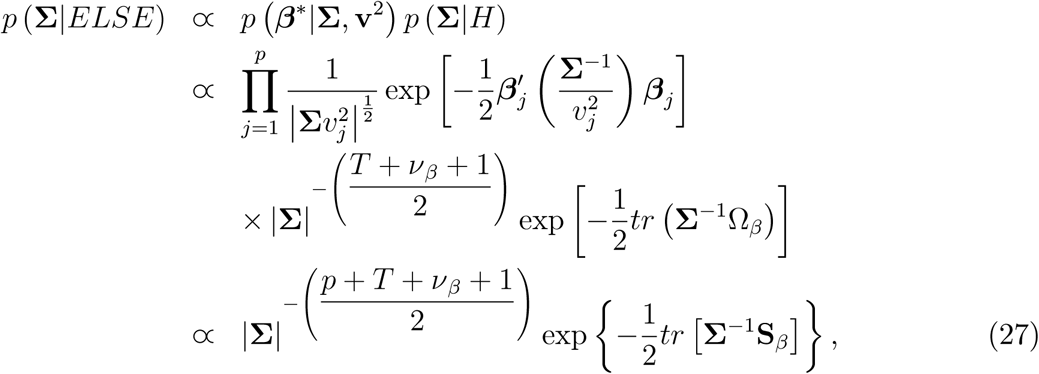

where

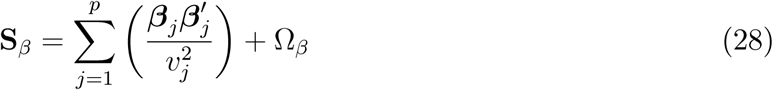

is a *T* × *T* matrix. Hence the conditional posterior distribution of **Σ** is *IW* (*p* + *T* + *ν*_*β*_, **S**_*β*_). The kernel of the density of **Σ** is often represented as 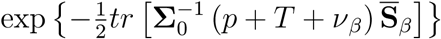, where 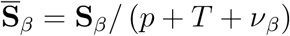.

##### 3.2.4 Fully conditional distribution of **v**^2^

From (17) and using (13)

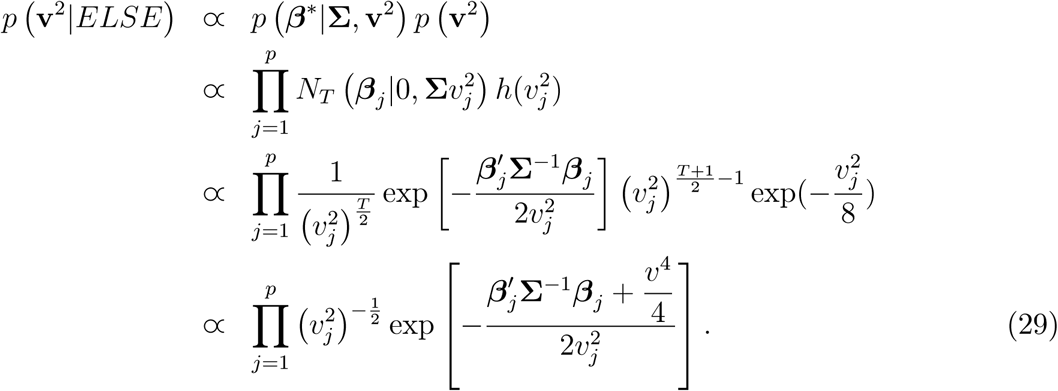

The preceding density is not in a recognizable form. Appendix C gives details of a Metropolis-Hastings algorithm tailored for making draws from the distribution having density (29). A brief description of the procedure follows.

#### 3.3 MCMC algorithm

- Starting values for **R**_0_ and **Σ** can be obtained “externally” from some estimates of **R**_0_ and **B** (the *T* × *T* matrix of variances and covariances of marker effects) calculated with standard methods such as maximum likelihood. Recall that **Σ** = **B**/[4(*T* + 1)].
- Sample each 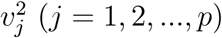 using the following Metropolis-Hastings sampler:
  1. At round *t*, draw *y* from 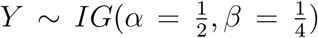 and evaluate *y* as proposal; *IG* stands for an inverse-gamma distribution.
  2. Draw *U* ∼ *U* (0, 1), with the probability of move being min(1, *R*), with *R* as in Appendix C.
  3. If *U* < *R*, set 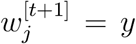 and form 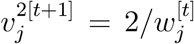 as a new state. Otherwise, set 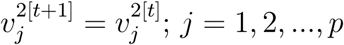.
- In a “single-pass” sampler, use (19) and (20) for sampling the entire location vector jointly. Otherwise, adopt a blocking strategy; for example draw ***µ*** and ***β***\* using (21), (22), (23) and (24).
- Sample **R**_0_ from *IW* (*N* + *T*, **S**_*e*_) and **Σ** from *IW* (*p* + *T* + *ν*_*β*_, **S**_*β*_).

#### 3.4 Remarks

Appendix E shows that the degree of shrinkage of marker effects results from a joint action between **Σ** and the strength of marker effects. A vector of effects of a marker with a short Mahalanobis distance away from **0** is more strongly shrunk towards the origin (i.e., the mean of prior distribution) than vectors containing strong effects on at least one trait. MLAP preserves the spirit of BL, producing “pseudo-selection” of covariates: all markers stay in the model, but some are effectively nullified. A marker with strong marginal and joint effects on the traits under consideration could flag potentially pleiotropic regions.

#### 3.5 Missing records for some traits

Often, not all traits are measured in all individuals, a situation that is more common in animal breeding than in plant breeding. A standard approach (“data augmentation”) treats missing phenotypes as unknowns in an expanded joint posterior distribution. As shown in Appendix F, a predictive distribution can be used to produce an imputation of missing data.

## 4 Alternative formulation in *TN* dimensions

The MCMC sampler described above is based on a regression on markers formulation stemming from either (1) or (4). In a “single-pass” sampler, *T* (1 + *p*) parameters must be drawn together; when *p* is very large, direct inversion is typically unfeasible so the scheme must be reformulated into a “block-sampling” one, i.e., by drawing some of the location parameters jointly by conditioning on the other location parameters, or by using a single-site sampler (Sorensen and Gianola 2002). Blocking or single-site sampling facilitate computation at the expense of slowing down convergence to the target distribution. Appendix D gives a scheme in which *T* (1 + *N*) effects (trait means and bivariate genomic breeding values) are inferred, and the *Tp* marker effects are calculated indirectly, following ideas of Henderson (1977) and adapted by Goddard (2009) to a genome-based model.

## 5 Data availability statement

The wheat yield data set in the `R` package BGLR (Pérez and de los Campos 2014) was employed to contrast MBL with GBLUP and Bayes C*π*. This wheat data set has been studied extensively, e.g., by Crossa et al. (2010), Gianola et al. (2011), Long et al. (2011) and Gianola et al. (2016). There are *n* = 599 wheat inbred lines, each genotyped with *p* = 1279 DArT (Diversity Array Technology) markers and each planted in four environments. The DArT markers are binary (0, 1) and denote presence or absence of an allele at a marker locus in a given line. Grain yields in environments 1 and 2 were employed to compare outcomes between analyses based on bivariate GBLUP and the bivariate BL. In the bivariate model, yields in the two environments are treated as distinct traits, conceptually, an idea that dates back to Falconer (1952). This type of setting arises in dairy cattle-breeding, where milk production of daughters of bulls in different countries are regarded as different traits and in multi-environment situations in plant breeding; both instances can be represented as special cases of a multiple-trait mixed effects model.

A publicly available Loblolly pine (*Pinus taeda*) data described in Cheng et al. (2018a) was used to carry out a predictive comparison between a Bayesian bivariate GBLUP with the bivariate Bayesian LASSO, as well as the latter versus a single-trait Bayesian LASSO. After edits, there were *n* = 807 individuals with *p* = 4828 SNP markers with measurements on rust bin scores and rust gall volume, two disease traits; see Cheng et al. (2018a).

## 6 Bivariate analysis of wheat yield: MBL versus GBLUP

### 6.1 Genomic BLUP and Bayesian BLUP

The bivariate model was

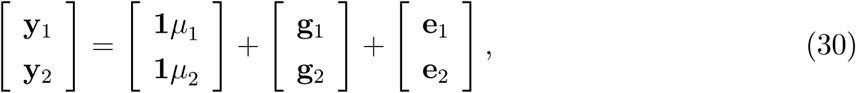

where **y**_1_ (**y**_2_) is the vector of grain yields in environment 1 (2) of the 599 inbred lines; *µ*_1_ and *µ*_2_ are the trait means in the two environments and **1** is a 599 × 1 incidence vector of ones; **g**_1_ and **g**_2_ are the “additive genomic values” of the lines and **e**_1_ and **e**_2_ are model residuals. In GBLUP (Van Raden 2008) the genetic signals captured by markers are represented as **g**_1_ = **X*β***_1_ and **g**_2_ = **X*β***_2_ where **X** is a 599 × 1279 centered and scaled matrix of genotype codes, and ***β***_1_ (***β***_2_) contains the marker allele substitution effects on trait 1 (2). The residual distribution was

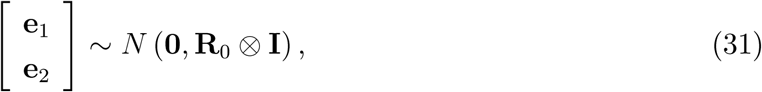

where, as-before, **R**_0_ is the 2 × 2 between-trait residual variance-covariance matrix. Effects of environment 1 are expected to be uncorrelated with those of environment 2. However, allowance was made for a non-null residual covariance because the additive genomic model may not capture extant epistasis involving additive effects, potentially creating correlations among residuals of the same lines in different environmental conditions.

GBLUP assumed 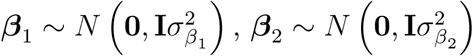 and 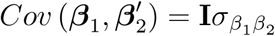, so

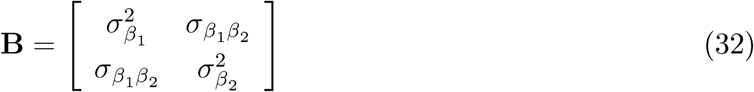

is the variance-covariance matrix of marker effects. It follows that

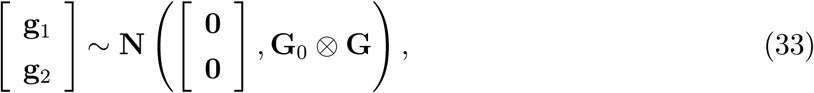

where

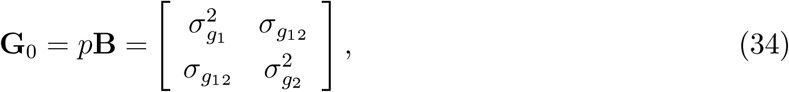

is a between-trait variance-covariance matrix of the additive genomic values (here, e.g., 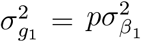) and **G** = **XX**′/*p* is a genomic-relationship matrix describing genome-based similarities among the 599 lines. The preceding assumptions induce the marginal distribution

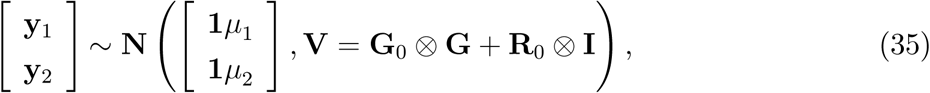

where **V** is the phenotypic covariance-matrix. The bivariate best linear unbiased predictor of **g**_1_ and **g**_2_ (Henderson 1975)

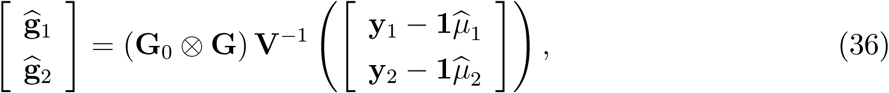

where

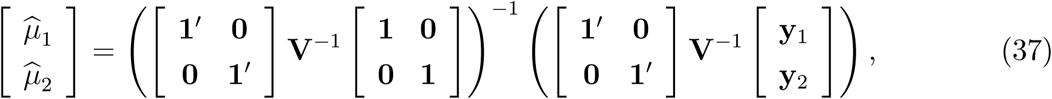

is the bivariate generalized least-squares (GLS) estimator of the trait means.

BLUP and GLS require knowledge of **G**_0_ and **R**_0_ and we replaced these unknown matrices by estimates obtained using a crude but simple procedure. Genomic and residual variance components were obtained by univariate maximum likelihood analyses of traits 1, 2 and 1 + 2, and covariance component estimates were calculated from the expression *Cov*(*X, Y*) = [*Var*(*X* + *Y*) − *Var*(*X*) − *Var*(*Y*)] /2. The resulting estimates of **G**_0_ and **R**_0_ were inside of their corresponding parameter spaces. An estimate of **B** was obtained by applying relationship (34) to the estimated **G**_0_.

Henderson (1977) showed how BLUP of vectors that are not likelihood identified can be obtained from best linear unbiased predictions of likelihood-identified random effects (see Gianola 2013). Goddard (2009) and Strandén and Garrick (2009) used this property to obtain predictions of marker effects (***β***) given predictions of signal (g). If ***β*** and g have a joint normal distribution, under (30) one has

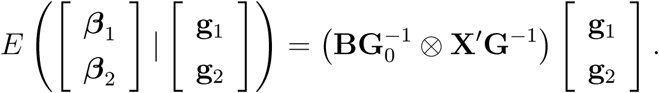

Using iterated expectations and recalling that BLUP can be viewed as an estimated conditional expectation (with fixed effects replaced by their GLS estimates), BLUP of marker effects is expressible as

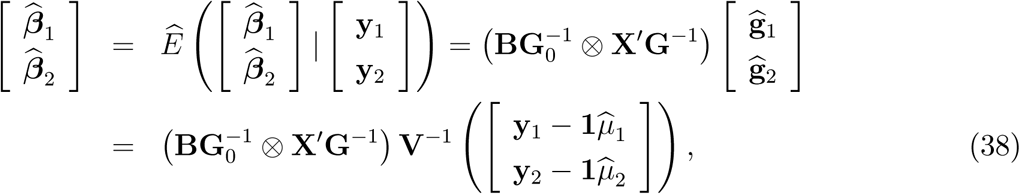

With 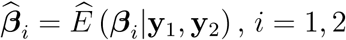. After lengthy algebra and using Henderson (1975), the prediction error variance-covariance matrix of the BLUP of marker effects is given by

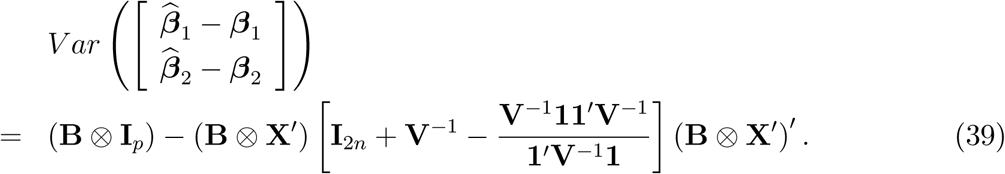

A set of *t* − *statistics* can be formed by taking the ratio between the BLUP of a given marker effect as in (38) and the square root of the corresponding diagonal element of (39). The statistic is a crude criterion for association between marker and phenotype as it ignores uncertainty associated with the fact that **B** and **R**_0_ are estimated from the data, as opposed to being “true values” required by BLUP theory.

The Bayesian bivariate GBLUP model used standard assumption as in Sorensen and Gianola (2002), i.e., it was a multivariate normal-inverse Wishart hierarchical specification. The only difference with GBLUP is that, in the Bayesian treatment, **G**_0_ and **R**_0_ were treated as unknown parameters, with the uncertainty about their values accounted for.

### 6.2 Bivariate LASSO

Our MCMC implementation for MBL was applied to markers directly, as opposed to inferring their effects from signal indirectly, as it is done for GBLUP. The model was as in (4) with *T* = 2. Each marker was assigned a conditional bivariate Laplace prior distribution with scale matrix **Σ**; in turn, **Σ** was given a two-dimensional inverse Wishart distribution on *ν*_*β*_ = 4 degrees of freedom and with scale matrix **Ω**_*β*_ = *ν*_*β*_**B**/12 = **B**/3. The residual variance-covariance matrix **R**_0_ was assigned the two-dimensional Jeffreys improper prior in (6).

The MCMC scheme employed the scale mixture of normals representation of the bivariate Laplace distribution. First, six independent chains of 1500 iterations each were run. The shrinkage diagnostic metric of Gelman and Rubin (1992) was calculated for *µ*_1_, *µ*_2_, **R**_0_ and **Σ**, for the effect of marker 10 on trait 1, and for the effect of marker 200 on trait 2; the R package CODA was used for this purpose. Supplementary Figures 4-13 gave no strong evidence of lack of convergence, as indicated by shrinkage factor values close to 1.

Post-burn in samples were collected for an additional 2000 iterations in each chain, so a total of 12,000 samples (without thinning) was used for inference. Supplementary Figures 14 and 15 depict post burn-in trace plots for the elements of **R**_0_ and **Σ**, respectively. The six chains “joined” eventually and sample values thereafter fluctuated within what seemed to be stationary distributions. To assess convergence further, a test suggested by Geweke (1992) was applied to the combined 12000 samples from the posterior distributions of 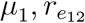 (residual correlation between yields in environments 1 and 2) and 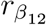, the correlation between effects of a marker on the two traits. The test compared means of two parts of the combined collection of 12,000 samples at each of 10 segments of the collection: there was no evidence of lack of convergence. In short, the implementation met successfully the convergence tests applied.

Figures 1 and 2 display estimated posterior densities of 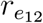 and 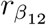. Mixing for 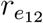 was poorer than for 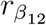; the effective number of samples was 220.6 and 979.0, respectively, and Monte Carlo errors were small enough. The residual correlation (posterior mean, 0.17) was positive and different from 0, whereas the 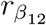 parameter was estimated at −0.35, also different from zero. However, the posterior densities were not sharp enough for precise inference, probably due to the small sample size (*n* = 599) and low density of the marker panel (*p* = 1279). The quality of these estimates is of subsidiary interest here as our objective was to demonstrate the MBL in a comparison with bibariate BLUP of marker effects.

**Figure 1.**
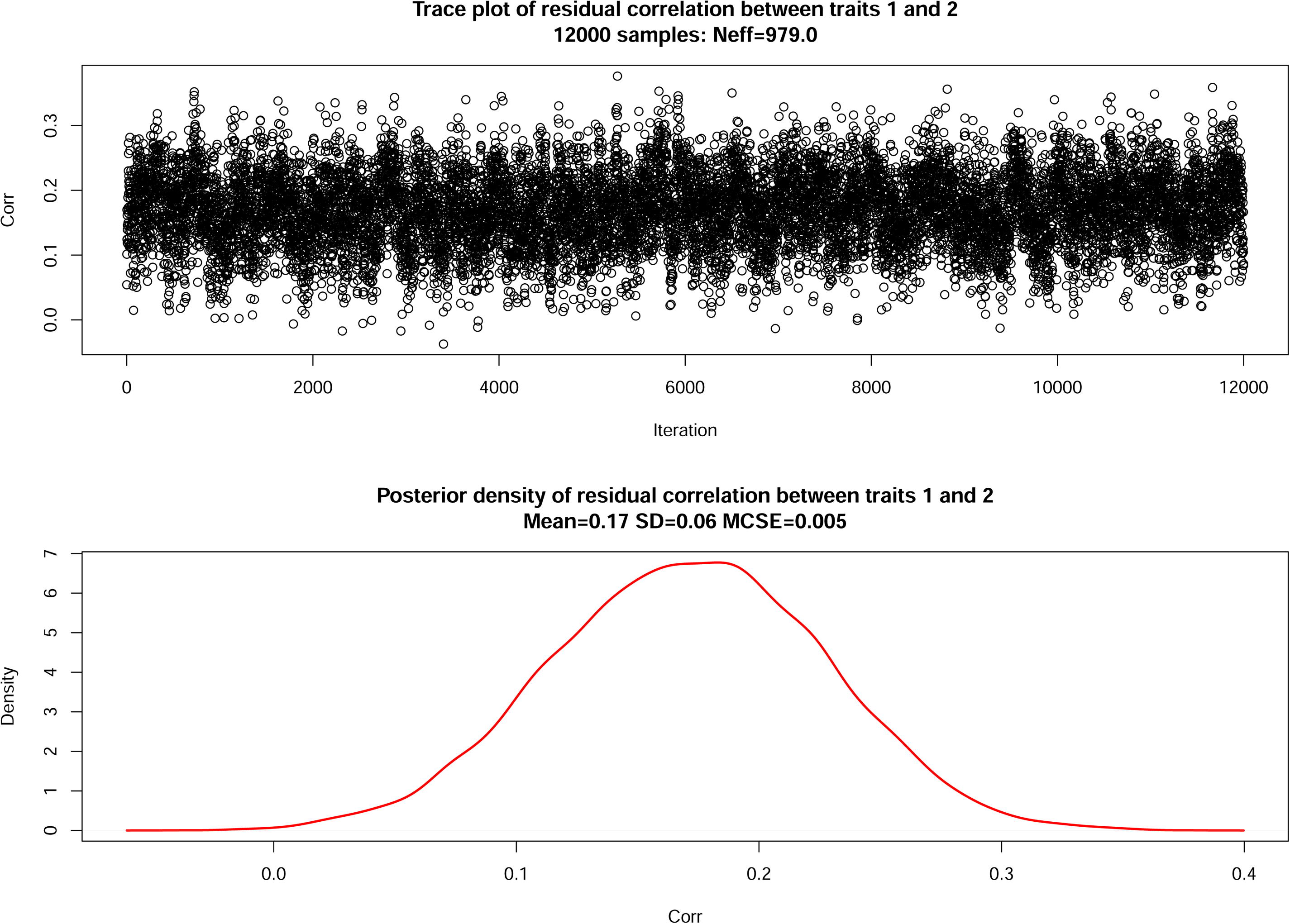
Bivariate Bayesian LASSO: trace plot and posterior density of residual correlation.

**Figure 2.**
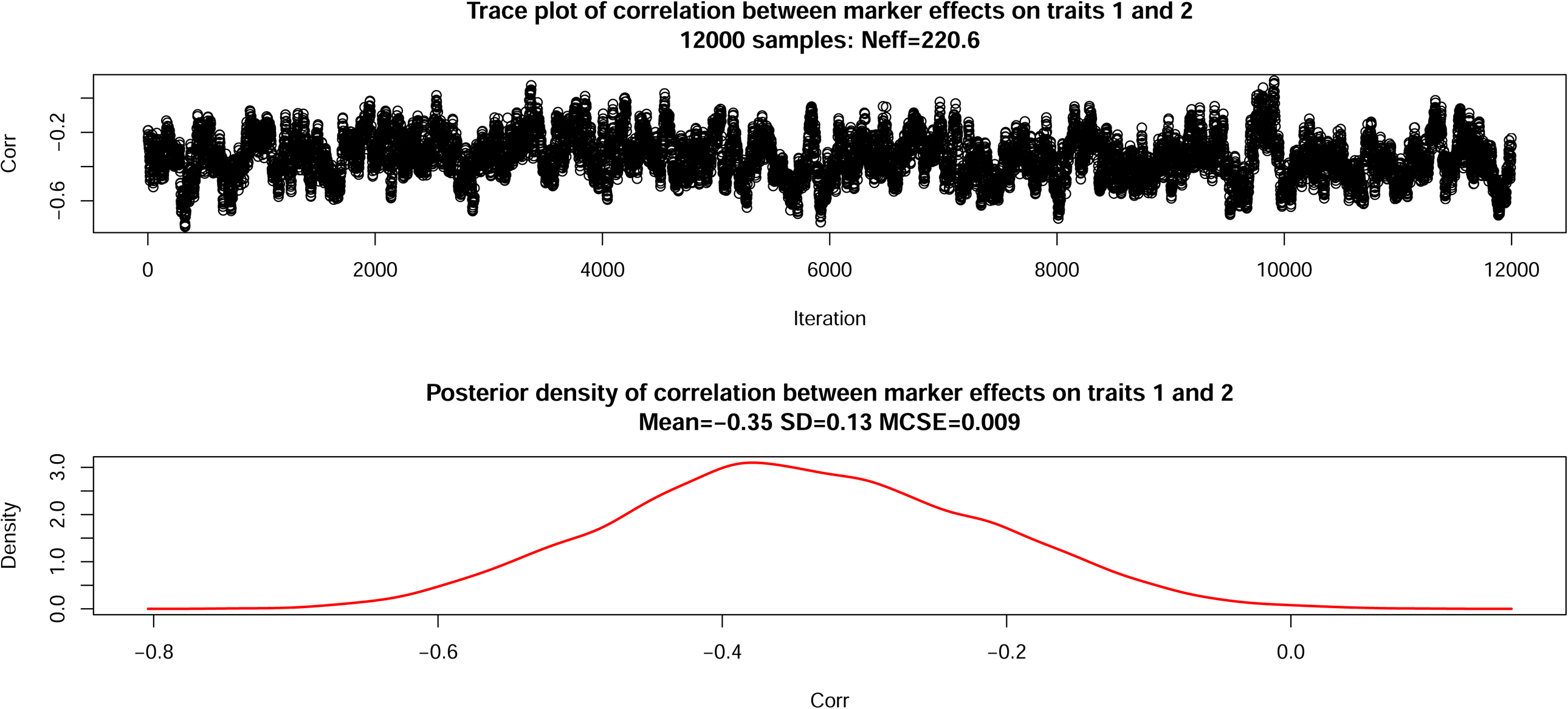
Bivariate Bayesian LASSO: trace plot and posterior density of correlation between marker effects.

Location parameters mixed well. For example, the average effective sample size of *µ*_1_ over the 6 chains during burn in was 1499 for a nominal 1500 iterations. For marker 10 effect on trait 1 it was 962 out of 1500, and for marker 200 effect on trait 2 effective size was 1130 out of 1500. These numbers suggest that all 2558 marker effects were estimated with a very small Monte Carlo error in our MBL implementation with 12,000 samples used for inference.

### 6.3 MBL vs BLUP estimates of marker effects

Figure 3 gives a comparison between bivariate BLUP and posterior mean estimates of effects from MBL. The upper panel shows good alignment between estimates, except at the extremes of the scatter plots. The lower panel depicts that markers with the strongest absolute effects, as estimated by BLUP, had an even stronger effect when estimated under the bivariate BL. Figure 4 presents standardized estimates of each of the 1279 marker effects, by trait. For GBLUP the *t* − *statistic* was the estimated marker effect divided by the square root of its prediction error variance; for MBL it was the posterior mean divided by its posterior standard deviation. There is no evidence that any of the markers had an effect differing from 0, corroborating the view that wheat yield is a typical quantitative trait affected by many variants each having a small effects (Singh et al. 1986; Sleper and Poehlman 2006). Using a univariate least-squares, GWAS-type analysis, there were 29 (yield 1) and 56 (yield 2) significant hits after a Bonferroni correction (1279 tests, *α* = 0.05). A comparison between the *t* − *statistics* from the GWAS-type analysis with the standardized BLUP and MBL effects is provided in Figure 5. As expected, shrinkage towards null-mean distributions (bivariate Gaussian in BLUP and bivariate Laplace in MBL) made *t* − *statistics* much smaller in absolute value than the corresponding ones from GWAS.

**Figure 3.**
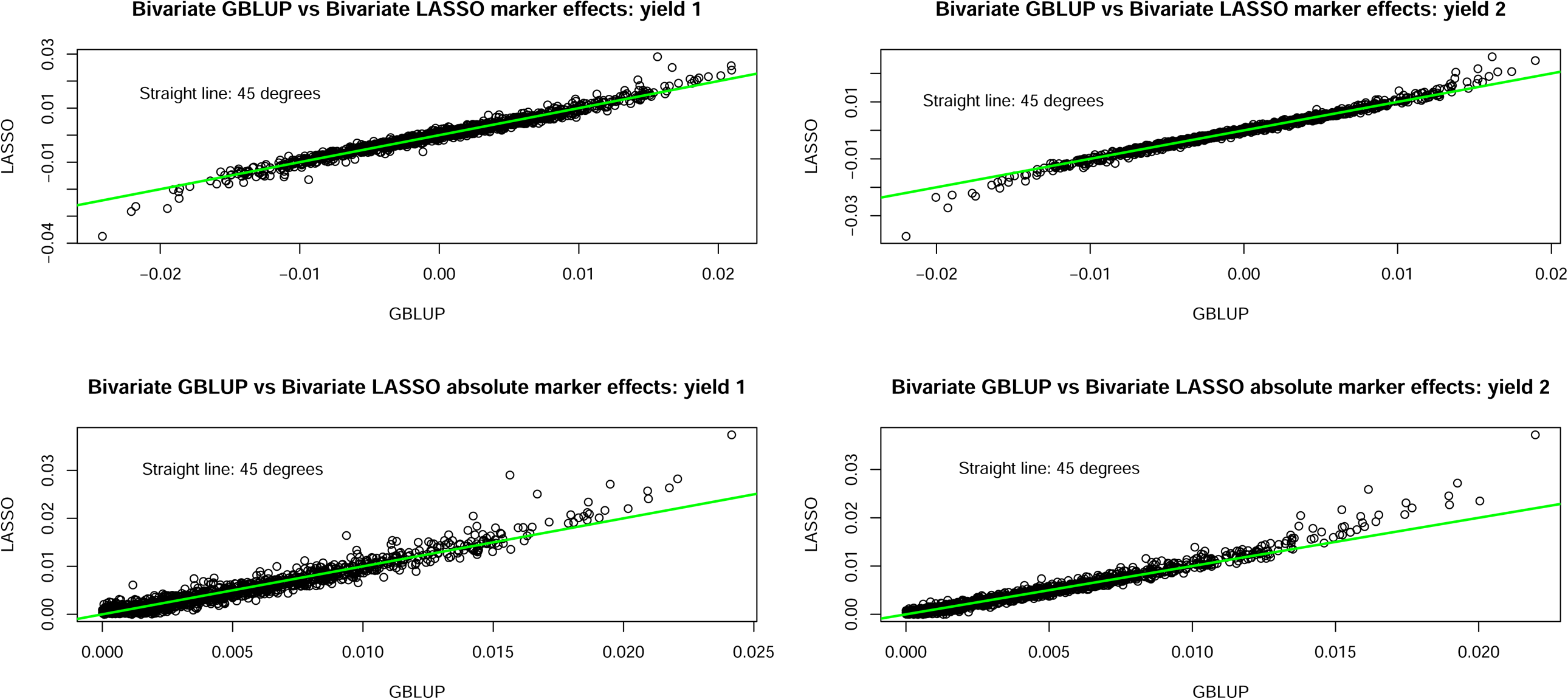
Bivariate GBLUP versus bivariate Bayesian LASSO (posterior mean) estimates of marker effects on wheat grain yield.

**Figure 4.**
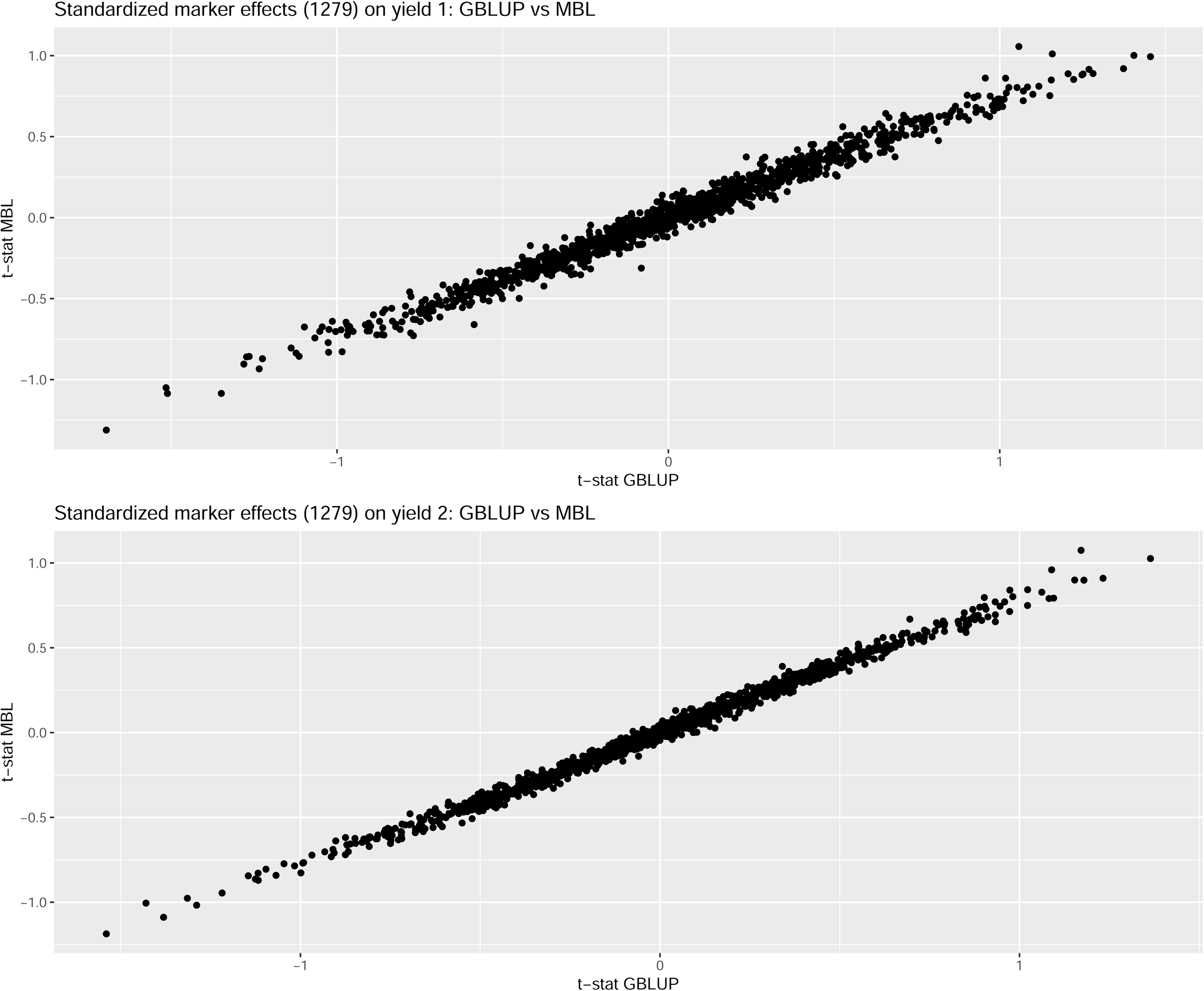
*t*-statistics for marker effects on wheat grain yield: GBLUP versus bivariate Bayesian LASSO (MBL)

**Figure 5.**
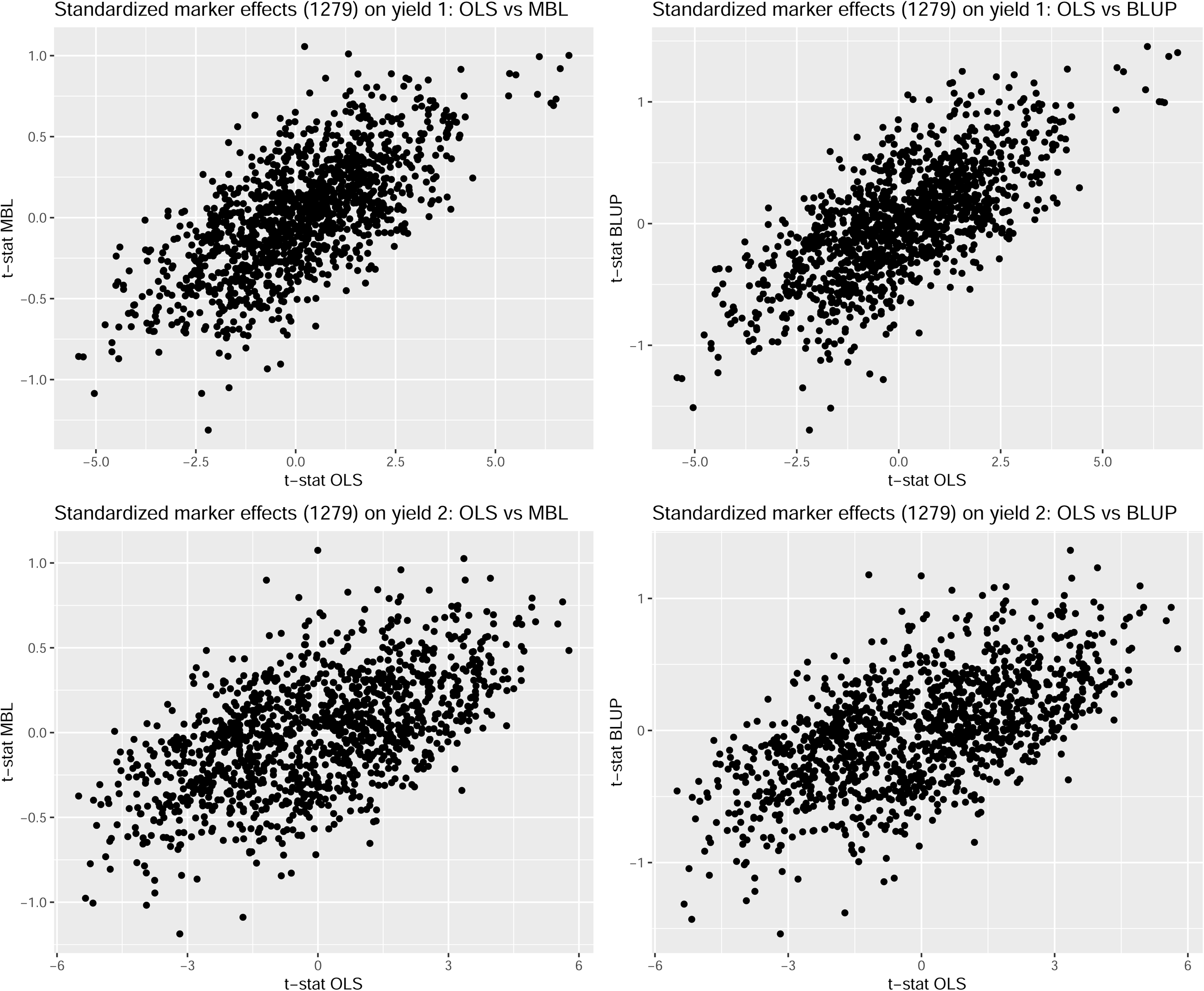
*t*-statistics for marker effects on wheat grain yield: ordinary leas-squares (OLS) versus bivariate Bayesian LASSO (MBL) and bivariate GBLUP.

Standard GWAS aims to find connections between markers and genomic regions having an effect on a single trait (e.g., Manolio et al. 2009, Visscher et al. 2012; Gianola et al. 2016; Schaid et al. 2018) A search for pleiotropy, on the other hand, focuses on markers having multi-trait effects. The latter can be viewed as a search for vectors of effects with non-null coordinates that are distant from a *T*–dimensional 0 origin. Mahalanobis squared distances 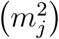 away from (0, 0) for each the 1279 bivariate vectors of marker effects were calculated for both BLUP and MBL. For BLUP and marker *j*, the squared distance was computed as 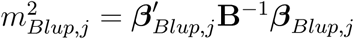, and for MBL it was 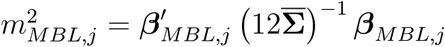 where ***β***_.,*j*_ are effect estimates for marker *j* and 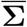 is the estimated posterior expectation of **Σ**. For BLUP, 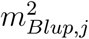 had median and maximum values of 0.16 and 2.94, respectively, over markers. For MBL the corresponding values were 0.14 and 3.83. Figure 6 shows that the largest estimated distances were obtained with MBL, supporting the view that the method produces less shrinkage of multiple-trait effect sizes than BLUP. If the 95% percentile of a chi-squared distribution on 2 degrees of freedom (5.99 and 14.4 without and with a Bonferroni correction at *α* = 0.05) is used as “significance threshold”, none of the 1279 markers could be claimed to have a bivariate effect on the trait, which is consistent with the *t* − *statistics*.

**Figure 6.**
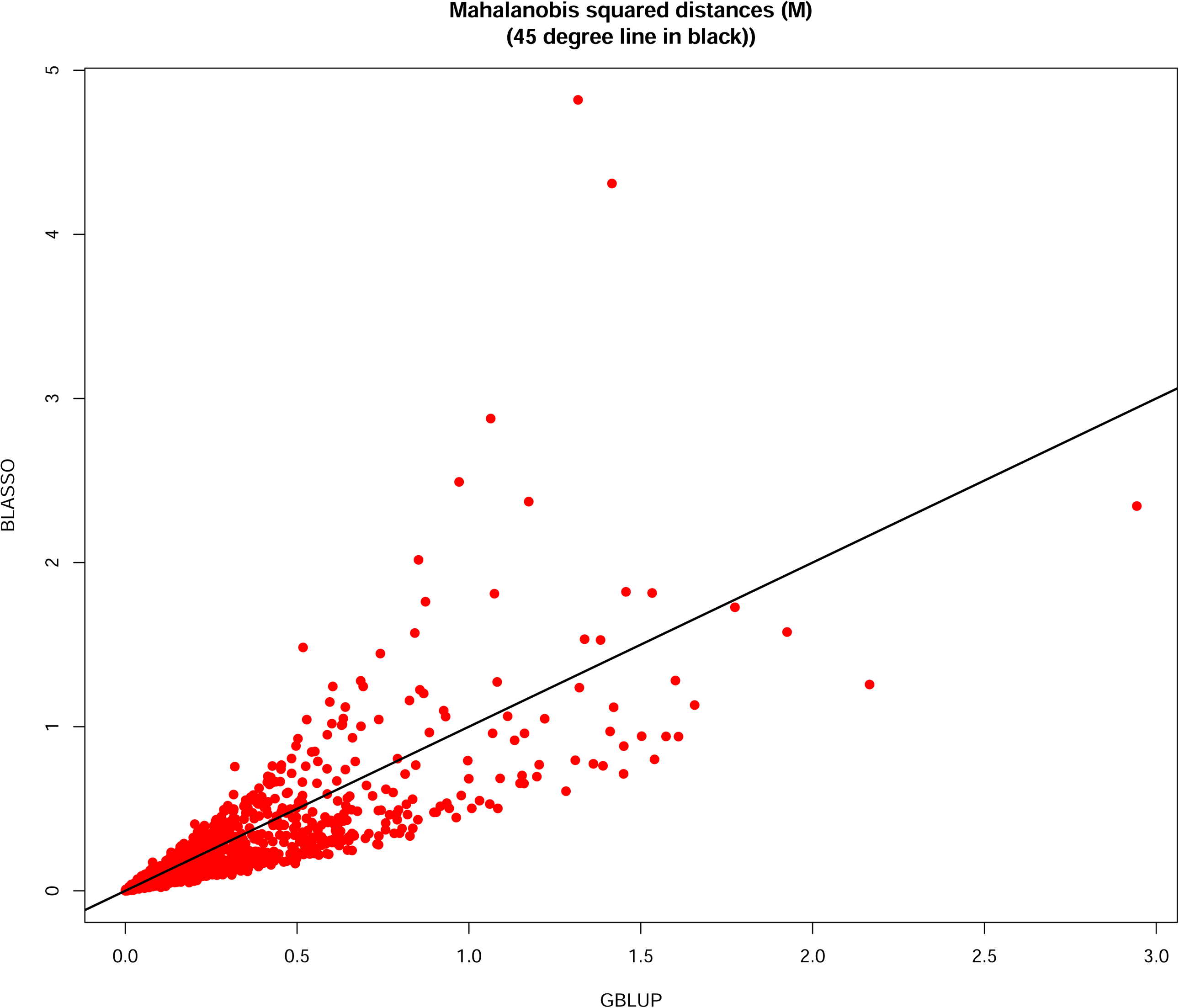
Mahalanobis squared distance (M) away from (0,0) for bivariate effects on grain yield of 1279 markers: GBLUP versus bivariate Bayesian LASSO (BLASSO)

## 7 Predictive comparison between MBL, MTGBLUP and MT-BayesC*π*: wheat

Bivariate Bayesian GBLUP and BayesC*π* models (Cheng et al. 2018a) were also fitted to the wheat data set. Multiple-trait Bayesian linear models are well known (e.g., Sorensen and Gianola 2002); BayesC*π* consisted of a bivariate mixture in which each of the 1279 markers was allowed to fall, *a priori*, into one of four disjoint classes: (0, 0), (0, 1), (1, 0), (1, 1), where (0, 0) means that a marker has no effect on either trait, (0, 1) indicates that a marker affects yield 2 only, and so on. The prior for the four probabilities of membership was a *Dirichlet* (1, 1, 1, 1) distribution. All three methods were run in each of 100 randomly constructed training sets and predictions were formed for lines in corresponding testing sets. Training and testing set sizes had 300 and 299 wheat lines, respectively, in each of the 100 runs. For all methods, the MCMC scheme was a single long chain of 50,000 iterations, with a burn-in period of 1,000 draws. The analyses were run using the JWAS package written in the JULIA language (Cheng et al. 2018b).

Figures 7 and 8 present pairwise plots (bivariate Bayesian GBLUP denoted as RR-BLUP in the plots) of predictive correlations and predictive mean-squared errors, respectively; the plots display less than 100 (*X, Y*) points because numbers were rounded to two decimal points. There were no appreciable differences in predictive performance between the three methods, supporting the view that cereal grain yield is multi-factorial and that there are none, if any, genomic regions, with large effects. The variability among replications of the training-testing layout is essentially random, reinforcing the notion of the importance of measuring uncertainty of prediction (Gianola et al. 2018). Many studies fail to replicate and often claim differences between methods based on single realizations of predictive analyses.

**Figure 7.**
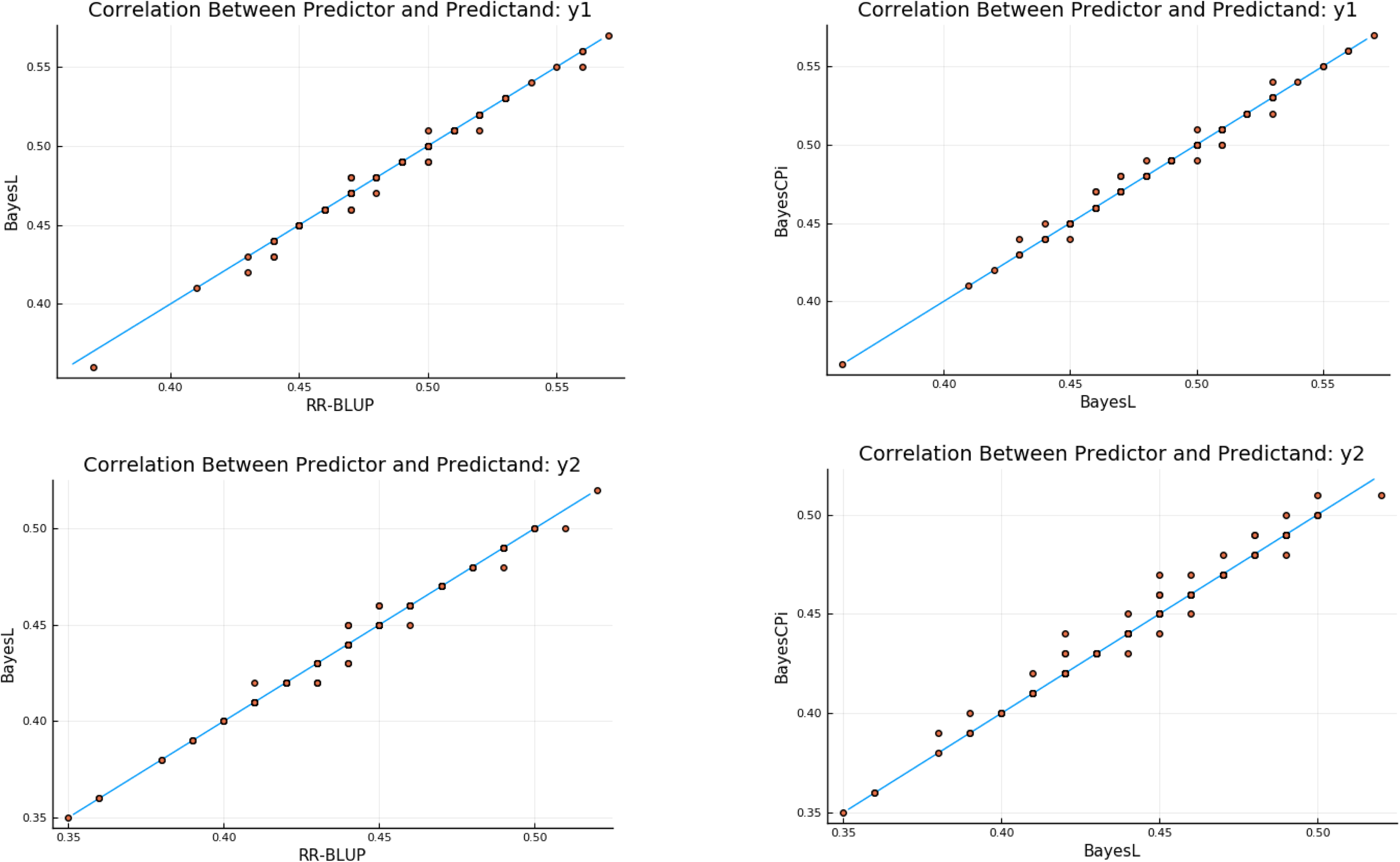
Predictive correlations for wheat grain yield: bivariate Bayesian LASSO (Bayes L) versus bivariate GBLUP (RR-BLUP).

**Figure 8.**
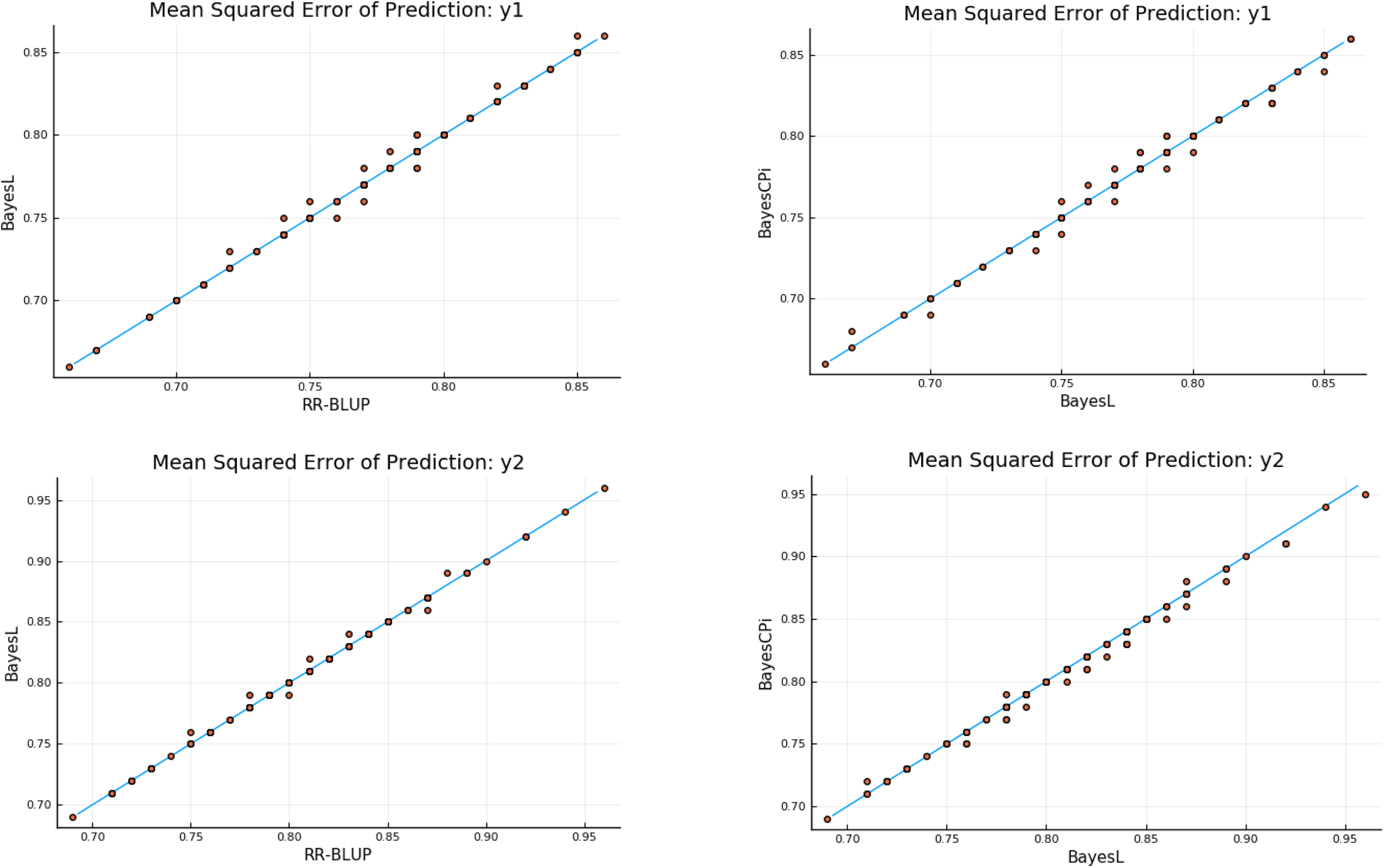
Predictive mean-squared error for grain yield: bivariate Bayesian LASSO (Bayes L), bivariate GBLUP (RR-BLUP) and bivariate Bayes C*π*.

## 8 Predictive comparison between MBL vs MTGBLUP and MBL vs single trait Bayesian LASSO: *Pinus*

Figures 9 and 10 present scatter-plots of the predictive performance (mean squared error and correlation, respectively) of the bivariate Bayesian LASSO and bivariate Bayesian GBLUP (MT-GBLUP, denoted as RR-BLUP in the plots) in the 100 testing sets. There were no obvious differences in mean-squared error for either rust bin or gall volume although, for the latter trait, a slight superiority of MBL was noted (Figure 9); the plot contains distinct 12 points because the overlap in numerical values produced “clusters” of points. On the other hand, there was a decisive superiority (Figure 10) of MBL over MTGBLUP in predictive correlation.

**Figure 9.**
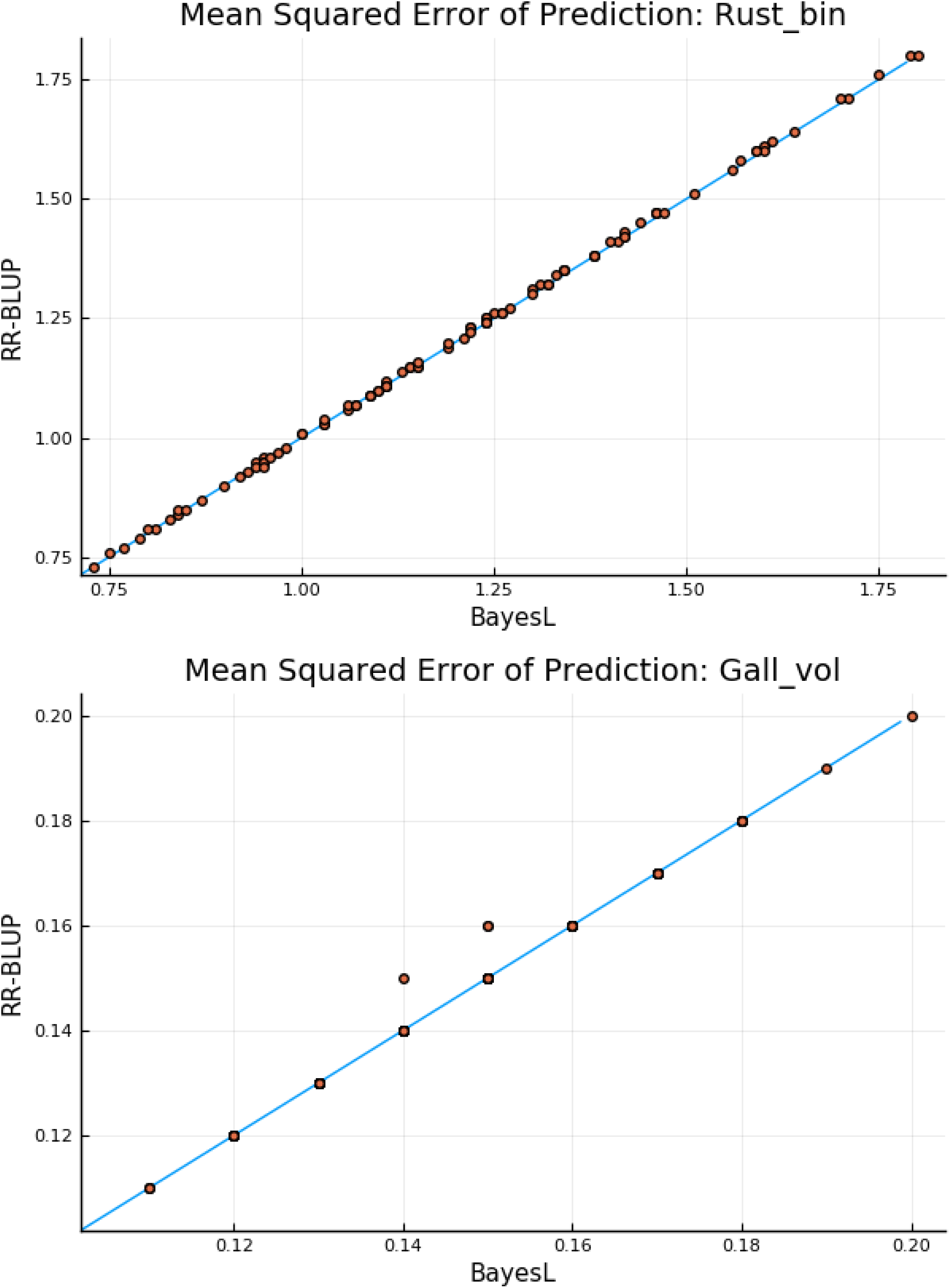
Mean-squared error of prediction for rust bin and gall volume in pine trees: bivariate Bayesian LASSO (Bayes L) versus bivariate Bayesian GBLUP (RR-BLUP).

**Figure 10.**
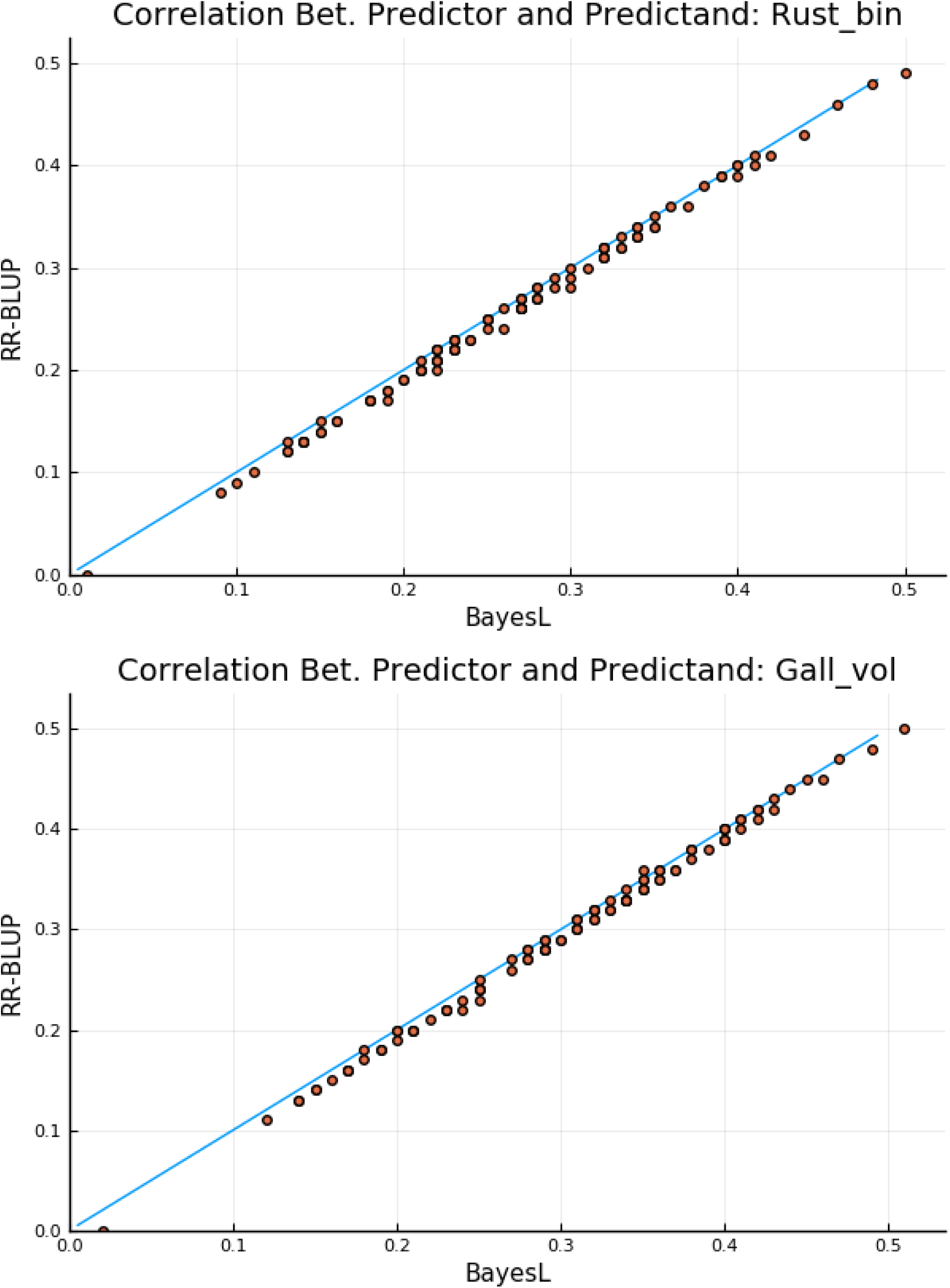
Predictive correlation for rust bin and gall volume in pine trees: bivariate Bayesian LASSO (Bayes L) versus bivariate Bayesian GBLUP (RR-BLUP).

Figure 11 contrasts the predictive performance of the bivariate Bayesian LASSO over the single trait Bayesian LASSO for gall volume. The two trait analysis tended to produce larger predictive correlations and smaller mean-squared errors, illustrating an instances in which a multiple-trait specification clearly constitutes a better prediction machine.

**Figure 11.**
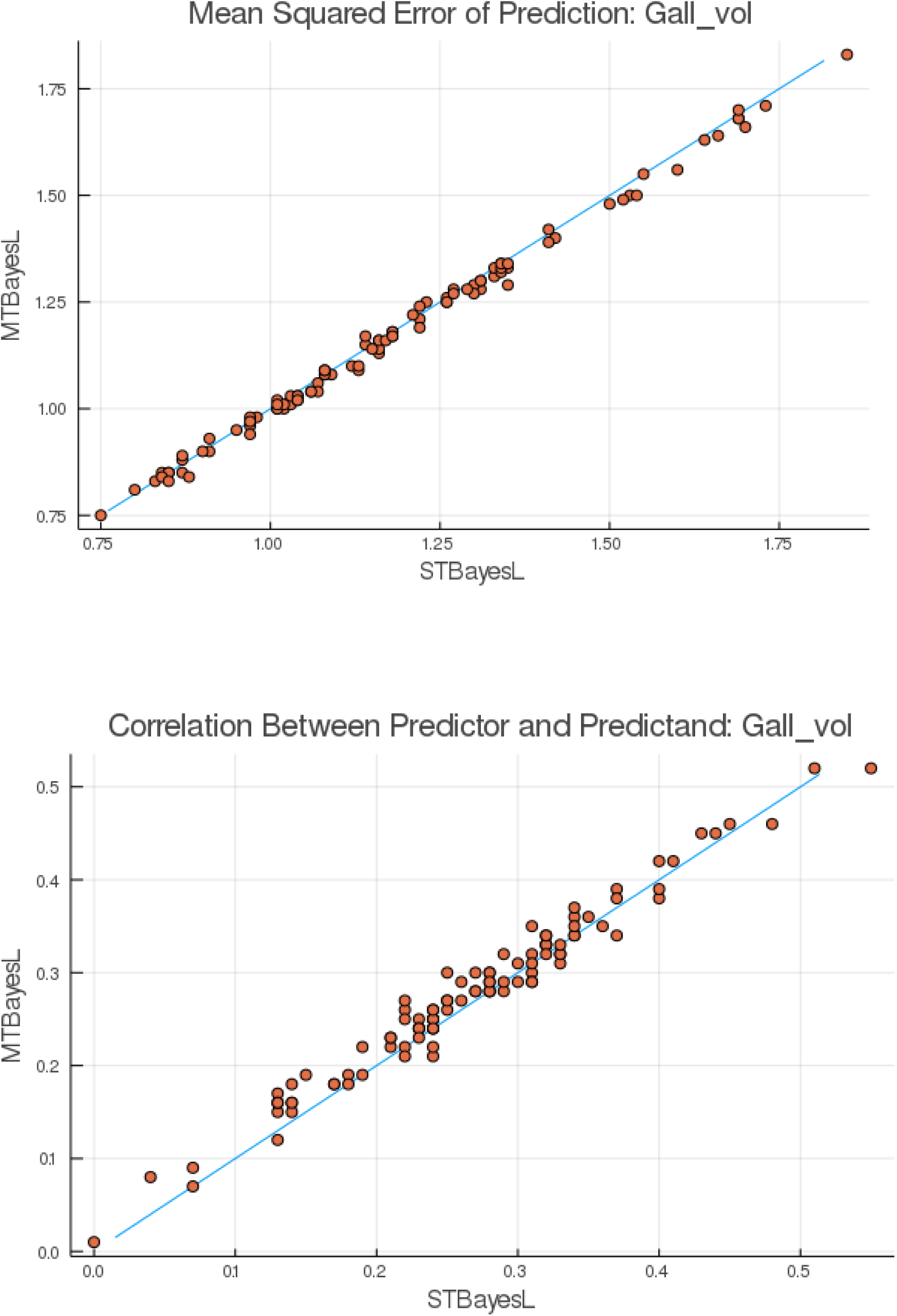
Predictive mean squared error and correlation for gall volue in pine trees: bivariate Bayesian LASSO (MTBayesL) versus univariate Bayesian LASSO (STBayesL)

## 9 Conclusion

Our study is possibly the first report in the quantitative genetics literature of a multiple-trait Bayesian LASSO (MBL), inspired by the BL of Park and Casella (2008). MBL assumes that vectors of marker effects on *T* traits follow a null-mean multivariate Laplace distribution, *a priori*. This distribution has a sharp peak at the origin and reduces to the double exponential prior of the BL when applied to a single trait. The implementation of MBL requires Markov chain Monte Carlo sampling and a relative simple Metropolis-Hastings algorithm based on a scaled mixture of normals representation (Gómez-Sánchez-Manzano et al. 2008) was presented. The algorithm was tested thoroughly with a wheat data set and found to mix well, with no evidence of lack of convergence to the posterior distribution and with a small Monte Carlo error.

A question that arises often in practice, is the extent to which a multiple-trait method will produce a better performance than a single-trait specification. If the parameters of the model (assuming it holds) representing the inter-trait distribution are either known or well estimated, one should expect more power for QTL detection and a better predictive performance for the multivariate specification. In our study we found that MBL outperformed the single trait in terms of delivering a better predictive performance for gall volume but not for rust bin in *Pinus*. On the other hand, a multiple-trait analysis is more complex and requires more assumptions, so it may be less robust than a single trait procedure and fail to deliver according to expectation in real-life circumstances. It is risky to make sweeping statements arguing in favor of a specific treatment of data as outcomes are heavily dependent on the biological architecture of the traits considered, and on the data structure as well. The picture emerging from two decades of experience in genome-enabled prediction in the fields of animal and plant breeding is that is largely futile to categorize methods in terms of expected predictive performance using broad criteria, in view of the large variability of performance with respect to data structure for any given prediction machine (Morota and Gianola 2014; Gianola and Rosa 2015; Momen et al. 2018; Montesinos-López et al. 2019 a,b,c,d; Azodi et al. 2019).

MBL is expected to shrink more strongly towards zero vectors of markers with small effects in their coordinates, thus producing differential shrinkage and preserving the *modus operandi* of BL. Mimicking the single-trait argument in Tibshirani (1996) which shows equivalence between LASSO and a posterior mode, the representation in Appendix E illustrates that the degree of shrinkage of the vectorial effects of a marker (*j*, say) on a set of traits is inversely proportional to the quadratic form 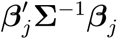. Thus, multivariate Bayesian pseudo-sparsity is induced by MBL to an extent depending on the heterogeneity of 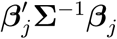 over markers. We note, in passing, that the term 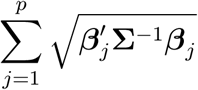 given in (66) of Appendix E is the counterpart of 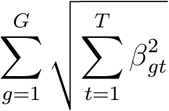, part of the “group-penalty” in Li et al. (2015), where *g* is some meaningful group of markers arrived at, say, on the basis of biological considerations, and 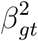 is the group regression coefficient for trait *t*. The latter penalty assigns the same weight to these regressions over trains, contrary to MBL where weights and co-weights are driven by **Σ**^−1^. The BL or MBL can be adapted to situations where a group structure may be needed via hierarchical modeling; this fairly straightforward issue is outside of the scope of the paper but may pursued in future extensions of MBL. Actually, Liquet et al. (2017) described a Bayesian multiple-trait analysis where a LASSO-type penalty is assigned to group effects and a spike-slab prior induces additional Bayesian sparsity at the level of individual regression coefficients. The authors did not address the predictive ability of their method so it would be interesting to compare it against MBL and the multiple-trait mixture model of Cheng et al. (2018). We plan to carry out this comparison in collaboration with CIMMYT (Centro Internacional de Mejoramiento de Maíz y Trigo, México) using a large number of data sets in various cereal crops.

Knowledge of the genetic basis of complex traits is limited and not vast enough to enable formulation of *a priori* prescriptions for any specific trait or situation. The number, location and effects of causal variants, the linkage disequilibrium structure between such variants and markers, and the mode of gene action of QTL are largely unknown, this holding for all species of domesticated plants and animals and for most common diseases in humans. Theoretically, MBL is expected to perform better than multiple-trait BLUP whenever appreciable heterogeneity exists over the effects of the markers in the panel employed, while behaving as multiple-trait GBLUP when all markers have tiny and similar effects. This consideration follows directly from the structure of the method, and computer simulations could be easily tailored to create scenarios where MBL has a better or a worse performance simply by design but without necessarily being relevant to a real-life inferential or predictive problem.

The expectation stated above was verified empirically: markers with stronger (positive or negative) effects on the wheat yields examined had larger Mahalanobis distances away from zero than markers with small effects. Further, markers with short distances in GBLUP had even shorter distances under MBL. None of the two methods was able to detect variants having a strong effect on wheat yield, contrary to least-squares GWAS. However, outcomes from GWAS are not strictly comparable with those from shrinkage-based procedures. In single-marker least-squares the estimator is potentially biased because other genomic regions are ignored in the model; further, short and long range linkage disequilibria create statistical ambiguity (Gianola et al. 2016). In WGR, on the other hand, regressions are akin to partial derivatives, i.e., the coefficient gives the net effect of the marker given that the other markers are fitted; typically, regressions become smaller as *p* is increased at a fixed *n*.

In plant and animal breeding, a focal point is the evaluation of genetic merit of candidates for artificial selection, and the prediction of expected performance in either collateral relatives or in descendants. Under the assumptions of additive inheritance, genome-enabled prediction (Meuwissen et al. 2001) produces estimates of marked additive genomic value, **g**, or signal as referred to in our paper. In MBL, **g** and marker effects can be inferred from their posterior mean or from a modal approximation (MAP-MBL) that does not involve MCMC which is described in Appendix E. A rough comparison between GBLUP, MBL and MAP-MBL was carried out with the wheat data. For the latter, we used **Σ** = **G**_0_/(12*p*), and starting values for the iteration were calculated using BLUP estimates of marker effects. MAP with *T* = 2 were iterated for 500 rounds. Supplementary Figure S16 shows that, at iteration 500, the metric used for monitoring convergence had stabilized at the third decimal place, but iteration could have stopped after 200 rounds, for our purposes. Supplementary Figure S17 presents a scatter plot of the 2558 (bivariate) marker effect solutions at iterations 1 and 500 against the corresponding BLUP or MBL posterior mean estimates. Clearly, MAP approach gave markedly different results, producing a stronger shrinkage to 0 of small-effect markers an, thus, an effectively more sparse model. Supplementary Figure S18 gives a comparison of the fitted genomic values, i.e., 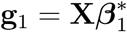 and 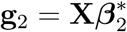 for the two traits. GBLUP and MBL estimates were closely aligned and fitted the data in a similar manner. On the other hand, MAP-MBL gave a larger mean-squared error of fit and a smaller correlation between fitted and observed phenotypes, possibly because of the larger effective sparsity of MAP-MBL. A worse fit to the data does not necessarily imply a poorer predictive ability. A thorough comparison of predictive ability between MBL and MAP-MBL will be carried out in future research.

Our predictive comparison in wheat involved three bivariate models: GBLUP, MBL and BayesC*π*, which employs Bayesian model averaging. A training-testing validation replicated 100 times at random indicated no differences among methods. However, it was found that MBL was better than MT Bayesian BLUP for the two pine tree traits. After almost two decades of genome-enabled prediction it is now clear that no universally best prediction machine exists (Gianola et al. 2011; Heslot 2012; de los Campos et al. 2013; Momen et al. 2018; Bellot et al. 2018; Montesinos-López et al. 2018a, b, c, d) even when non-parametric or deep learning techniques are brought into the comparisons.

As far as we know, our paper represents the first report in the quantitative genetics literature of a multiple-trait LASSO, implemented in a Bayesian or empirical Bayes (Appendix E) manner. MBL adds to the armamentarium of genome-enabled prediction and expands the family of members of the Bayesian alphabet (Gianola et al. 2009; Habier et al. 2011; Gianola 2013). Further, it has been implemented in the publicly available JWAS software (Cheng et al. 2018b). We take the view that every prediction problem is unique and that no claims about the superiority of a specific procedure over others should be made without qualifications. For instance, MBL could perform worse or better than here when applied to other species, traits, or when confronted with different data structures. Most quantitative and disease traits are truly complex and it is dangerous to offer simplistic solutions or predictive panaceas (Goddard et al. 2019).

## 10 Acknowledgments

Professor Chris-Carolin is thanked for useful discussions and comments. J. M. Marín provided technical information on the MLAP distribution. Two anonymous reviewers are thanked for helpful suggestions. Financial support to DG was provided by the Deutsche Forschungsgemeinschaft (grant No. SCHO 690/4-1 to CCS) and by the J. Lush Endowment as Visiting Professor at Iowa State University in 2018. The University of Wisconsin-Madison, USA, the Technical University of Munich (TUM) Institute for Advanced Study, TUM School of Life Sciences, the Institut Pasteur de Montevideo, Uruguay, are thanked for providing office space, library and computing resources and logistic support.

## 13 Appendices

### 13.1 Appendix A: Excursus on the MLAP distribution

#### 13.1.1 Three bivariate Laplace distributions

For illustration, consider three bivariate Laplace distributions, all having null means but distinct scale matrices, as follows:

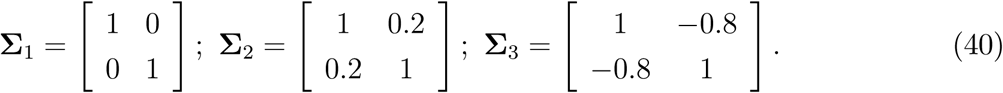

Using (7), the density under **Σ**_1_ is

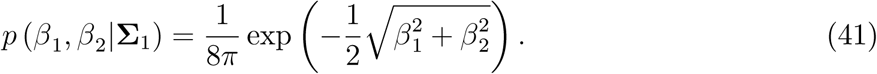

The covariance matrix here, **B**_1_ = 12**Σ**_1_, is diagonal, so the random variables are uncorrelated but not independent since (41) cannot be written as the product of two marginal densities. Under **Σ**_2_ and **Σ**_3_, the densities are

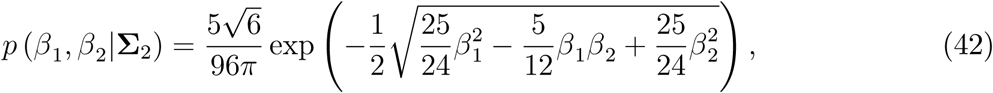

and

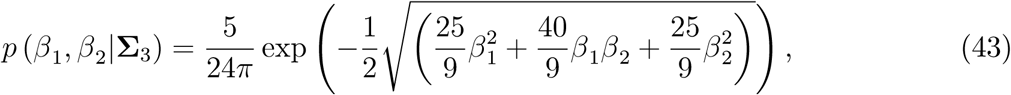

Five bivariate Laplace densities are shown in Supplementary Figure 1. (a) gives the density of the distribution of the two uncorrelated bivariate Laplace random variables (**Σ**_1_), and (b) and (c) show the positively (i.e., with **Σ**_2_) and negatively (with **Σ**_3_) correlated situations, respectively. These three densities have a sharp mode at *β*_1_ = *β*_2_ = 0 indicating that a bivariate Laplace prior would strongly shrink vectors to the (0, 0) point, acting similarly to the DE prior in the univariate Bayesian LASSO. (d) and (e) displays bivariate Laplace densities of distributions with non-null means.

#### 13.1.2 Conditional distributions

Dropping subscript *j* denoting a specific marker, partition the *T* × 1 vector of effects into 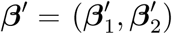 where the sub-vectors have orders *T*_1_ and *T*_2_, respectively; recall that *T* is the number of traits. Correspondingly, put

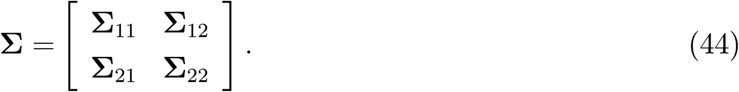

According to J. M. Marín (personal communication), the conditional distribution [***β*** _2_| ***β***_1_] has density

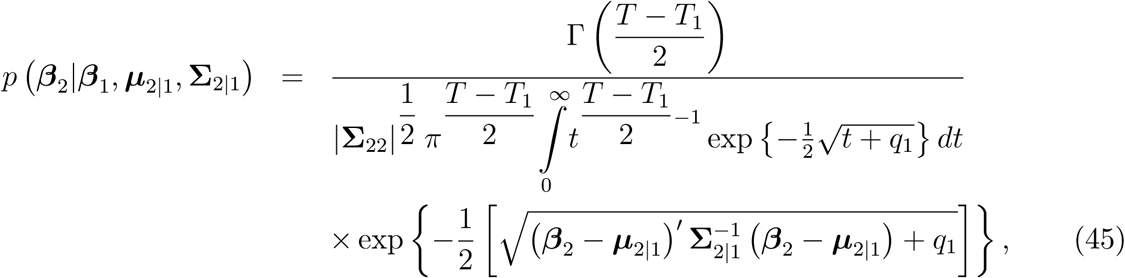

where 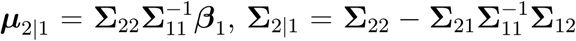 and 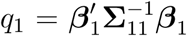. Similar to multivariate normal distribution, the conditional expectation is linear on the conditioning variable and **Σ**_2|1_ does not involve ***β***.

#### 13.1.3 Simulation of a multivariate Laplace distribution

Gómez et al. (2007) showed that *S* independent draws from a MLAP distribution with a null mean vector can be made as

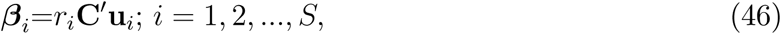

where **C**′ results from the Cholesky decomposition **Σ** = **C**′**C, u** is a *T* × 1 vector uniformly distributed on a *T* -dimensional unit sphere and *r* is a realization of a Gamma distribution with shape parameter *T* and scale 2. Vector **u** can be simulated by effecting *T* independent draws (*x*_*i*_; *i* = 1, 2, …, *T*) from a *N* (0, 1) distribution, and then forming the *t*^*th*^ element of **u** as 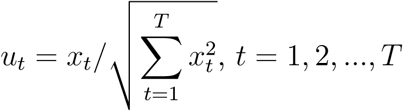.

Marginal distributions for the three bivariate Laplace distributions with scale matrices **Σ**_1_, **Σ**_2_ and **Σ**_3_ given above were estimated by sampling *S* = 300, 000 independent realizations; (46) was employed. Using the samples, zero-mean DE and normal distributions with the same variances were fitted, and the resulting densities were compared with the estimated densities based on the draws. As shown in Supplementary Figure 2, a normal distribution provided a poor approximation to the marginals from the three bivariate Laplace cases, and the sharp peak at 0, characteristic of a DE density, was not matched by such marginals. This is a corroboration of theoretical results in Gómez et al. (2007): marginals from MLAP distributions are elliptically contoured and not DE.

### 13.2 Appendix B: Mean vector of location parameters given ELSE

Consider (19). For *T* = 3, let

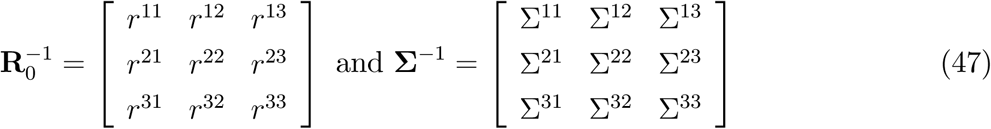

Expansion of the Kronecker products in (19) produces the system

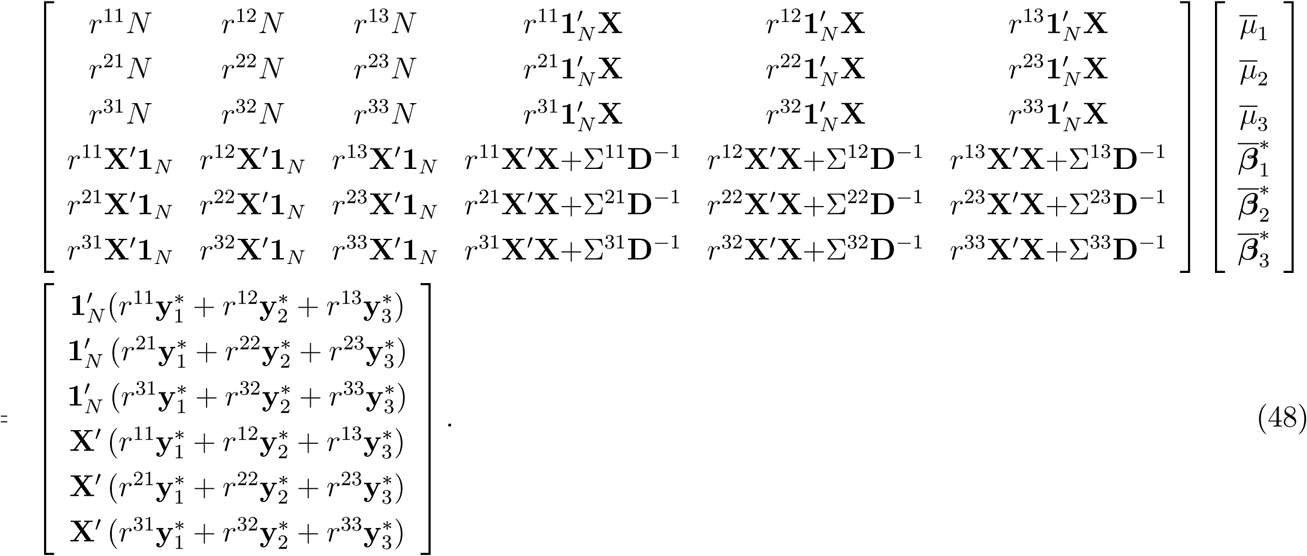

Observe how phenotypes for all traits contribute to the solutions of trait-specific effects.

### 13.3 Appendix C: Sampling from the conditional posterior distribution of 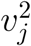

Consider (29). Let 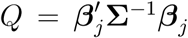 take values 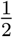, 1, 4, and 10, say. Numerical integration of (29) between 0 and 1000 produces 3.5203, 3.040 7, 1.844 3, 1.031 4 as reciprocal of the resulting integration constants, with the normalized densities shown in Supplementary Figure 3. The distributions are skewed, and as *Q* increases the density becomes flatter.

Let 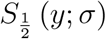 be the Lévy density of a positive random variable *Y* having a positive stable distribution with parameter *σ* (Samorodnitsky and Taqqu 2000). From Gómez et al. (2007) and Gómez-Sánchez-Manzano et al. (2008) the Lévy density is

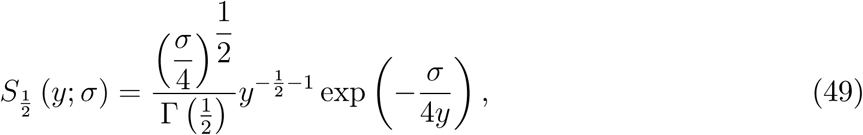

which is that of an inverse Gamma (*IG*) distribution with parameters 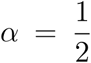 and 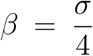 Consider now the transformation (Gómez et al. 2007) 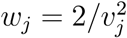 so using (29)

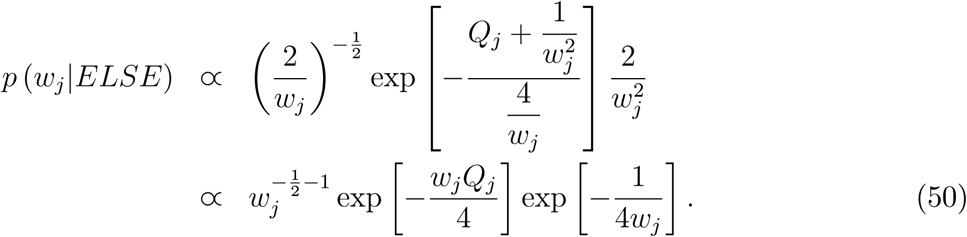

Consider a Metropolis-Hastings ratio *R* using (49) with *σ* = 1 as proposal distribution, and (18) let *y*_*j*_ be a proposed value and *w*_*j*_ be a member of the target distribution. The ratio is then

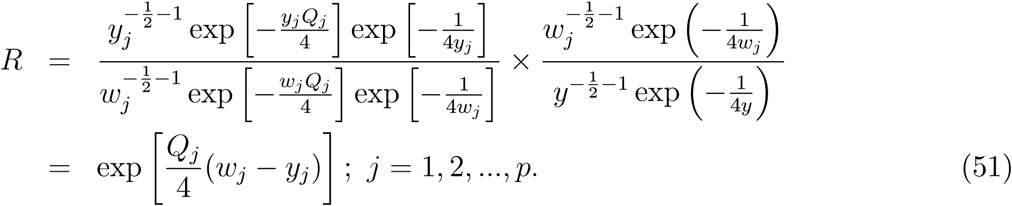

Hence if a proposal *y*_*j*_ is drawn from 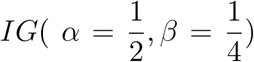, it can be accepted as belonging to the conditional posterior distribution of *w*_*j*_, with probability equal to *R* above. If accepted, a “new” 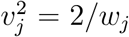 is a member of 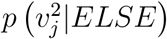 with probability *R* as well; otherwise stay with the current 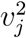.

### 13.4 Appendix D: Alternative algorithm for indirect sampling of marker effects

An alternative sampling scheme that uses an equivalent formulation of the model is presented; a two-trait (*T* = 2) situation is employed for ease of presentation. Let 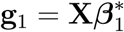 and 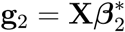 be the genomic values of the *N* individuals for each of the traits. A model could be

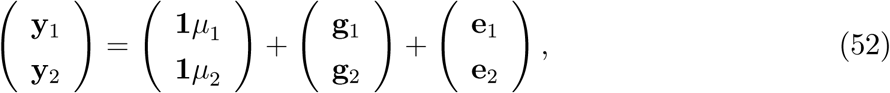

where residuals are as before. In a standard genomic best linear unbiased prediction (GBLUP, Van Raden 2008) setting, it is assumed that

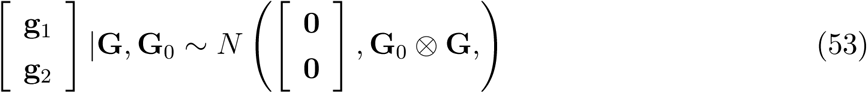

where **G** is an *N* × *N* marker-based matrix of “genomic relationships”, and

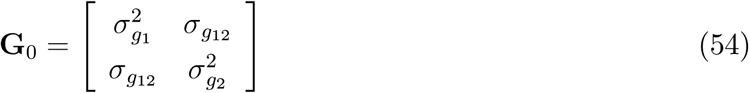

is a matrix containing the trait-specific genomic variances and their covariances. Specifically, from the definition of **g**_1_ and **g**_2_, and assuming that 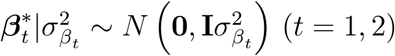

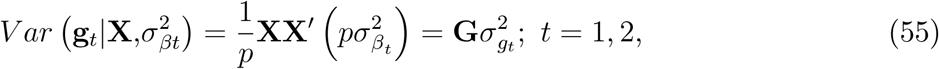

for **G** = **XX**′/*p* and 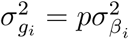. Similarly, 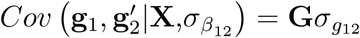, where 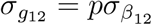 and 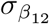 is the covariance between marker effects on traits 1 and 2. Let 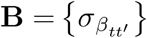 be the 2 × 2 variance-covariance matrix of marker effects

For a bivariate Bayesian LASSO model, conditionally on the *p* × 1 vector **v**^2^, one has

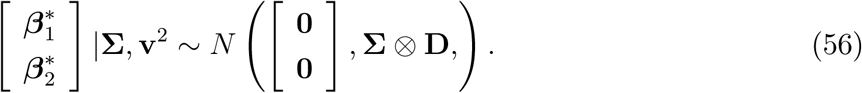

Hence

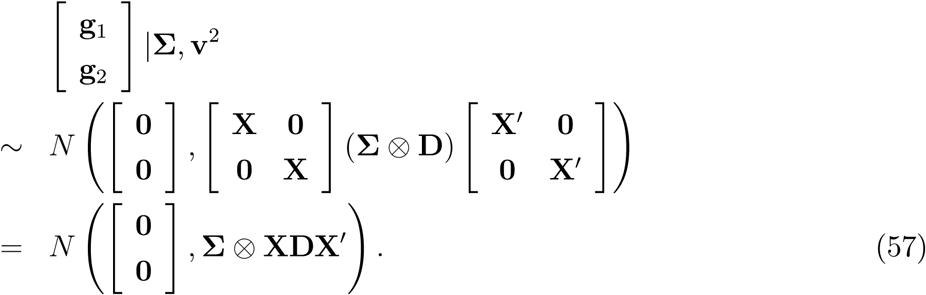

Let **C**_*Cond*_ = **Σ** ⊗ **XDX**′. Further,

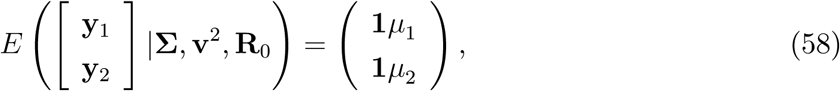

and

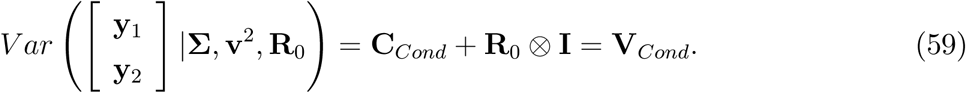

After assigning a flat prior to each of *µ*_1_ and *µ*_2_, standard results give that posterior distribution of the genotypic values given **Σ, v**^2^, **R**_0_ is normal, with mean vector

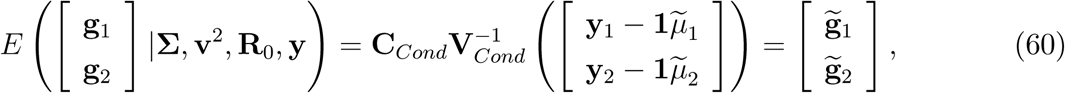

where

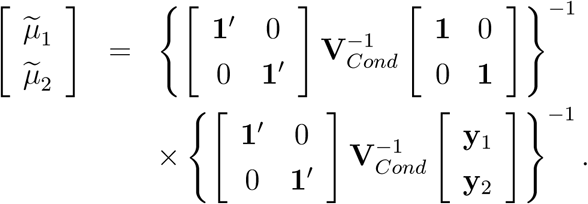

Further (Henderson 1975)

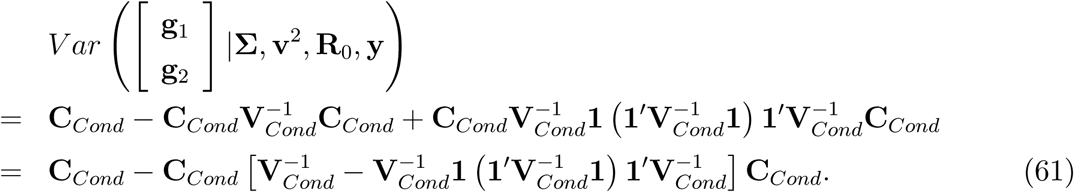

Hence, draws from the conditional posterior distribution of **g** = [**g**_1_ **g**_2_]′ given **Σ, v**^2^ and **R**_0_ can be obtained by sampling from a multivariate normal distribution with mean vector (60) and covariance matrix (61).

Assuming that, given **Σ, v**^2^ and **R**_0_, the vector 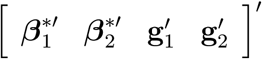 has a multivariate normal distribution, and let 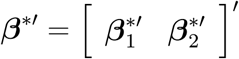. Hence

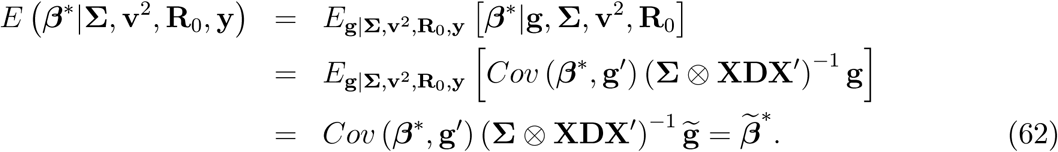

Now,

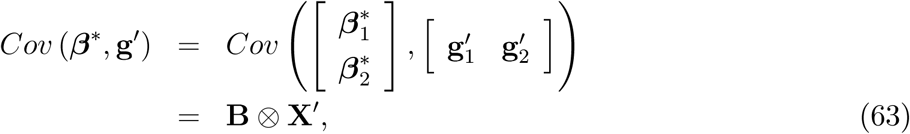

so

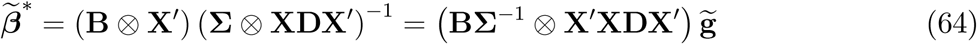

Similarly

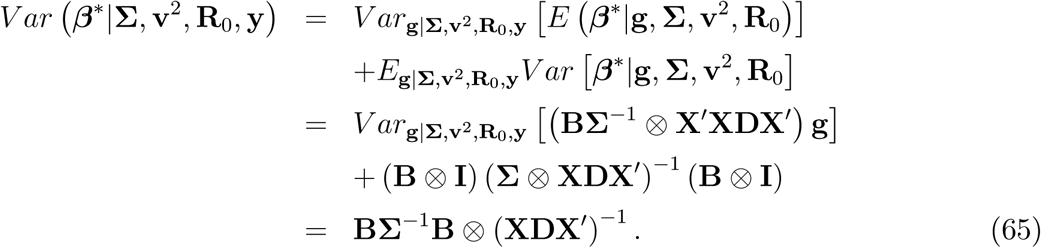

### 13.5 Appendix E: A conditional posterior mode approximation to marker effects

In spite of important advances in high-throughput computing, routine genetic evaluation of plants and animals is seldom done with MCMC methods. As an alternative to MCMC, we describe an iterative algorithm that produces point estimates of marker effects (and of linear functions thereof) and approximate measures of uncertainty in a computationally simpler manner. The algorithm uses a re-weighted set of linear “mixed model equations”, for which extremely efficient solvers exist. It is assumed that “good” estimates of **R**_0_ (the residual covariance matrix) and of **B** (the *T* × *T* variance-covariance matrix of markers effects) are available. From (8) **Σ** = **B**/ [4 (*T* + 1)], e.g., for *T* = 3 then **Σ** = **B**/16; hence, an assessment of the scale matrix of the MLAP distribution is easily available.

We make use of (2) and of (7) but employ the “markers within trait” representation given in (4). Letting ***θ*** = (***µ***′, ***β***′)′, the logarithm of the joint (conditionally on the dispersion matrices) posterior density of location effects, apart from a constant, is

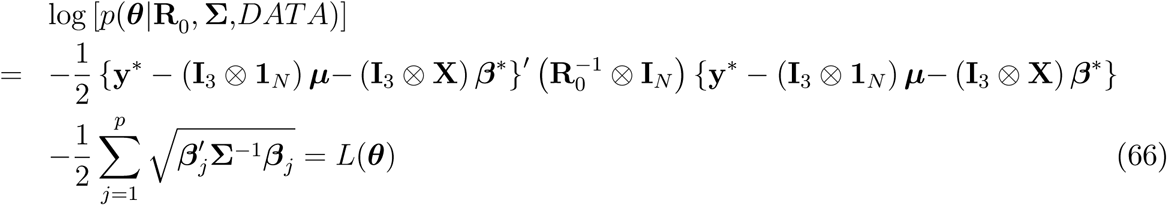

Let

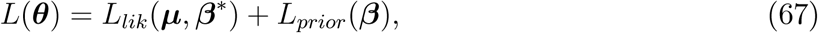

where *L*_*lik*_(***µ, β***\*) and *L*_*prior*_(***β***) are the two terms in (66). Then

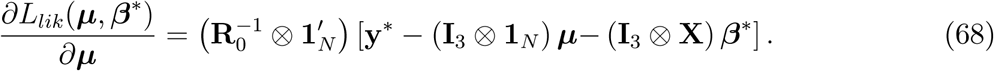

and

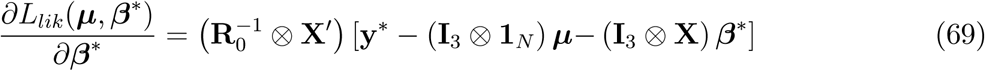

Observe now that the relationship between ***β*** and marker effects sorted within traits (***β***\*) can be expressed as ***β*** = **L*β***\* where **L** is a 3*p* × 3*p* non-singular matrix of elementary operators that rearrange rows and columns. For example, for *T* = 3 and *p* = 2 and with *β*_*jt*_ representing the effect of marker *j* on trait *t*,

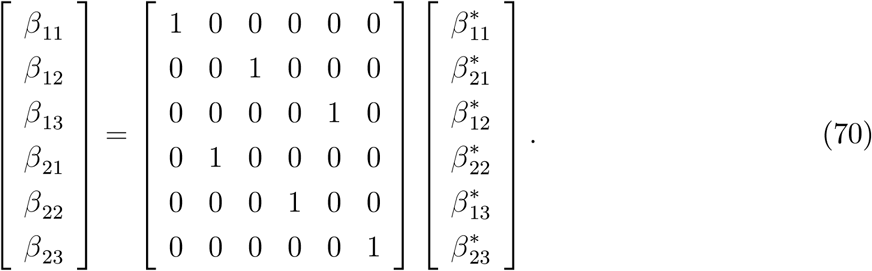

Since **L** is a matrix of elementary operators, **L**^−1^ = **L**′ (orthogonality) and ***β***\* = **L**′ ***β***; the absolute value of the Jacobian of the transformation from ***β*** to ***β***\* is equal to 1. The contribution of the prior to the gradient for marker effects is then

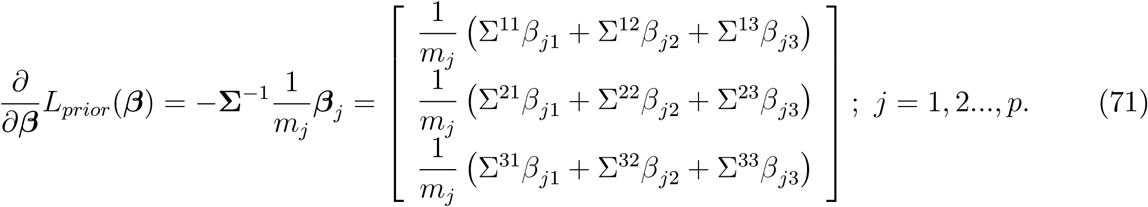

where 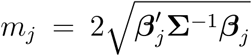 is proportional to the the Mahalanobis distance of ***β***_***j***_ away from (0, 0, 0) for *T* = 3. Hence, the 3*p* × 1 vector of derivatives with respect to all marker effects, sorted by traits within individuals is

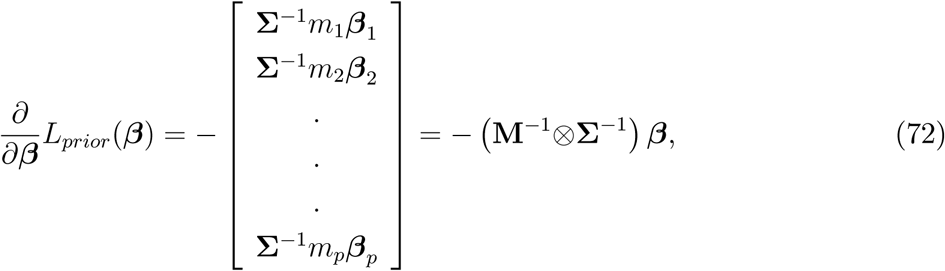

where **M** = *Diag* {*m*_*j*_} is a *p* × *p* diagonal matrix with typical element *m*_*j*_. Rearranging the differentials such that the sorting is by markers within traits

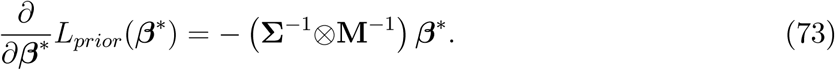

Collecting (69) and (73),

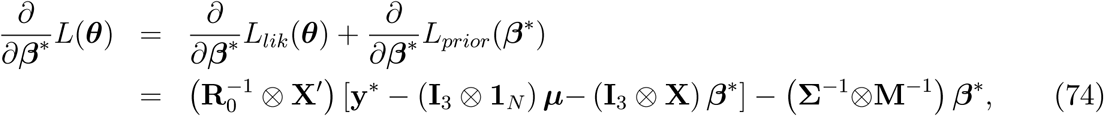

Setting (68) and (74) simultaneously to **0** produces the system of equations (not explicit)

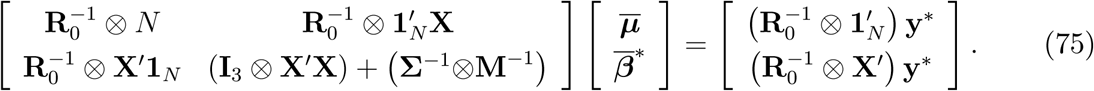

Expanding the equations above for *T* = 3 yields

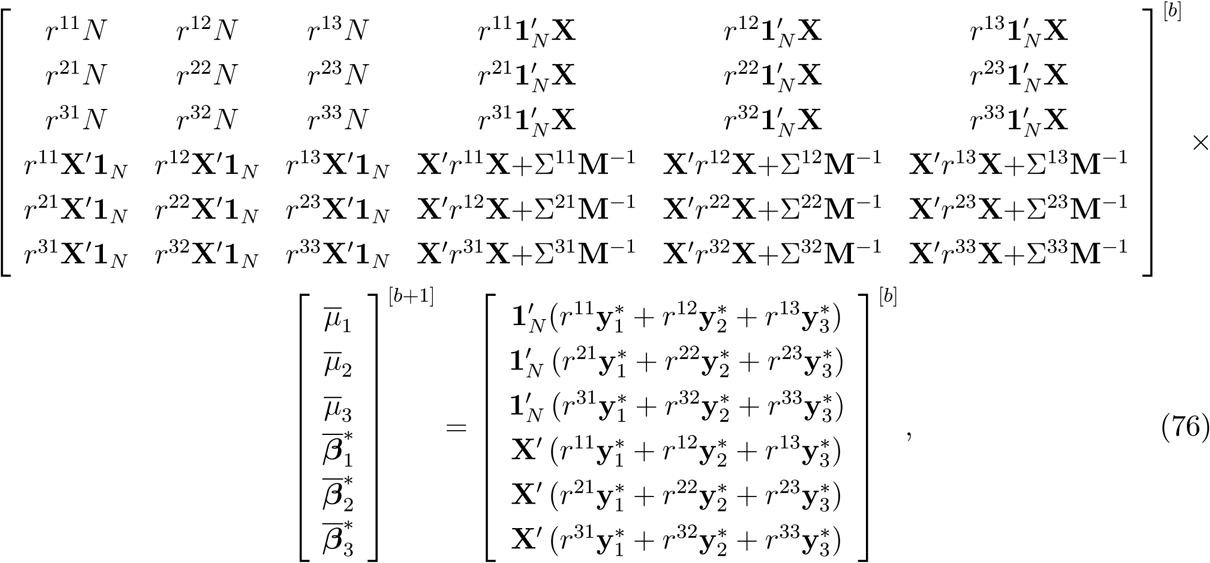

where *b* is iterate round. Matrix **M** =*Diag*(*m*_*j*_) changes at every round of iteration, so the system needs to be reconstituted repeatedly. Marker effects producing small values of the Mahalanobis distance away from 0 result in tiny *m*-values and, consequently, **M**^−1^ will have large diagonal elements. Hence, vectors of markers with weak effects are more strongly shrunk towards the 0 coordinate than those having strong effects in at least one trait

The variance-covariance matrix of the conditional posterior distribution can be approximated as

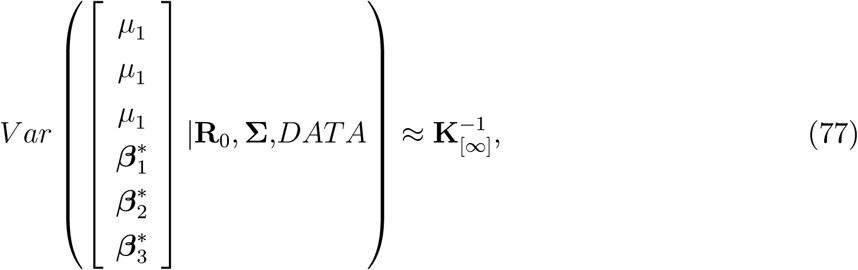

with ∞ indicating parameters evaluated at converged values, assuming that convergence has been attained at a hopefully global mode.

### 13.6 Appendix F: Treatment of missing data

Let a multi-trait data point (*T* × 1) on individual *i* be represented as

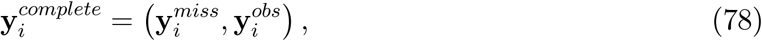

where “miss” denotes a missing record, e.g., if *T* = 2, a record could be missing for trait 1 or for trait 2; 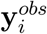 represents the phenotypes for the traits observed in individual *i*. The posterior predictive distribution of 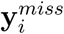 has density

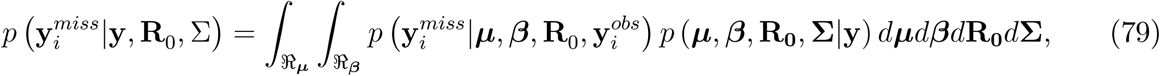

provided that data points in *i* are conditionally (given ***µ***, ***β***, **R**_0_) independent of any other *i*′ in the sample, and with **y** being all observed data. The preceding formulae implies that 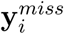 can be imputed by sampling ***µ***, ***β***, **R**_0_, **Σ** from their posterior distribution and then drawing from

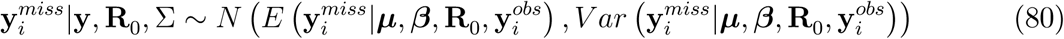

Since the sampling model is normal, for *T* = 3 one has

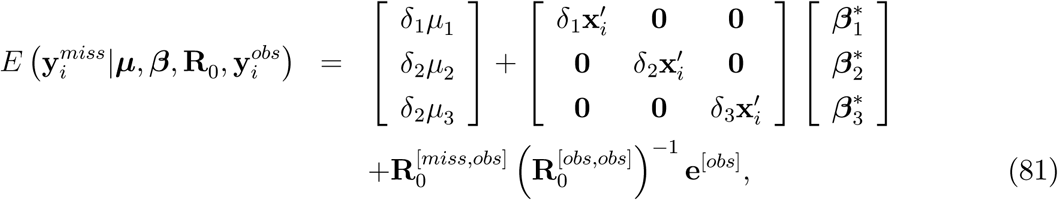

where *δ*_1_, *δ*_2_, *δ*_3_ take the value 1 when a given trait is missing in case *i*, or denote “exclude from formula” otherwise; 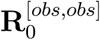 is the part of **R**_0_ pertaining to observed phenotypes for case *i*, and 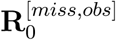 is the submatrix of residual covariances between missing and observed traits. Further,

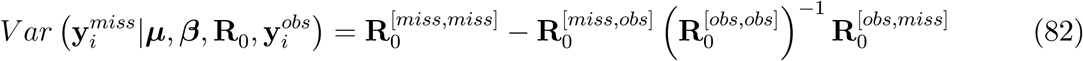

For example, let *T* = 3 and suppose that trait 1 is missing in case 250 but that traits 2 and 3 have been recorded; here

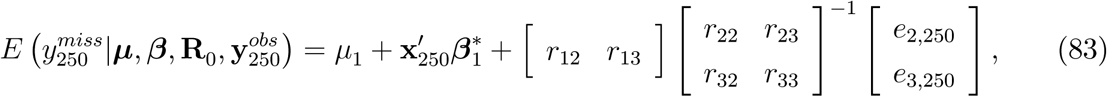

and

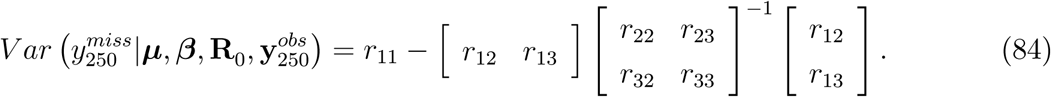

In the MCMC algorithm, missing data are sampled independently across cases, but dependently within case by addressing the pattern of missingness peculiar to each observation. Samples for missing observations can be used to estimate predictive distributions for the missing data (Gelfand et al. 1992; Sorensen and Gianola 2002; Gelman et al. 2014).

### 14 Legend for Supplemental Figures

Figure S1. Five bivariate Laplace densities.

Figure S2. Double exponential versus marginal (from bivariate Laplace) versus normal densities

Figure S3. Normalized densities of mixing variable in MCMC algorithm

Figure S4. Shrinkage factor: mean trait 1

Figure S5. Shrinkage factor: mean trait 2

Figure S6. Shrinkage factor: marker 10, trait 1

Figure S7. Shrinkage factor: marker 200, trait 2

Figure S8. Shrinkage factor: R0[1,1]

Figure S9. Shrinkage factor: R0[1,2]

Figure S10. Shrinkage factor: R0[2,2]

Figure S11. Shrinkage factor: SIGMA[1,1]

Figure S12. Shrinkage factor: SIGMA[1,2]

Figure S13. Shrinkage factor: SIGMA[2,2]

Figure S14. Trace plots of R0[1,1], R0[1,2], R0[2,2]

Figure S15. Trace plots of SIGMA[1,1], SIGMA[1,2], SIGMA[2,2]

Figure S16. Path to convergence in MAP-MBL (maximum a posteriori-multiple trait Bayesian LASSO)

Figure S17. BLUP of marker effects versus MBL posterior means and MAP-MBL solutions Figure S18. Fitted genetic values: BLUP, MBL and MAP-MBL

