## Supplementary material for "A multiple-trait Bayesian Lasso for genome-enabled analysis and prediction of complex traits": SUPPLMENTAL FIGURES

Figure S1. Five bivariate Laplace densities.
Figure S2. Double exponential versus marginal (from bivariate Laplace) versus normal densities
Figure S3. Normalized densities of mixing variable in MCMC algorithm
Figure S4. Shrinkage factor: mean trait 1
Figure S5. Shrinkage factor: mean trait 2
Figure S6. Shrinkage factor: marker 10, trait 1
Figure S7. Shrinkage factor: marker 200, trait 2
Figure S8. Shrinkage factor: R0[1,1]
Figure S9. Shrinkage factor: R0[1,2]
Figure S10. Shrinkage factor: R0[2,2]
Figure S11. Shrinkage factor: SIGMA[1,1]
Figure S12. Shrinkage factor: SIGMA[1,2]
Figure S13. Shrinkage factor: SIGMA[2,2]
Figure S14. Trace plots of R0[1,1], R0[1,2], R0[2,2]
Figure S15. Trace plots of SIGMA[1,1], SIGMA[1,2], SIGMA[2,2]
Figure S16. Path to convergence in MAP-MBL (maximum a posteriori-multiple trait Bayesian LASSO)
Figure S17. BLUP of marker effects versus MBL posterior means and MAP-MBL solutions
Figure S18. Fitted genetic values: BLUP, MBL and MAP-MBL

a) Density of two uncorrelated bivariate Laplace random variables with null means and unit scales. e1 and e2 are coordinates of bivariate vectors.

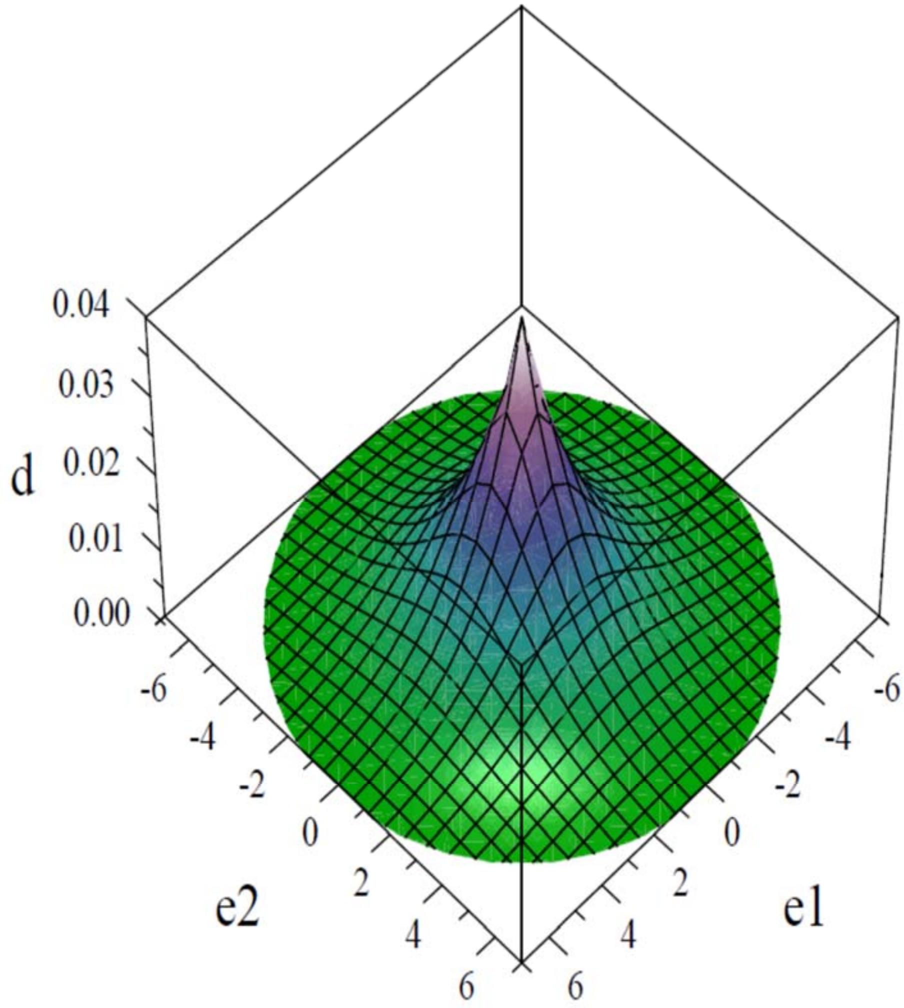

b) Density of two positively correlated bivariate Laplace random variables with null means and unit scales. e1 and e2 are coordinates of bivariate vectors

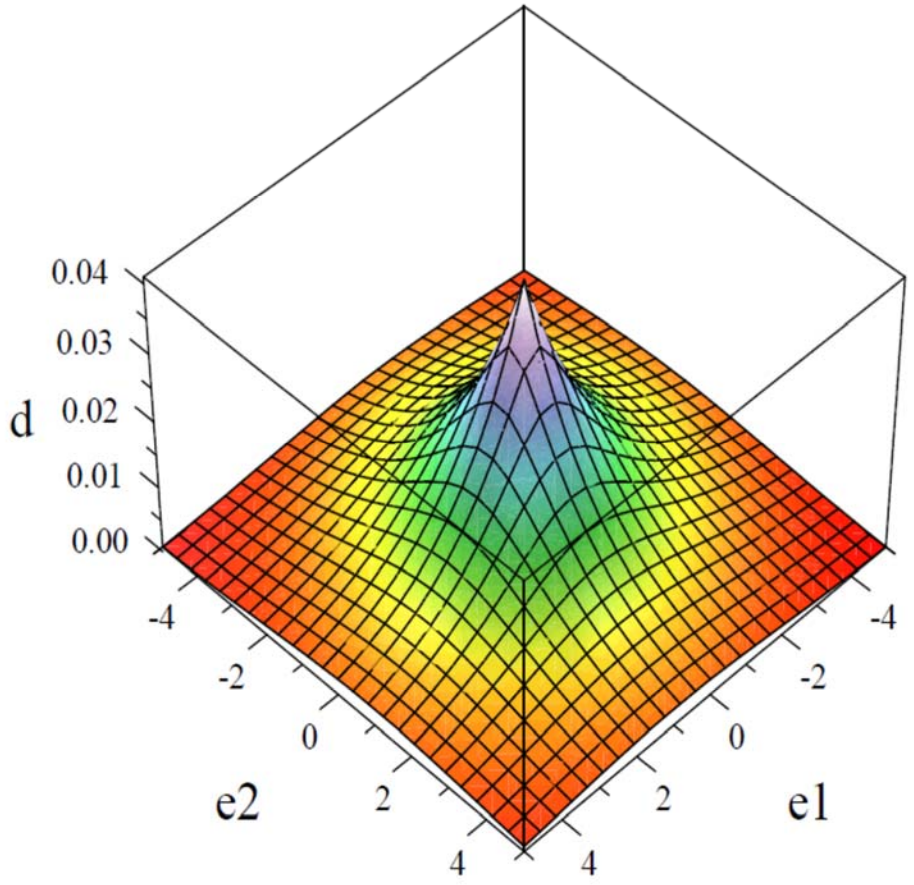

c) Density of two negatively correlated bivariate Laplace random variables with null means and unit scales. e1 and e2 are coordinates of bivariate vectors

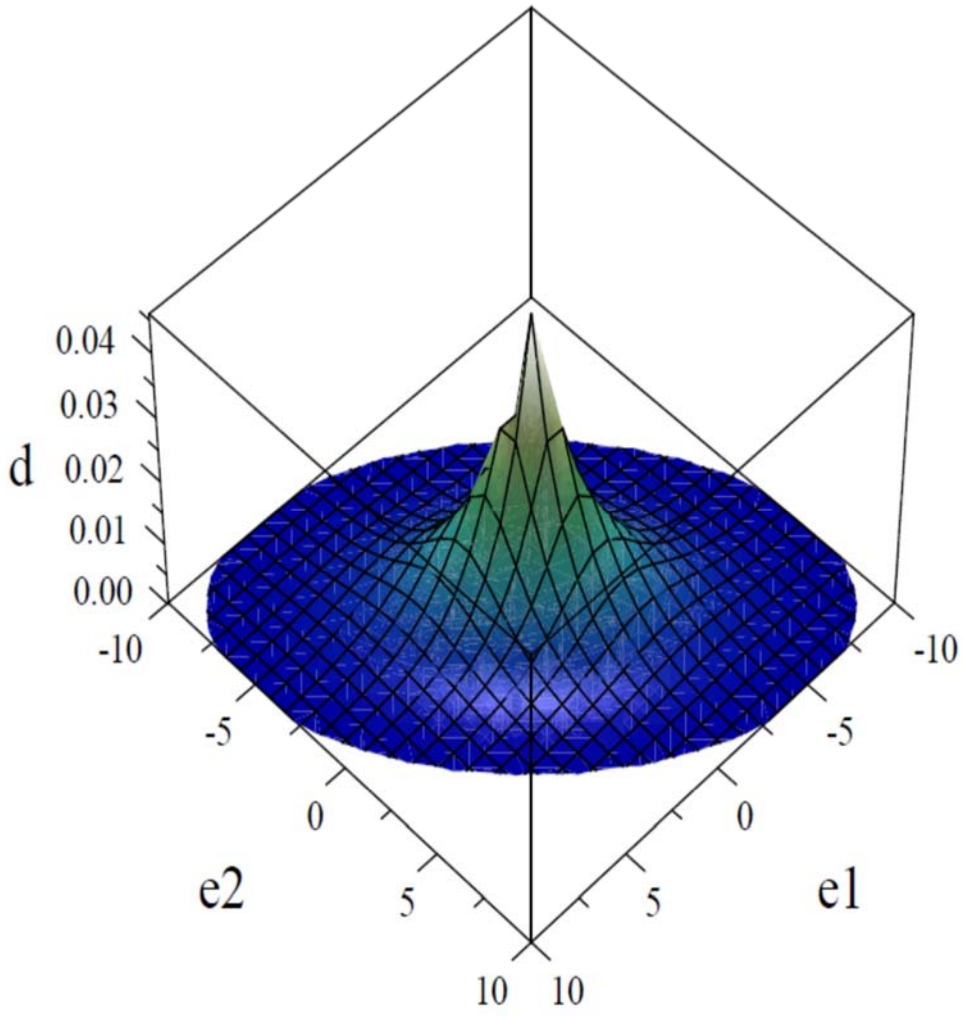

d) Bivariate Laplace density with means (scales) -4 (1) and 4 (4); correlation= 0. C1 and C2 are coordinates of bivariate vectors.

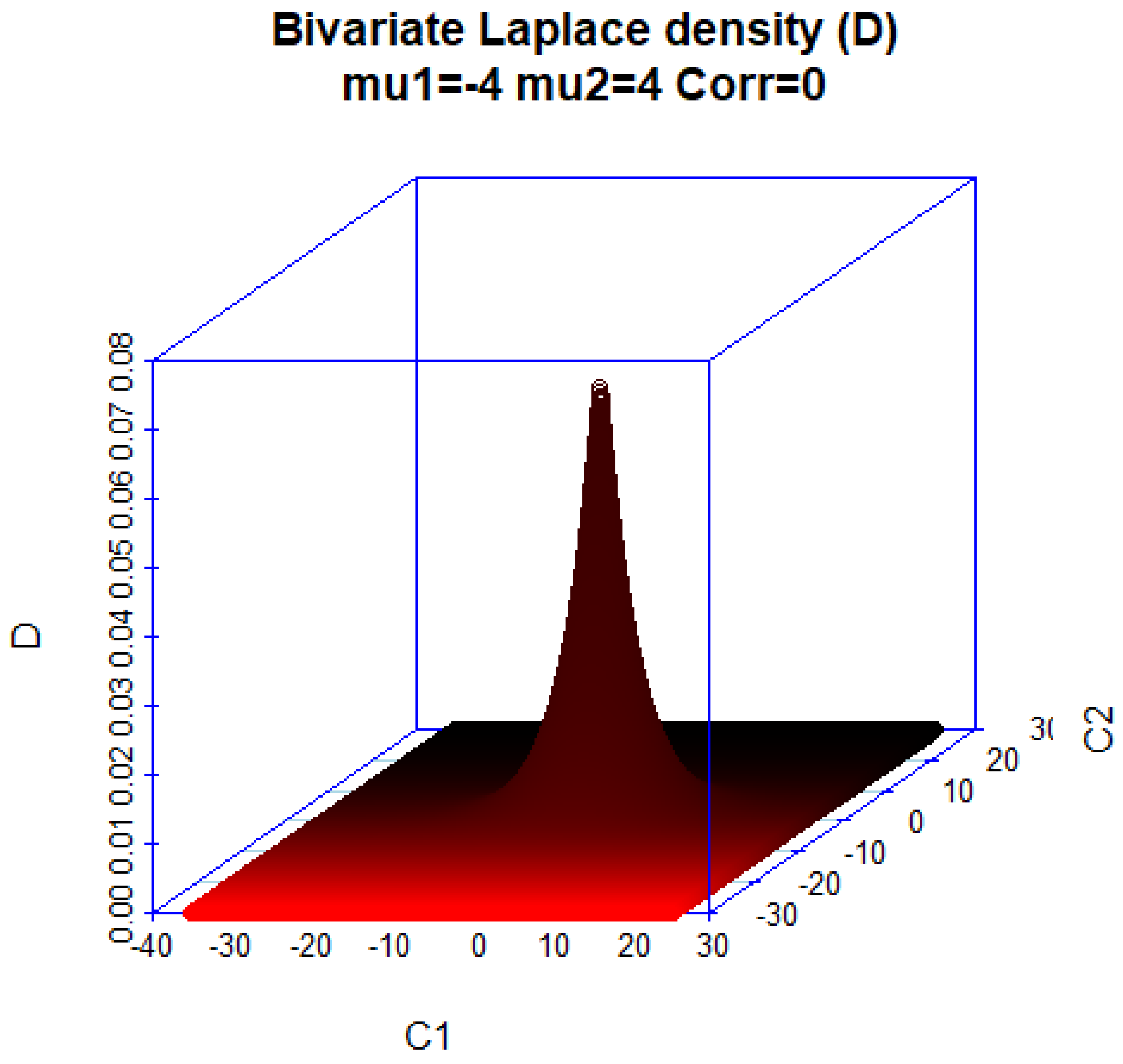


e) Bivariate Laplace density with means (scales) -4 (1) and 4 (4); correlation=- 0.75. C1 and C2 are coordinates of bivariate vectors.

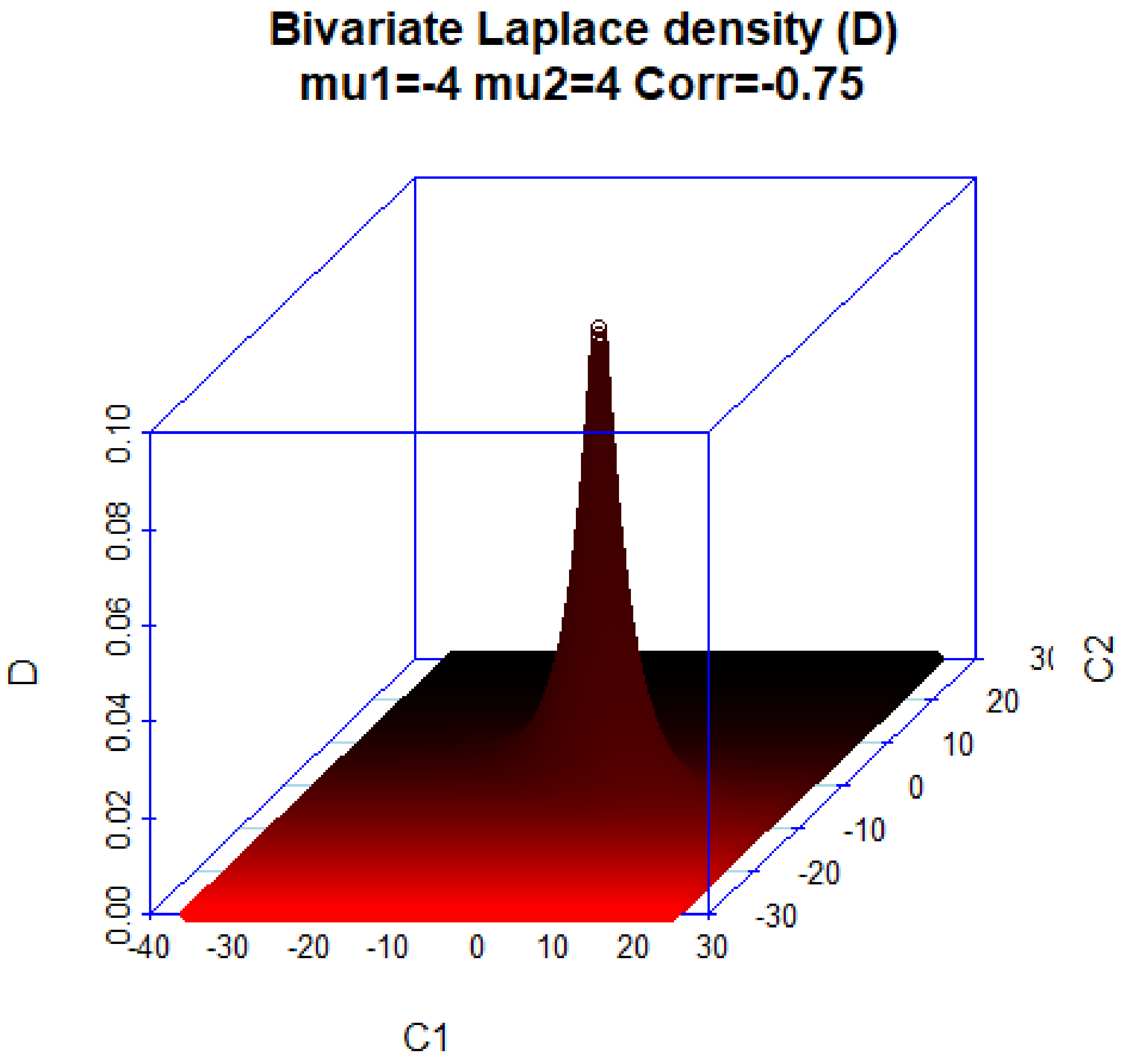
Bivariate Laplace density (D)
mu1=-4 mu2=4 Corr=-0.75
D
0.00
0.02
0.04
0.06
0.08
0.10
C1
-40
-30
-20
-10
0
10
20
30
C2
-30
-20
-10
0
10
20

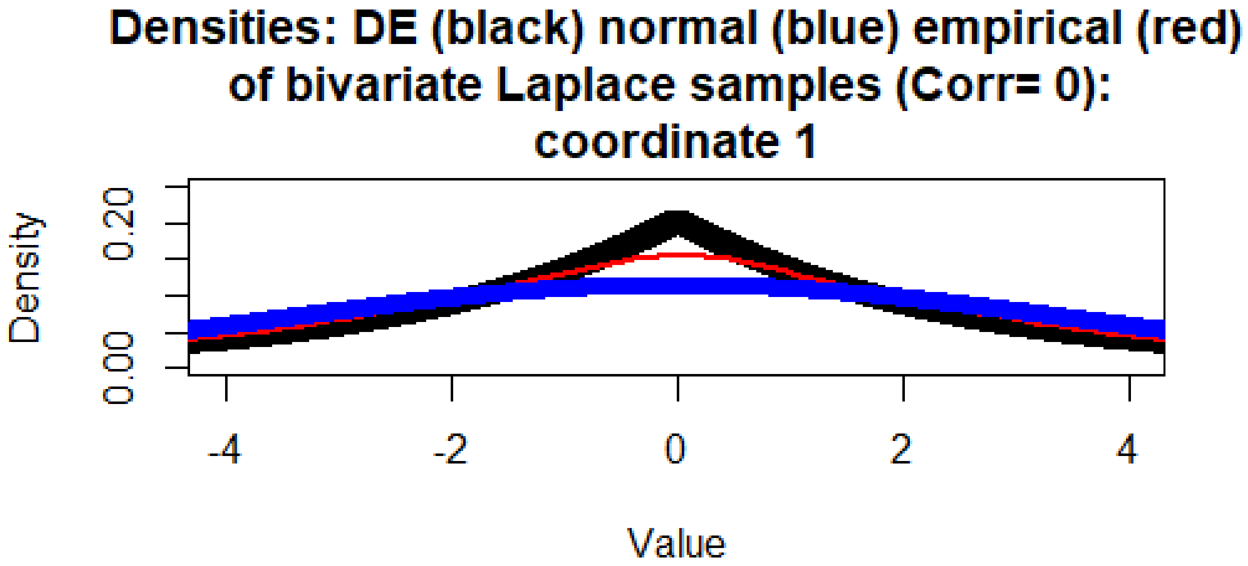
Densities: DE (black) normal (blue) empirical (red)
of bivariate Laplace samples (Corr= 0):
coordinate 1
Density
0.00
0.20
-4
-2
0
2
4
Value

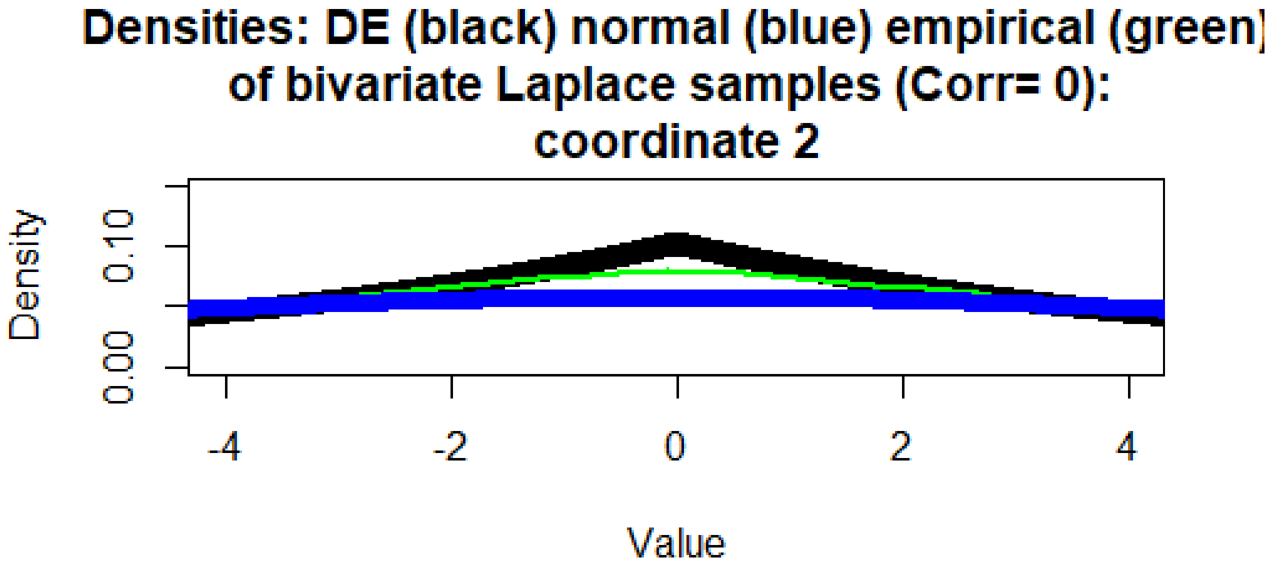
Densities: DE (black) normal (blue) empirical (green)
of bivariate Laplace samples (Corr= 0):
coordinate 2
Density
0.00
0.10
-4
-2
0
2
4
Value

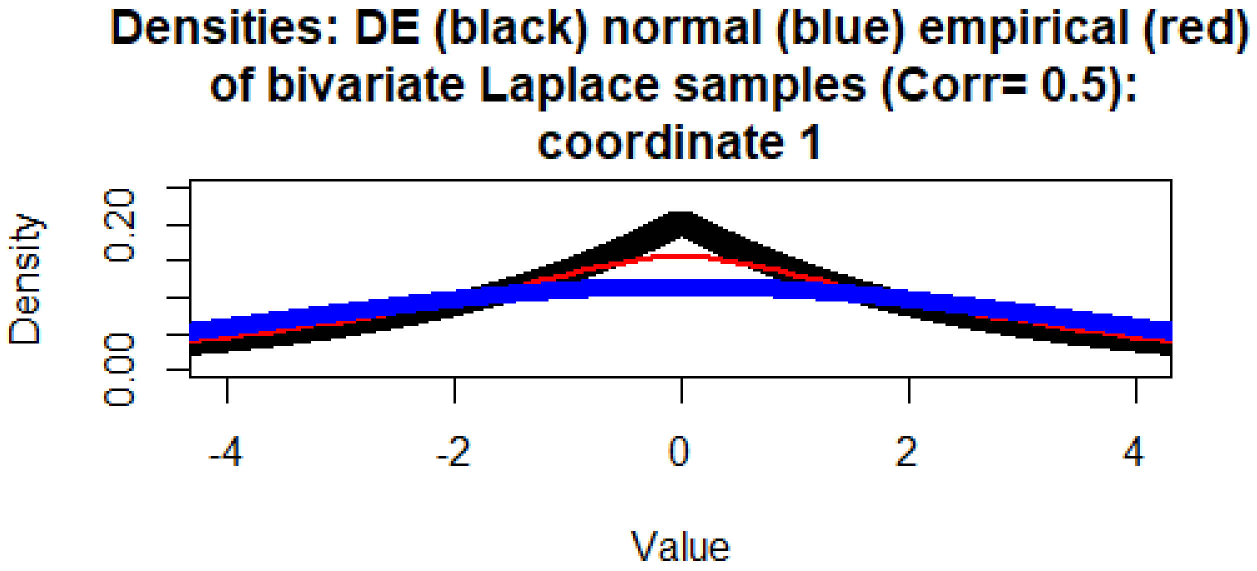
Densities: DE (black) normal (blue) empirical (red)
of bivariate Laplace samples (Corr= 0.5):
coordinate 1
Density
0.00
0.20
-4
-2
0
2
4
Value

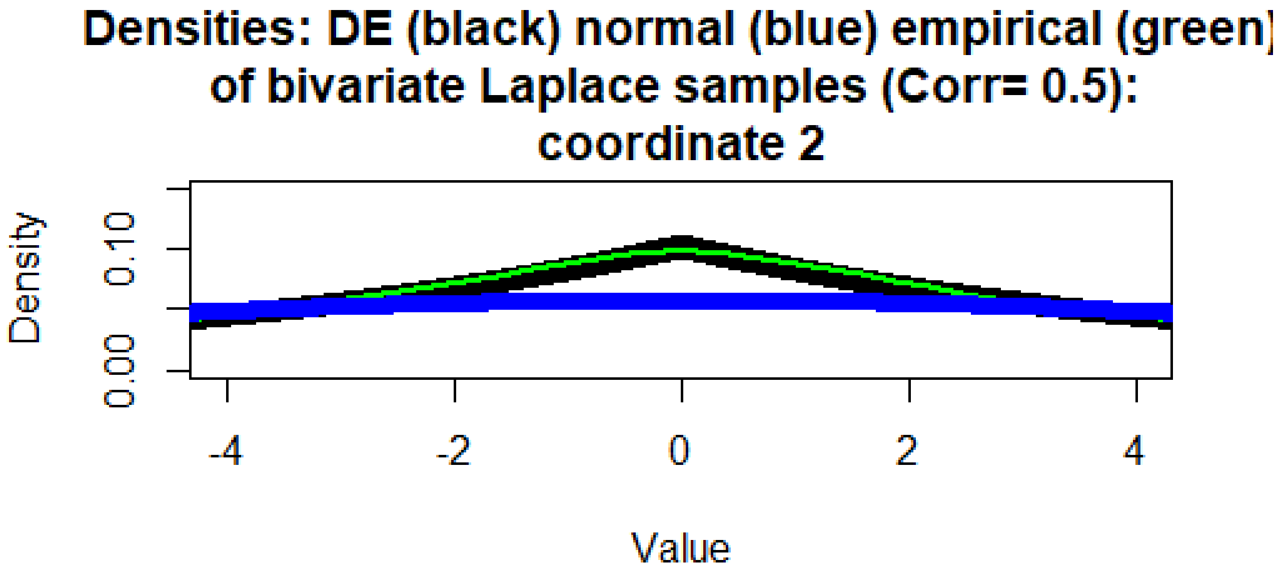
Densities: DE (black) normal (blue) empirical (green)
of bivariate Laplace samples (Corr= 0.5):
coordinate 2
Density
0.00
0.10
-4
-2
0
2
4
Value

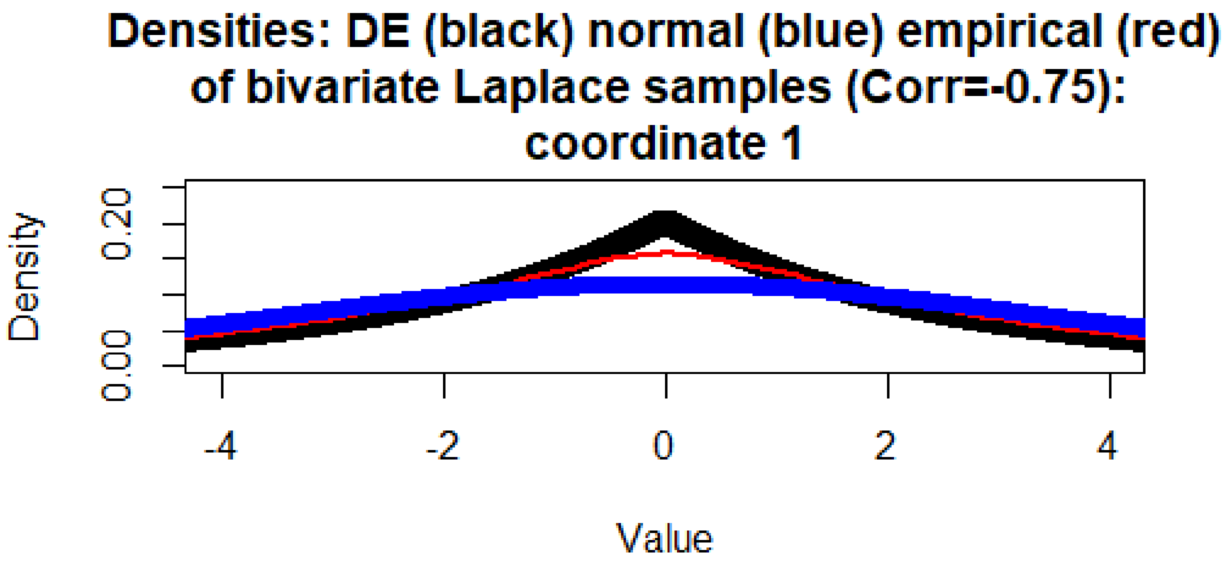
Densities: DE (black) normal (blue) empirical (red)
of bivariate Laplace samples (Corr=-0.75):
coordinate 1
Density
0.00
0.20
-4
-2
0
2
4
Value

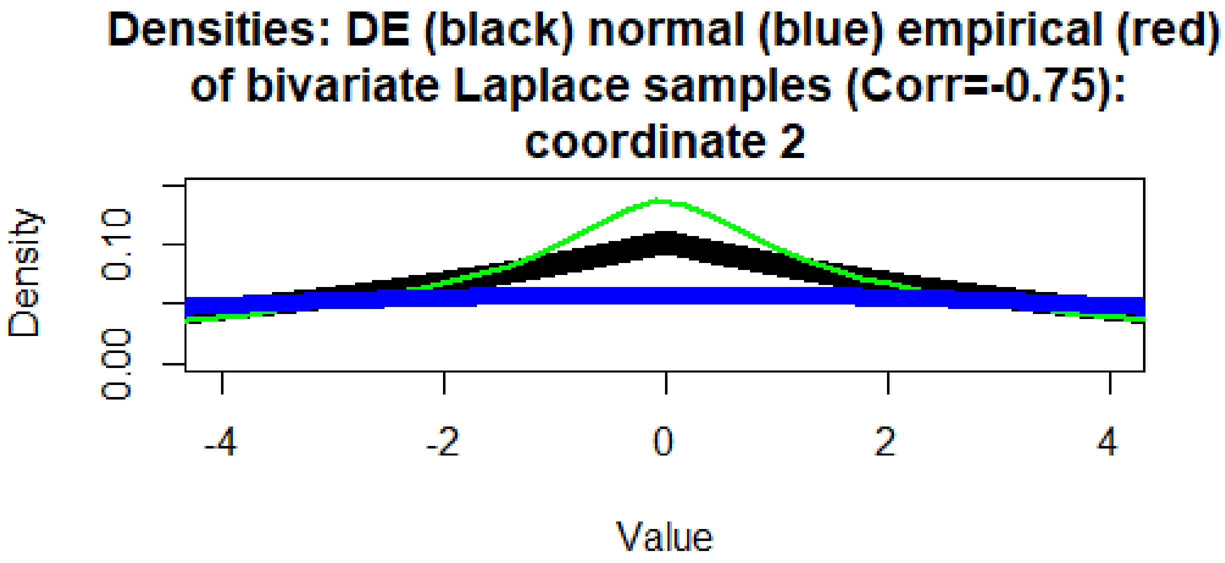
Densities: DE (black) normal (blue) empirical (red)
of bivariate Laplace samples (Corr=-0.75):
coordinate 2
Density
0.00
0.10
-4
-2
0
2
4
Value

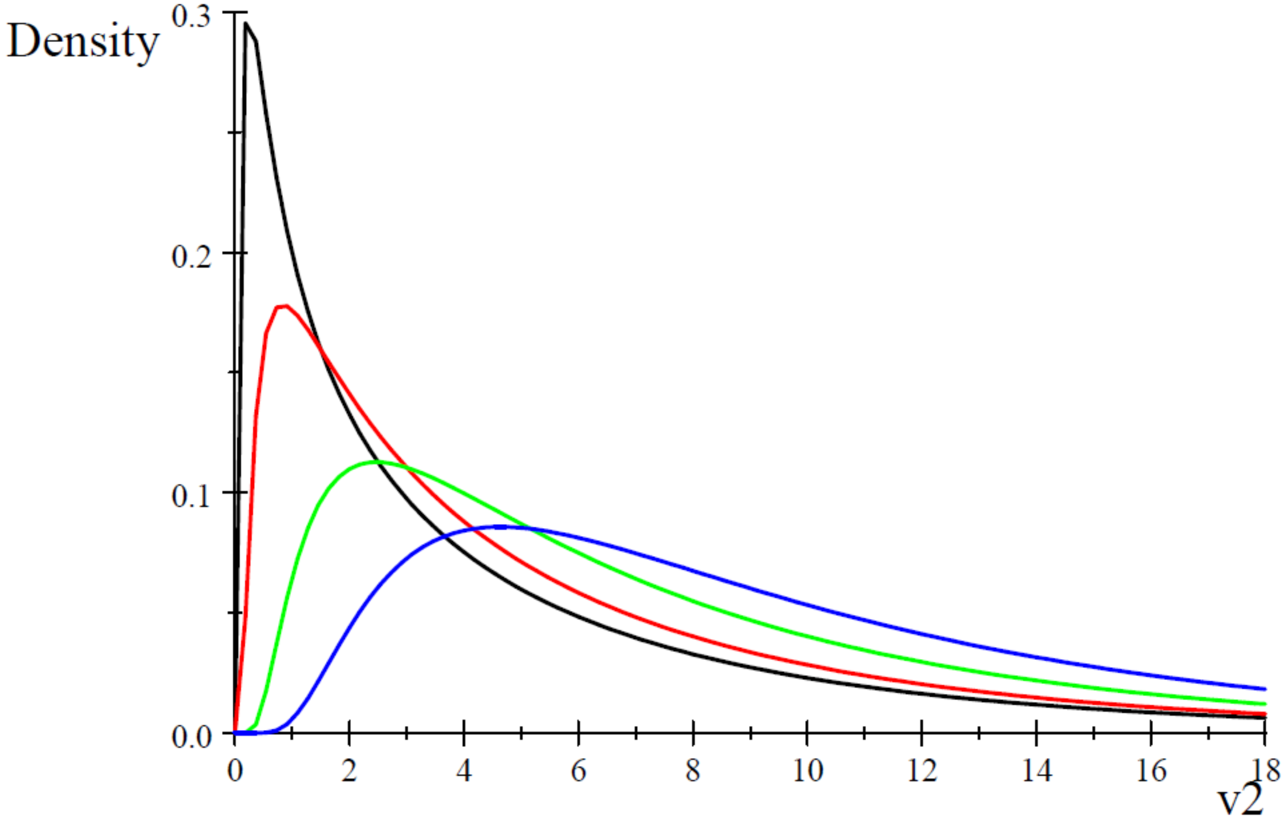


Figure 3. Normalized densities of conditional posterior distributions of the mixing variable $v^2$ at $Q = \frac{1}{2}$ (Black); 1 (Red); 4 (Green), and 10 (Blue).

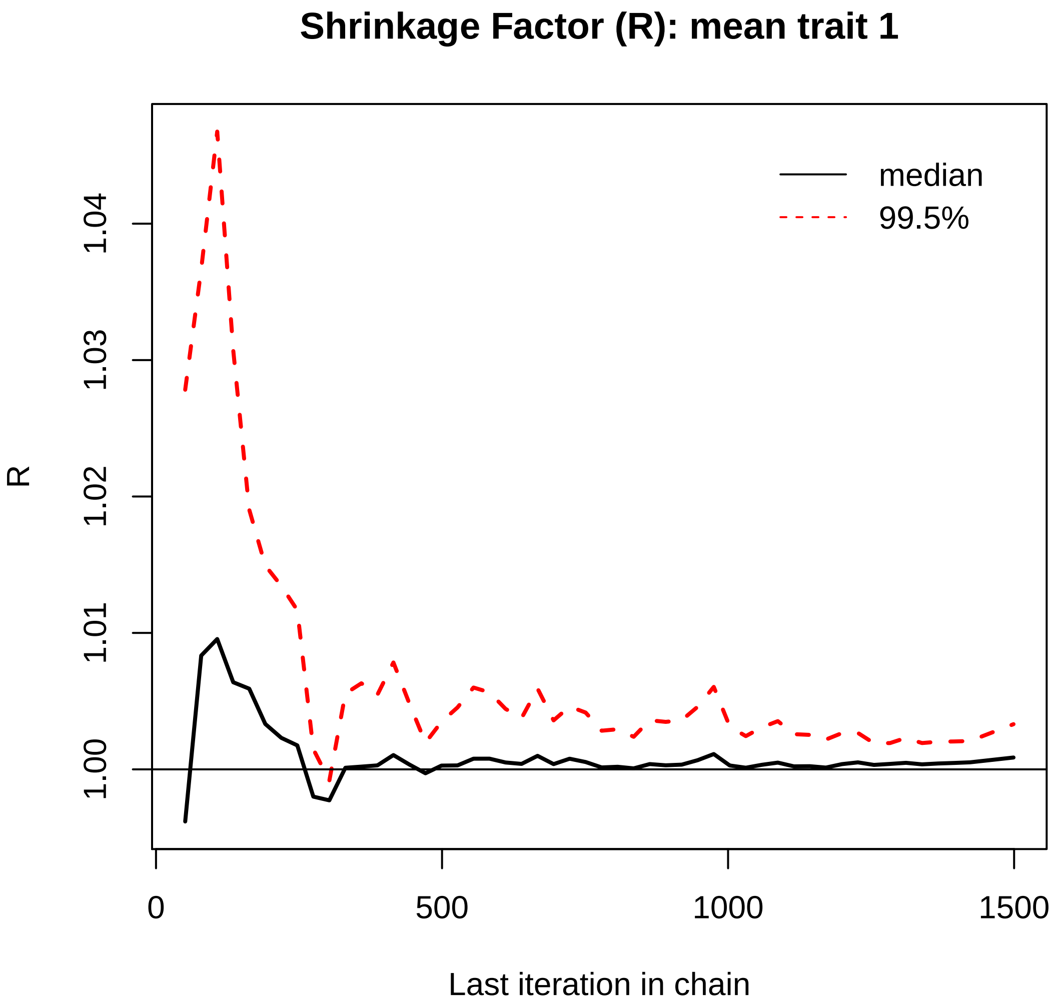
Shrinkage Factor (R): mean trait 1
median
99.5%
R
1.00
1.01
1.02
1.03
1.04
0
500
1000
1500
Last iteration in chain

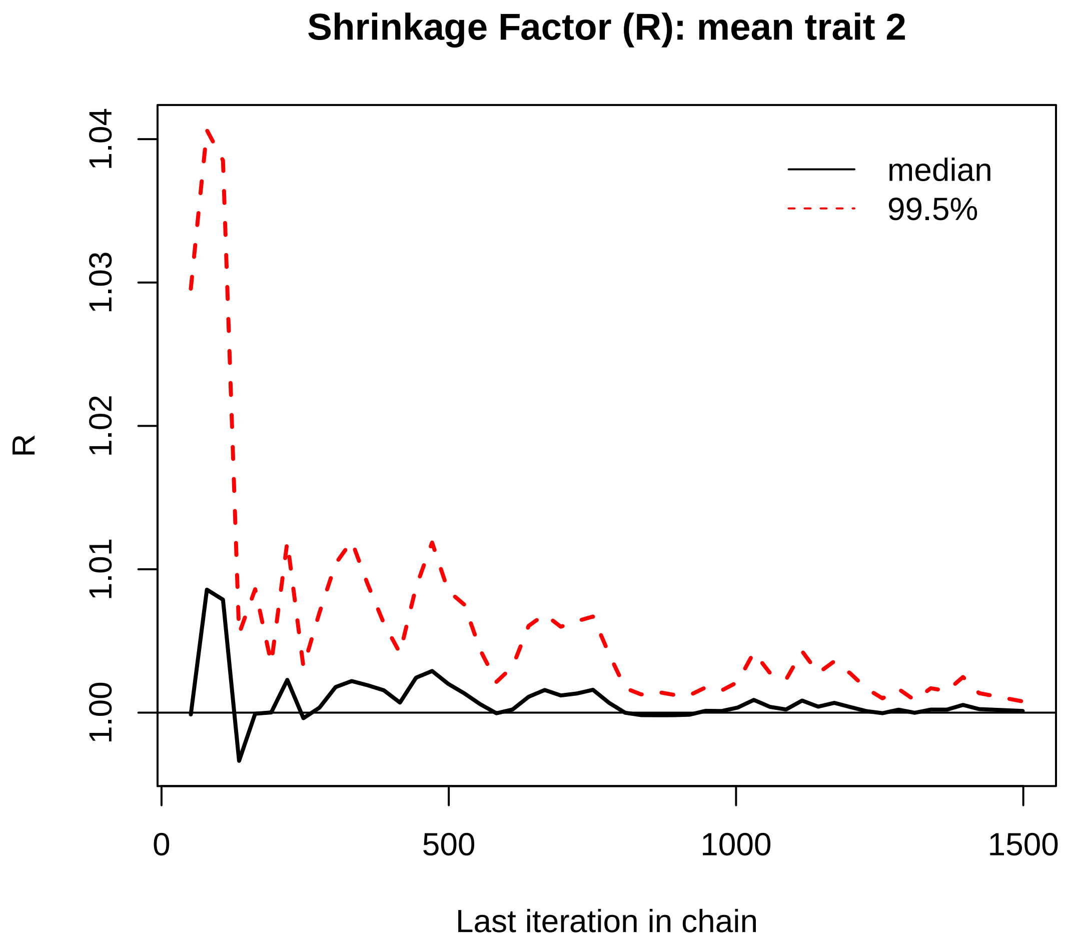
Shrinkage Factor (R): mean trait 2
R
1.04
1.03
1.02
1.01
1.00
median
99.5%
0
500
1000
1500
Last iteration in chain

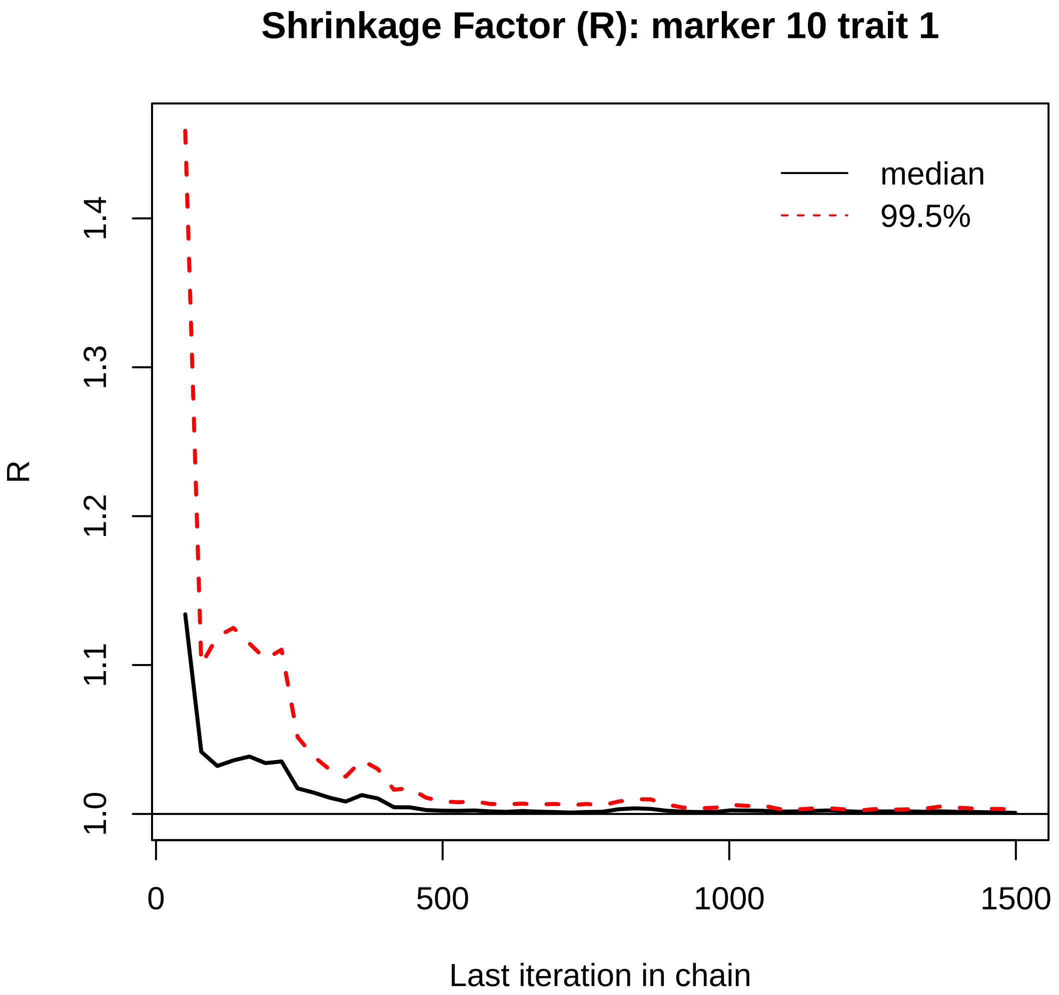
Shrinkage Factor (R): marker 10 trait 1
median
99.5%
R
1.0
1.1
1.2
1.3
1.4
0
500
1000
1500
Last iteration in chain

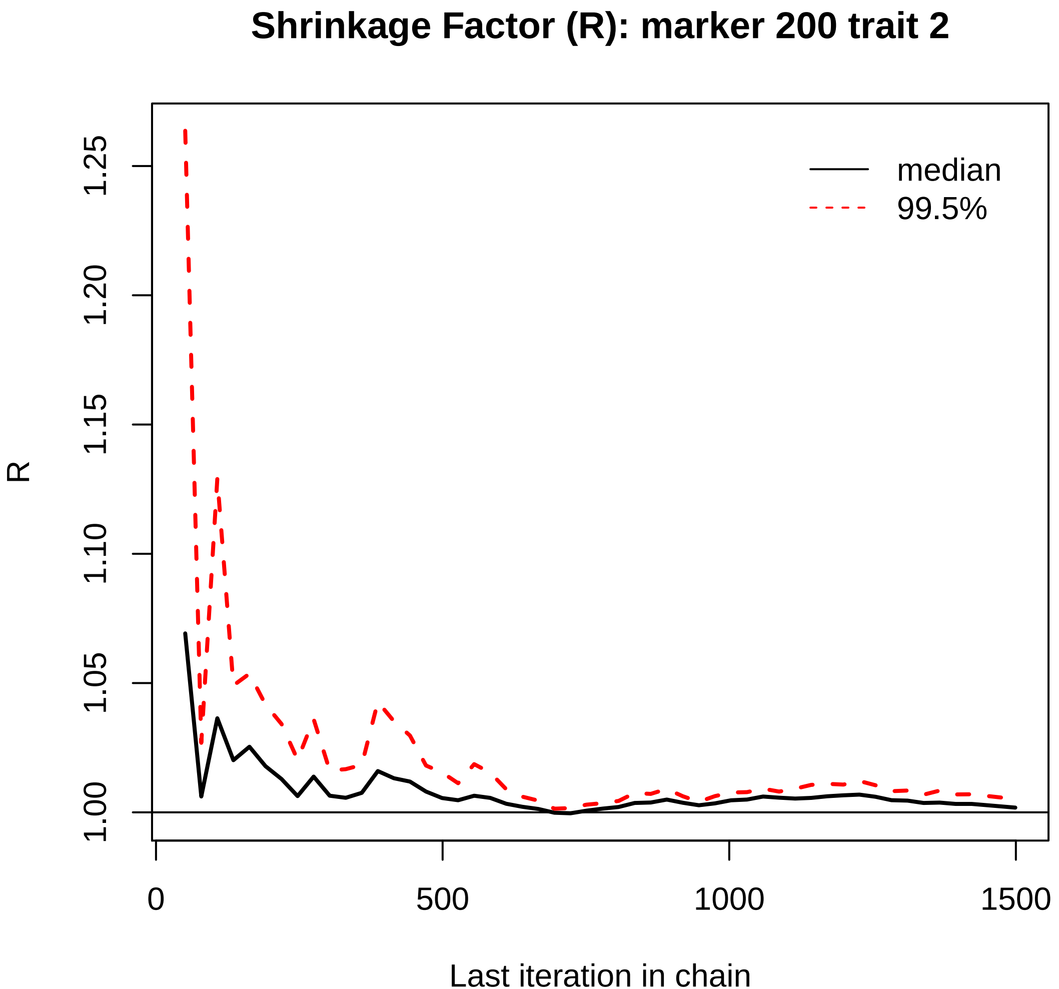

Shrinkage Factor (R): marker 200 trait 2
R
1.00
1.05
1.10
1.15
1.20
1.25
0
500
1000
1500
Last iteration in chain
median
99.5%

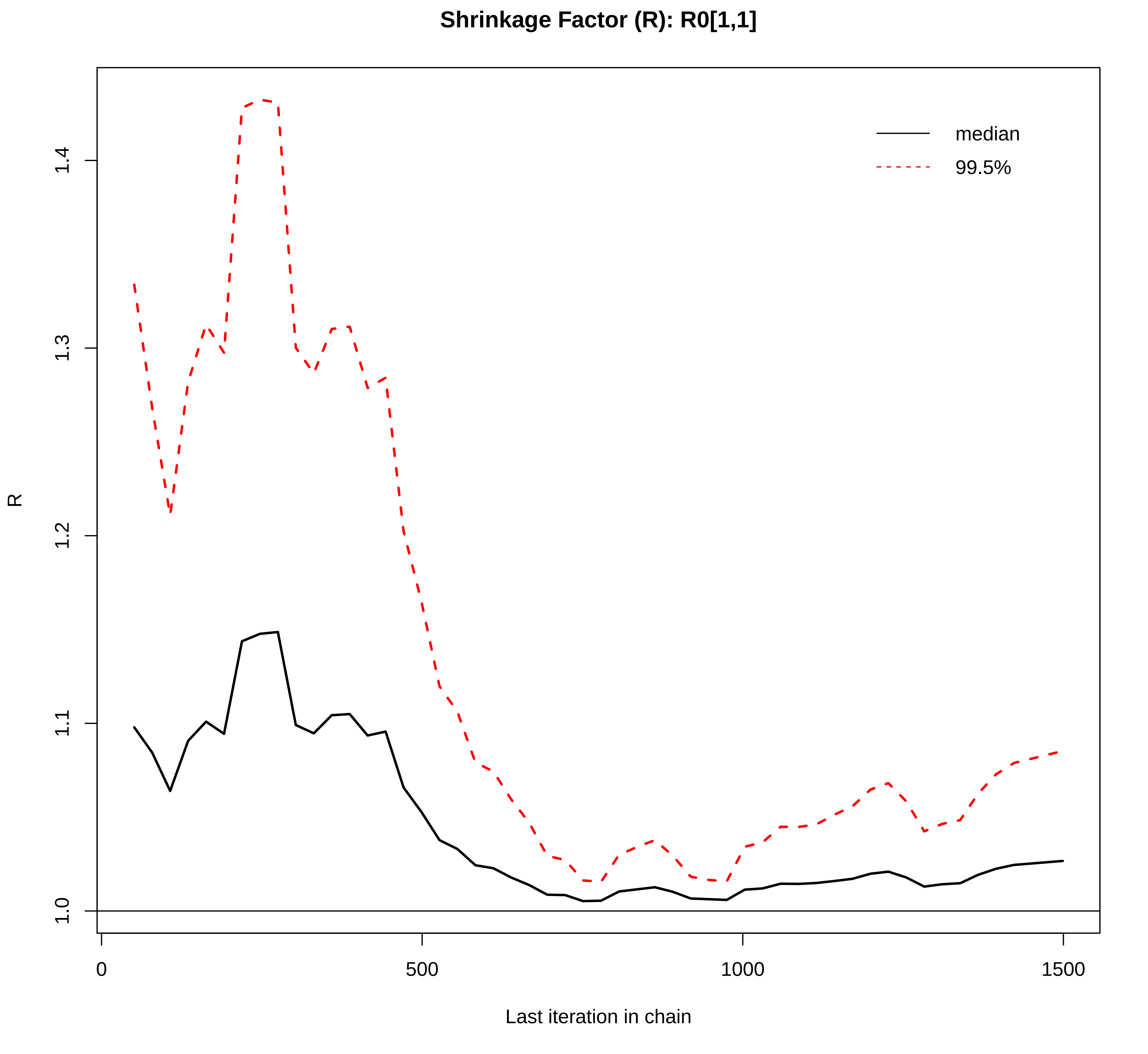

Shrinkage Factor (R): R0[1,1]
median
99.5%
R
1.0
1.1
1.2
1.3
1.4
0
500
1000
1500
Last iteration in chain

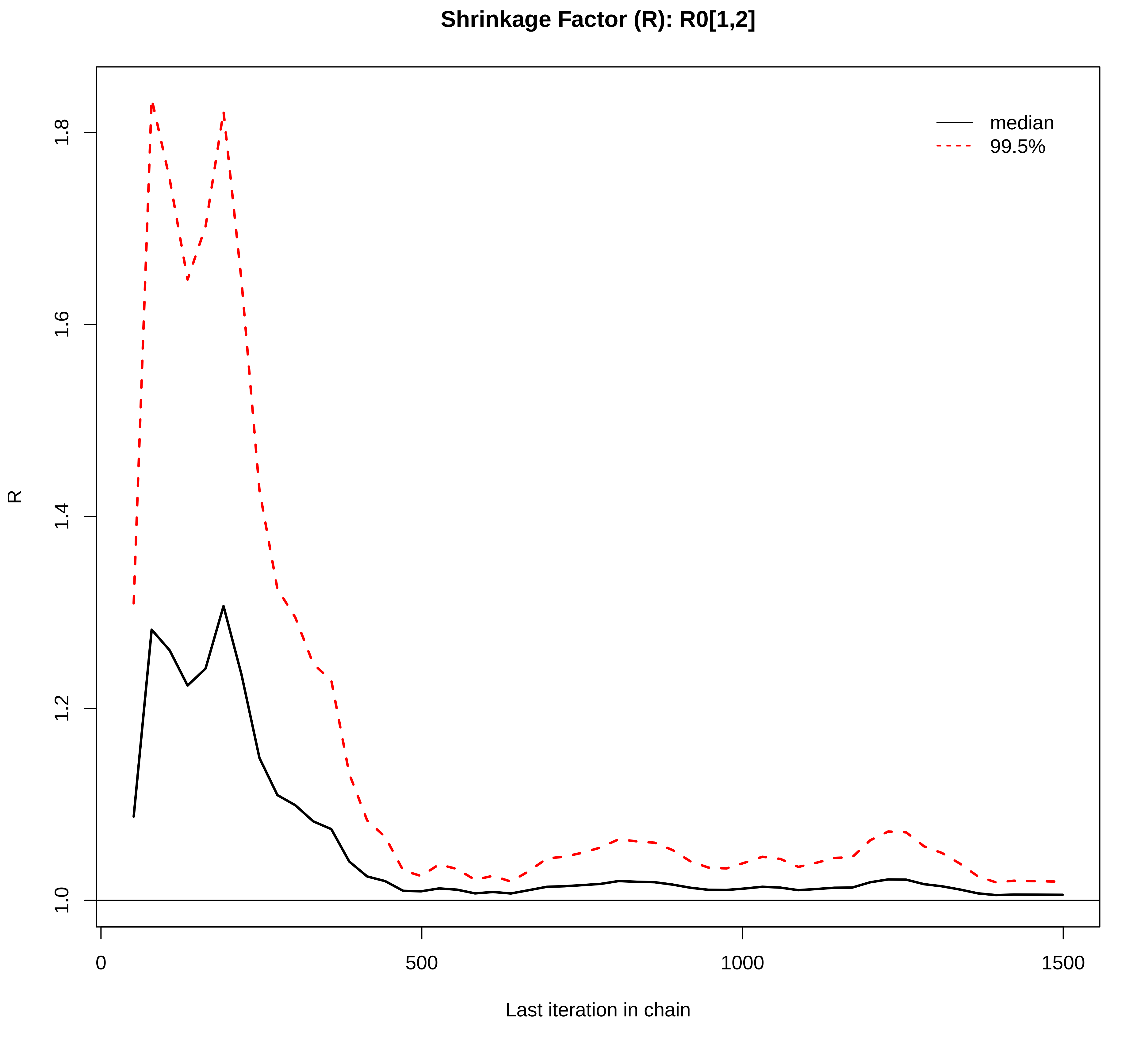
Shrinkage Factor (R): R0[1,2]
median
99.5%
R
1.0
1.2
1.4
1.6
1.8
0
500
1000
1500
Last iteration in chain

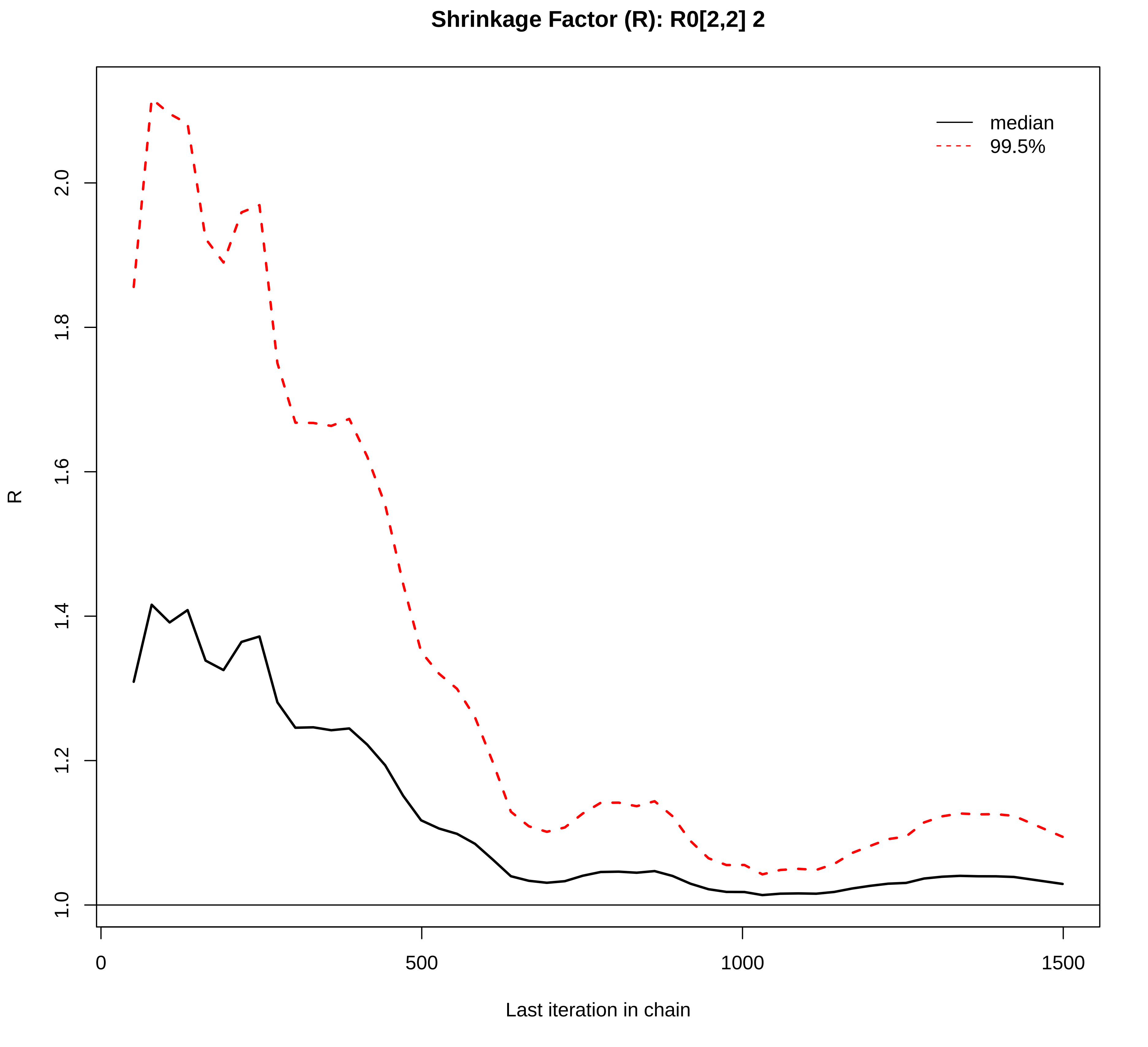

Shrinkage Factor (R): R0[2,2] 2
median
99.5%
R
2.0
1.8
1.6
1.4
1.2
1.0
0
500
1000
1500
Last iteration in chain

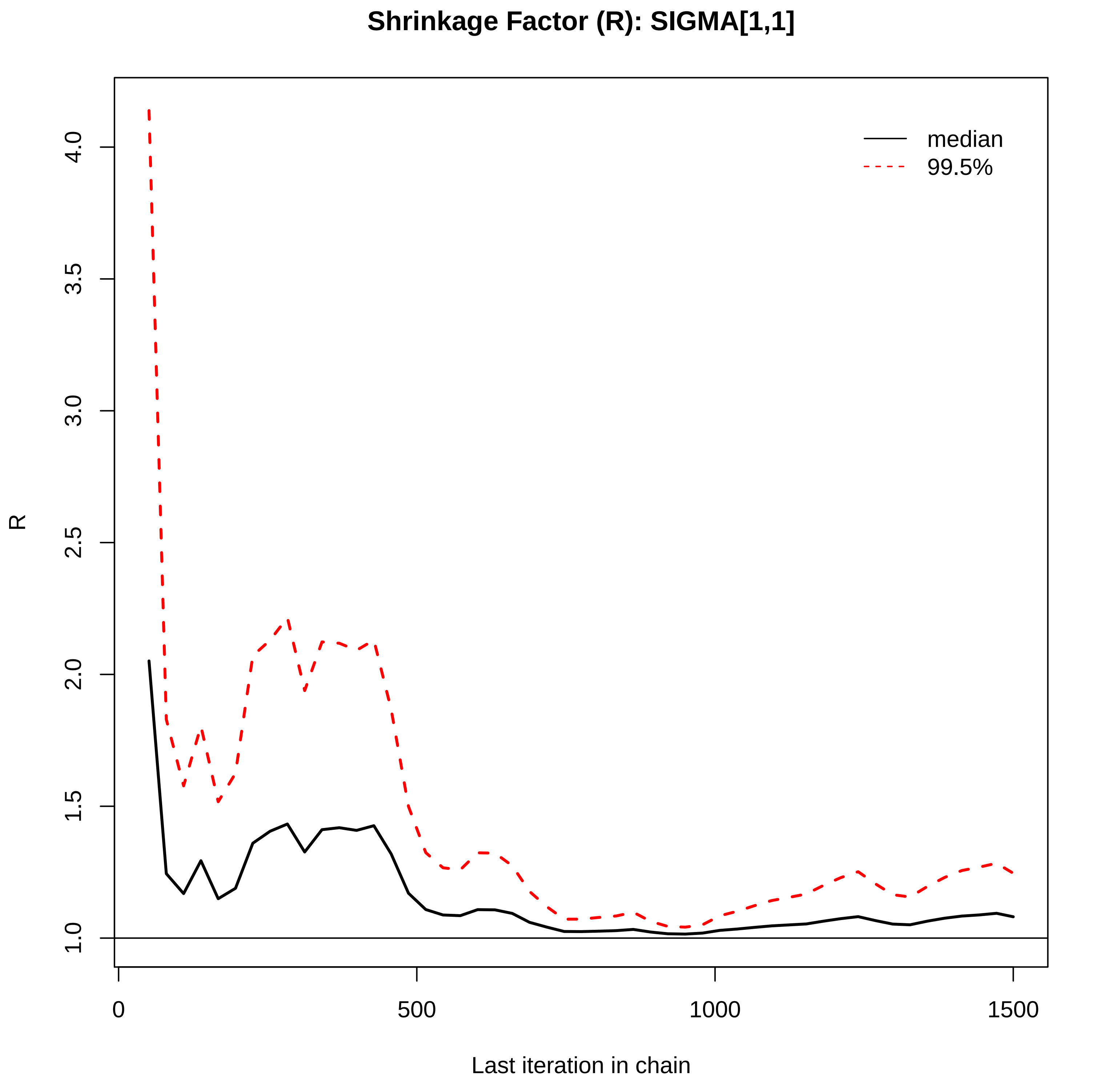
Shrinkage Factor (R): SIGMA[1,1]
median
99.5%
R
4.0
3.5
3.0
2.5
2.0
1.5
1.0
0
500
1000
1500
Last iteration in chain

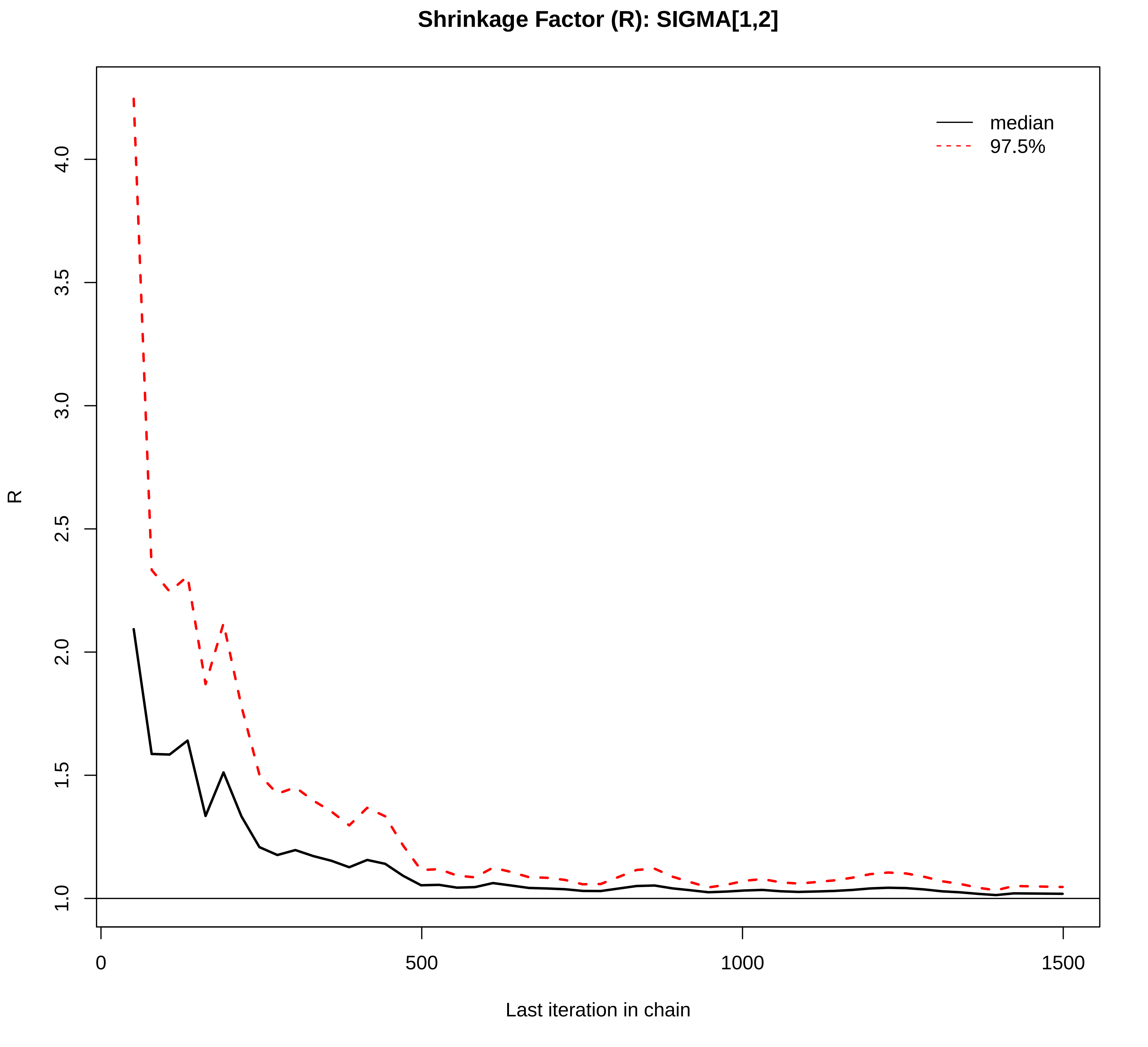
Shrinkage Factor (R): SIGMA[1,2]
median
97.5%
R
1.0
1.5
2.0
2.5
3.0
3.5
4.0
0
500
1000
1500
Last iteration in chain

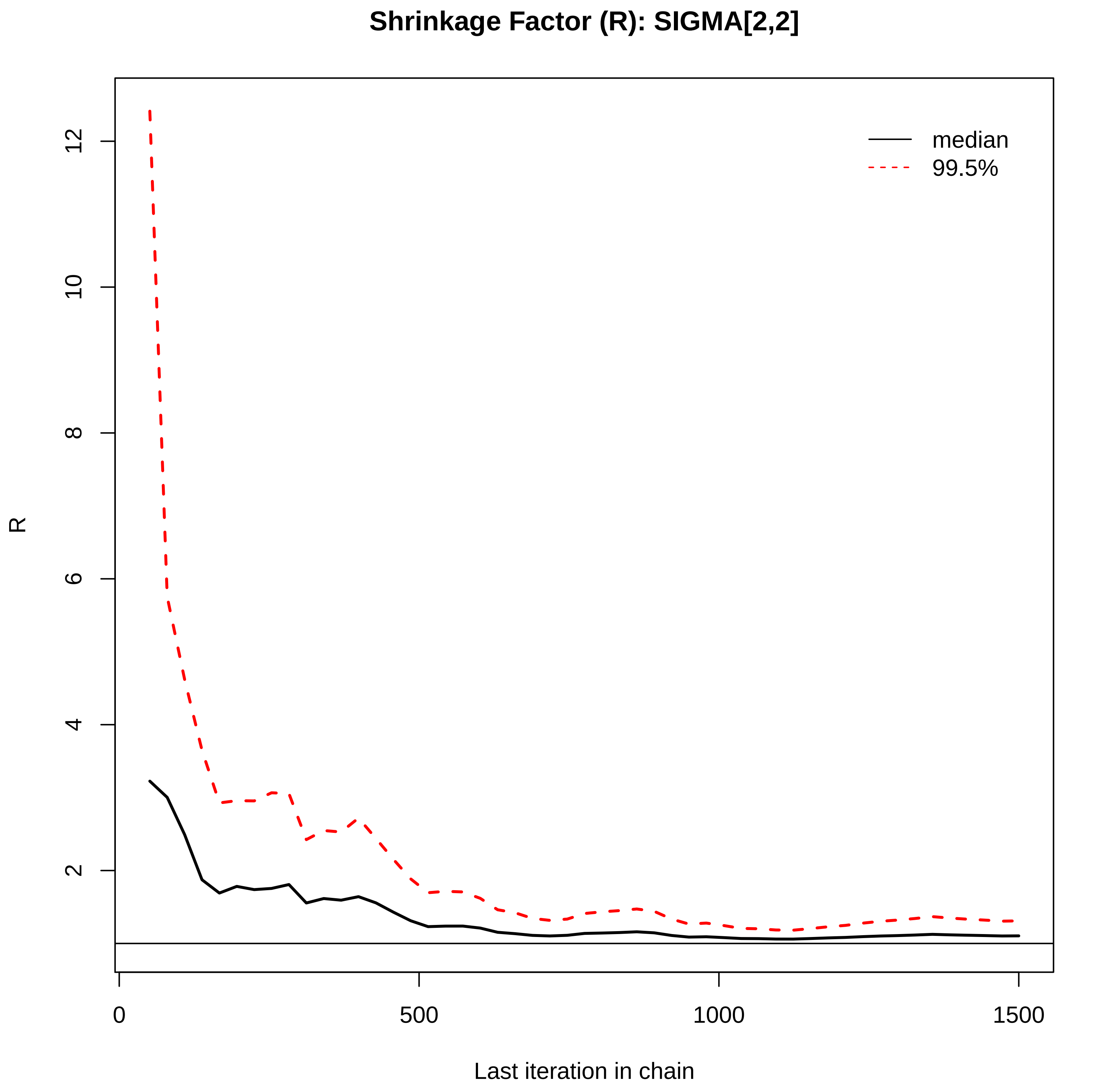
Shrinkage Factor (R): SIGMA[2,2]
median
99.5%
R
12
10
8
6
4
2
0
500
1000
1500
Last iteration in chain

**Trace plot of R011 post−burn (2000 samples in each of chains 1−6)**
**1:Black 2:Red 3:Blue**
**4:Green 5:Magenta 6:Gray**

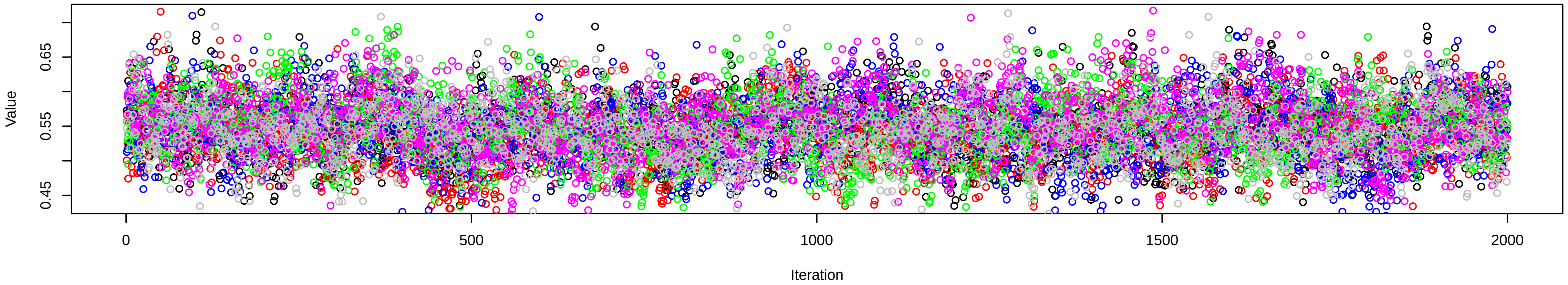


**Trace plot of R012 post−burn (2000 samples in each of chains 1−6)**
**1:Black 2:Red 3:Blue**
**4:Green 5:Magenta 6:Gray**

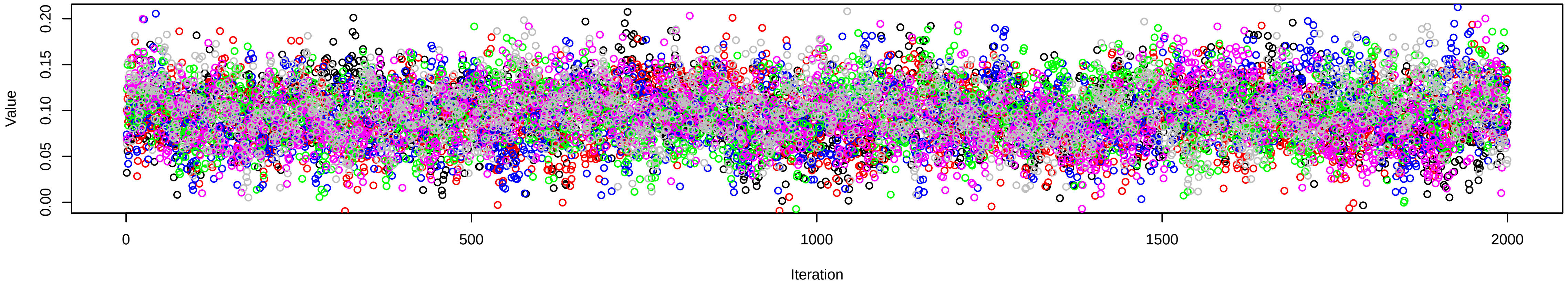


**Trace plot of R022 post−burn (2000 samples in each of chains 1−6)**
**1:Black 2:Red 3:Blue**
**4:Green 5:Magenta 6:Gray**

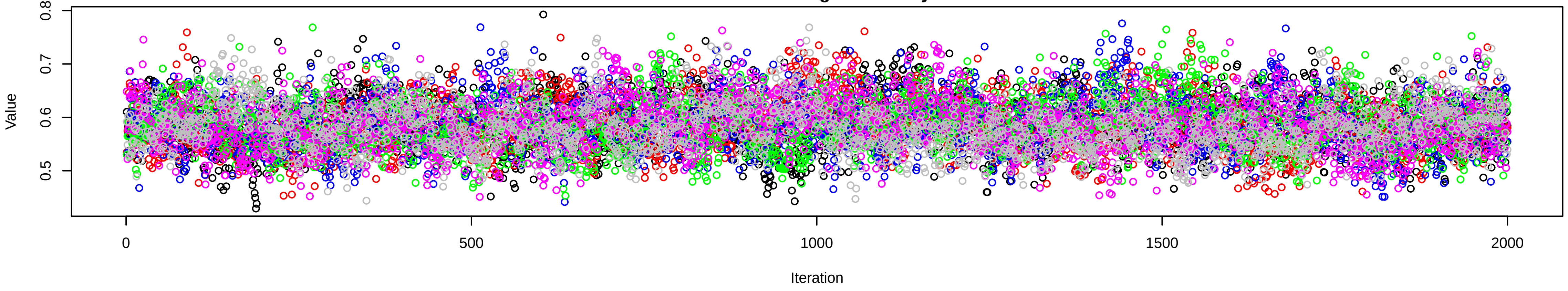

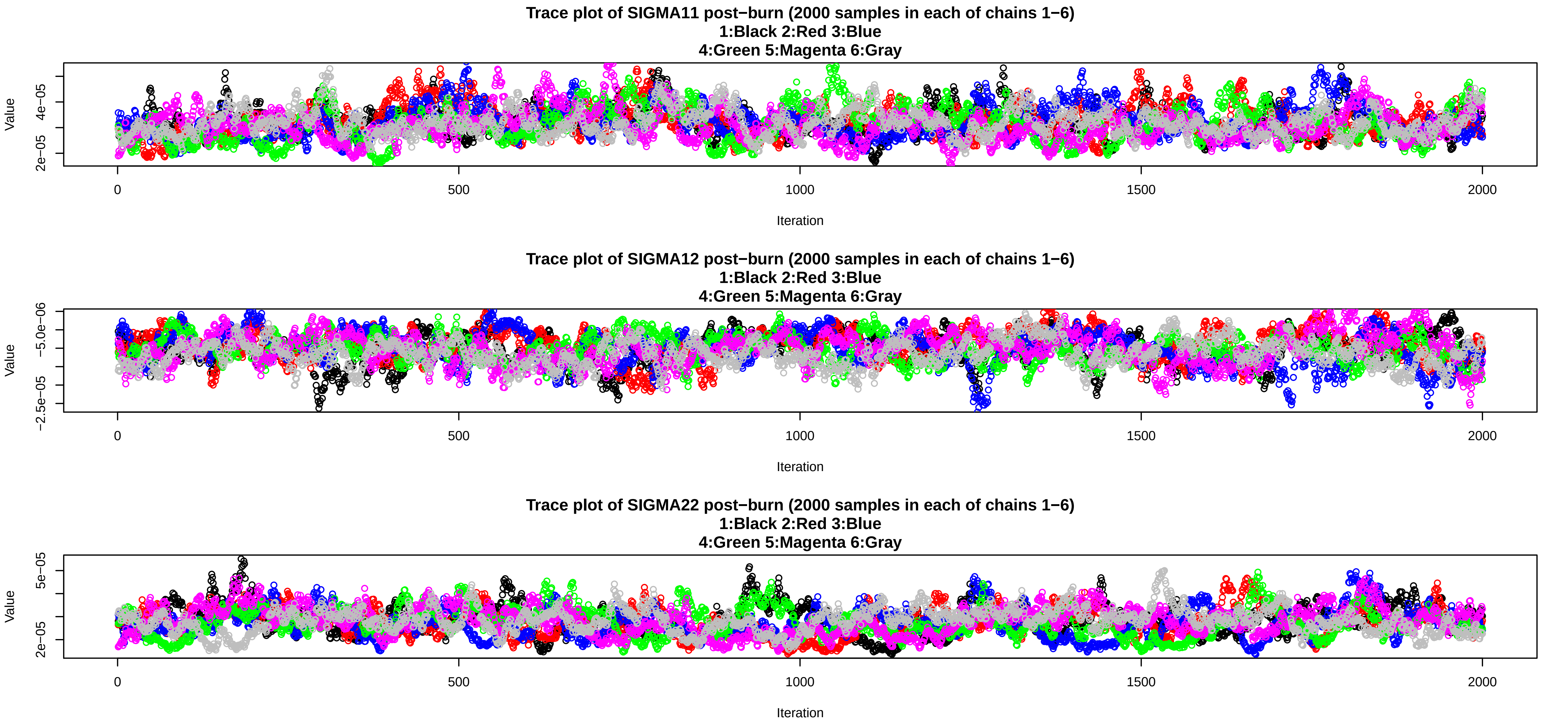
Trace plot of SIGMA11 post−burn (2000 samples in each of chains 1−6)
1:Black 2:Red 3:Blue
4:Green 5:Magenta 6:Gray
Value
2e−05
4e−05
0
500
1000
1500
2000
Iteration
Trace plot of SIGMA12 post−burn (2000 samples in each of chains 1−6)
1:Black 2:Red 3:Blue
4:Green 5:Magenta 6:Gray
Value
−2.5e−05
−5.0e−06
0
500
1000
1500
2000
Iteration
Trace plot of SIGMA22 post−burn (2000 samples in each of chains 1−6)
1:Black 2:Red 3:Blue
4:Green 5:Magenta 6:Gray
Value
2e−05
5e−05
0
500
1000
1500
2000
Iteration

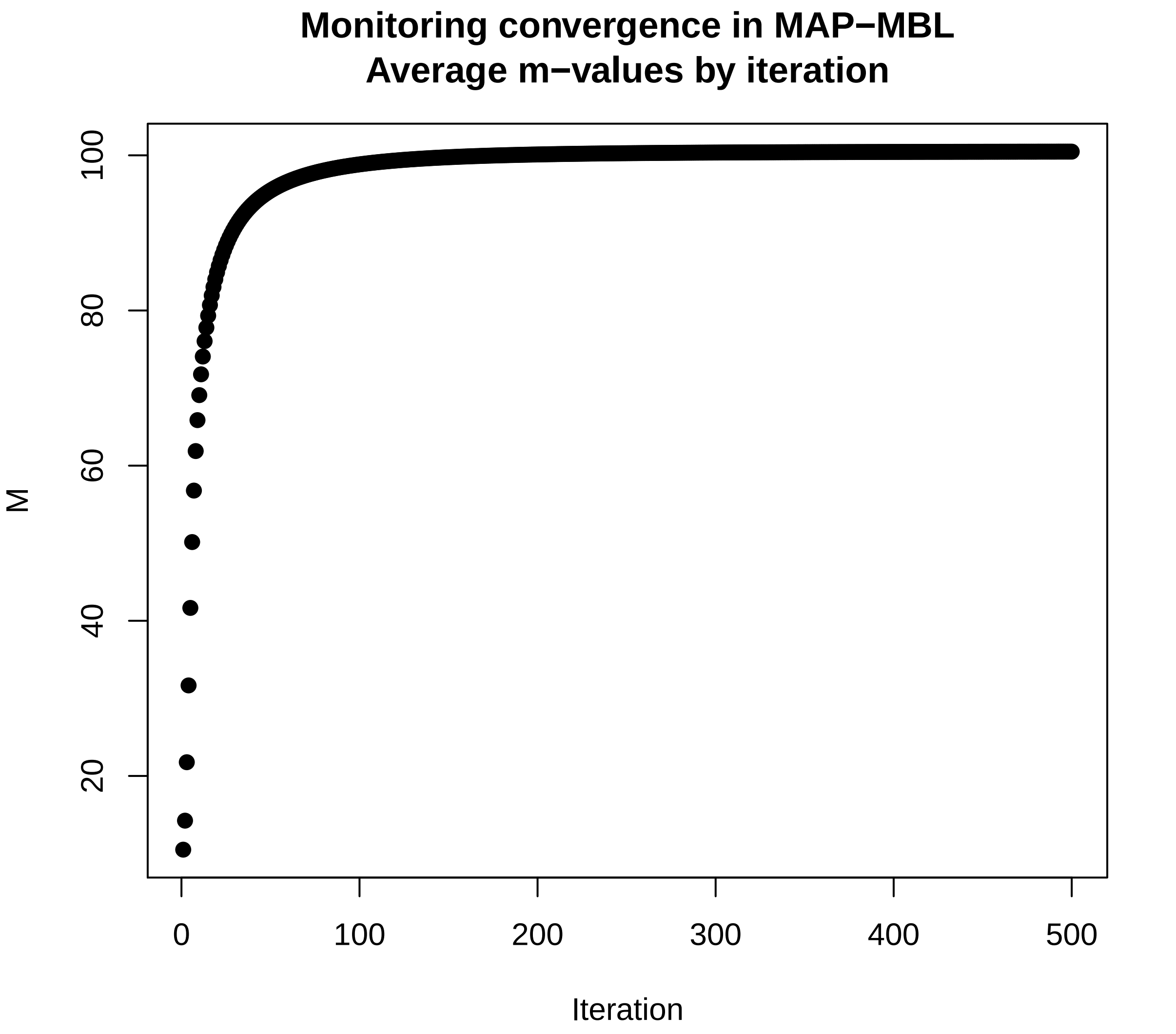
Monitoring convergence in MAP−MBL
Average m−values by iteration
M
Iteration
20
40
60
80
100
0
100
200
300
400
500

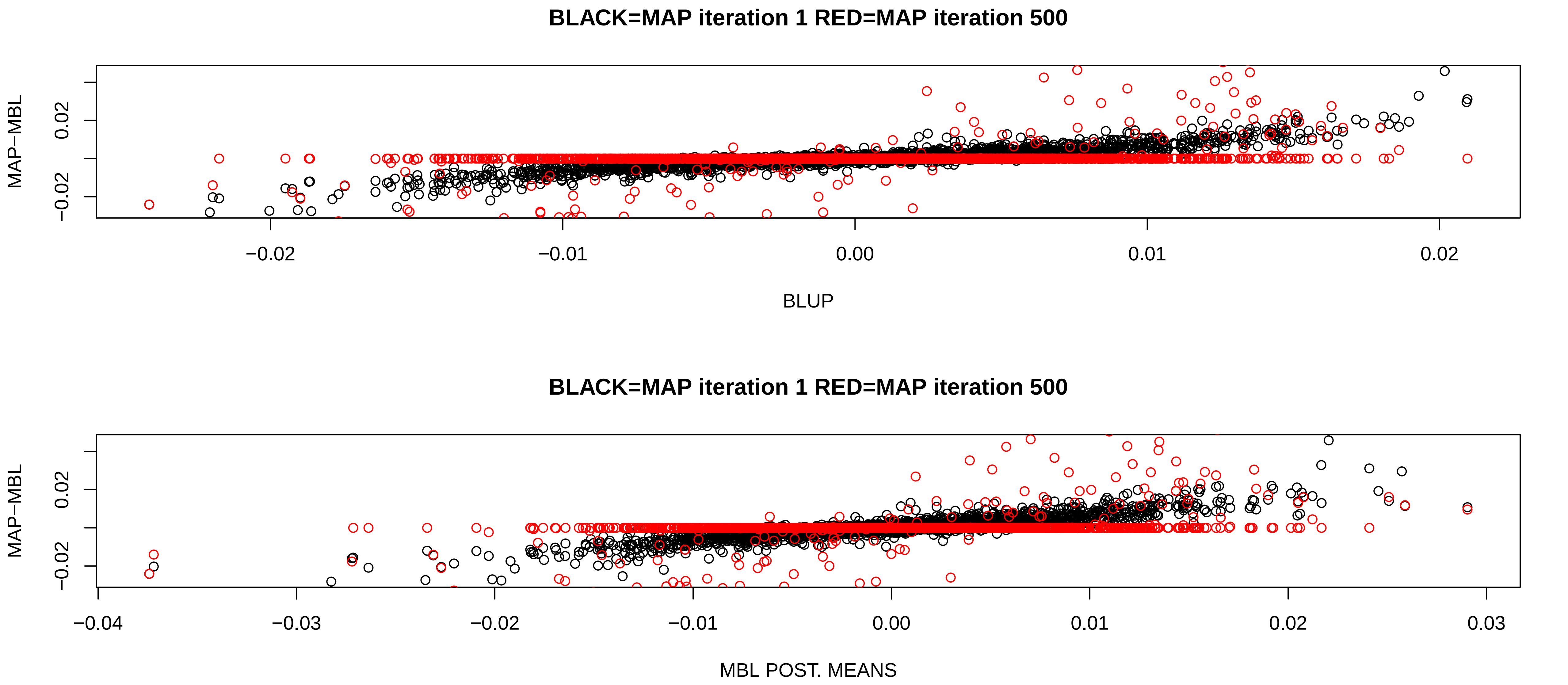
BLACK=MAP iteration 1 RED=MAP iteration 500
MAP−MBL
0.02
−0.02
−0.02
−0.01
0.00
0.01
0.02
BLUP
BLACK=MAP iteration 1 RED=MAP iteration 500
MAP−MBL
0.02
−0.02
−0.04
−0.03
−0.02
−0.01
0.00
0.01
0.02
0.03
MBL POST. MEANS

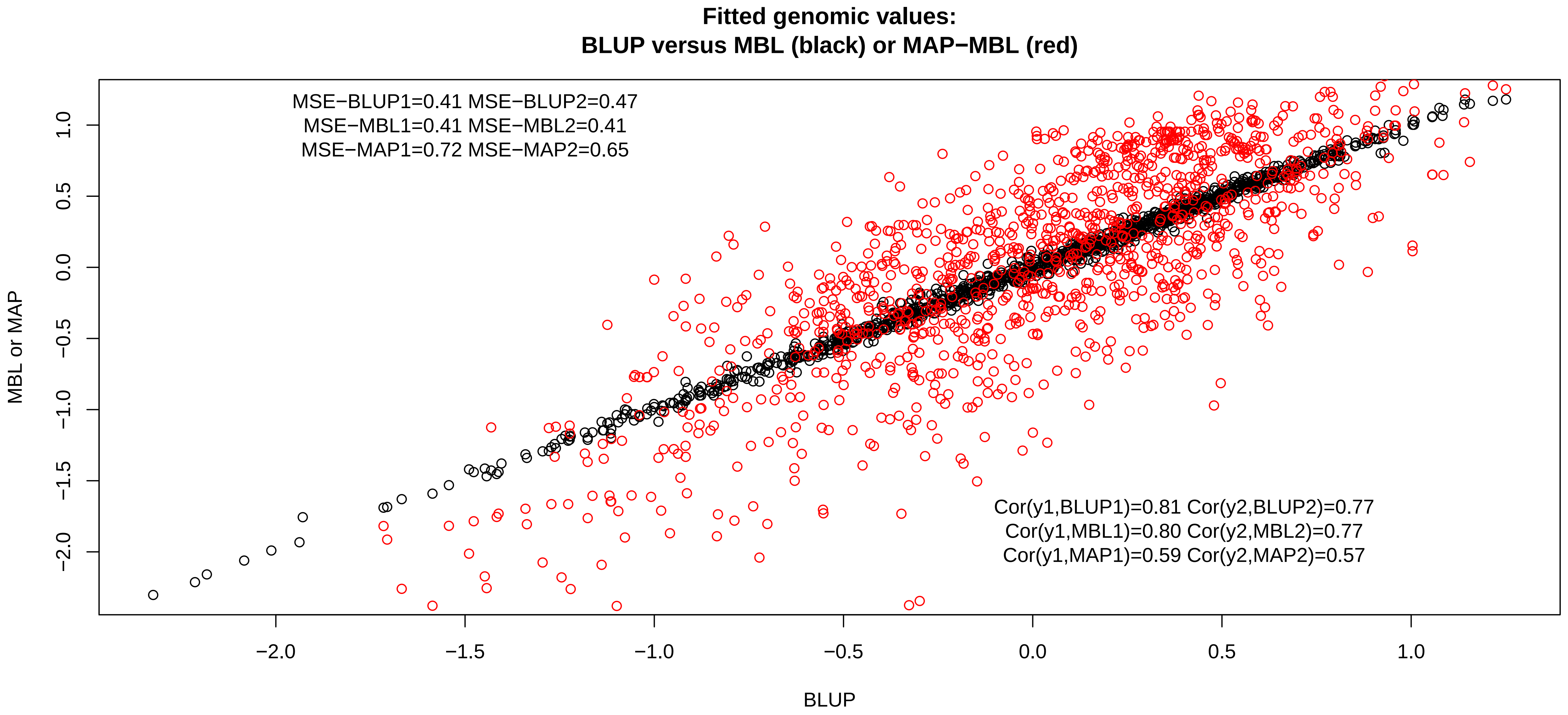
Fitted genomic values:
BLUP versus MBL (black) or MAP−MBL (red)
MSE−BLUP1=0.41 MSE−BLUP2=0.47
MSE−MBL1=0.41 MSE−MBL2=0.41
MSE−MAP1=0.72 MSE−MAP2=0.65
Cor(y1,BLUP1)=0.81 Cor(y2,BLUP2)=0.77
Cor(y1,MBL1)=0.80 Cor(y2,MBL2)=0.77
Cor(y1,MAP1)=0.59 Cor(y2,MAP2)=0.57
MBL or MAP
BLUP
1.0
0.5
0.0
−0.5
−1.0
−1.5
−2.0
